# An All-Optical Approach to Probe Chloride Transport with a Bright ChlorON

**DOI:** 10.64898/2026.03.09.710669

**Authors:** Jasmine N. Tutol, Vishaka Pathiranage, Shelby M. Phelps, Alice R. Walker, Sheel C. Dodani

**Affiliations:** Department of Chemistry and Biochemistry, The University of Texas at Dallas, Richardson, TX 75080; Department of Chemistry, Wayne State University, Detroit, MI 48202

## Abstract

Chloride transport across cellular membranes is fundamental to physiology. Yet, this dynamic process remains difficult to capture with existing methods that rely on electrophysiology or indirect iodide-quenching assays, leaving real-time imaging of chloride transport a largely unexplored frontier. To address this gap, we upgrade our first-generation fluorescent protein indicator ChlorON-1 into ChlorON-1-PRO through targeted mutagenesis of an evolutionarily conserved gatepost residue. A single mutation (C139N) preserves the turn-on sensing mechanism (13.9-fold response) while boosting affinity (*K*_d_ = 47.4 mM) and bound-state brightness (13.6). Molecular dynamics simulations provide atomic-level insights for these enhancements, supporting a model in which the mutation globally rigidifies the β-barrel and locally prearranges the binding pocket while stabilizing the chromophore. Finally, we showcase the utility of ChlorON-1-PRO for real-time monitoring of endogenous chloride transport under basal and pharmacologically modulated conditions in the U-2 OS cell model.

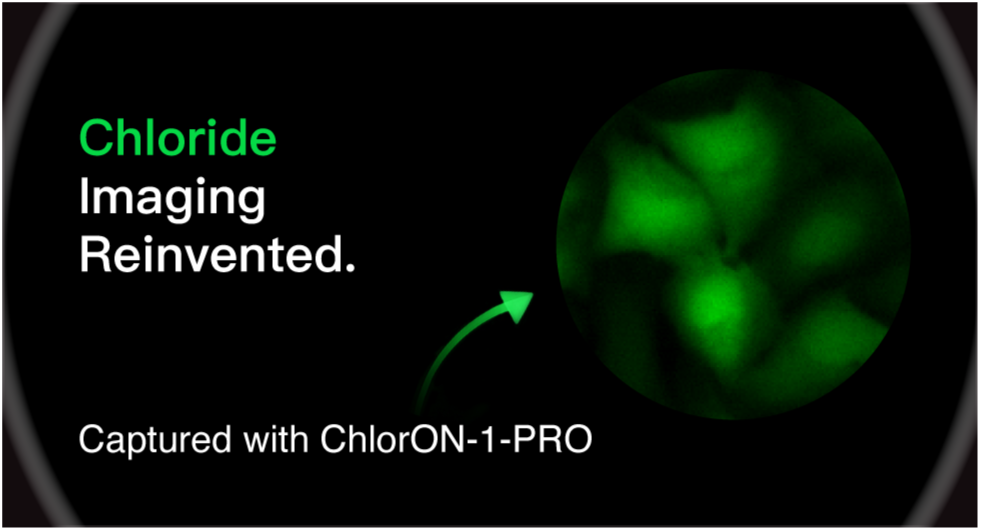

## Introduction

Chloride is the most prevalent inorganic anion in the human body.^1–3^ Across tissues and organs, more than fifty chloride-transporting proteins, spanning channels, exchangers, antiporters, and symporters, mobilize chloride at plasma and organellar membranes.^4–9^ Their activity is regulated by a myriad of physiological cues including accessory proteins, ions, ligands, pH, post-translational modifications, redox state, and voltage.^10–18^ Depending on the cellular context, this diversity gives rise to distinct chloride gradients that can be dynamically tuned over the low-to high-millimolar range, allowing chloride not only to balance charge but also to act as a signal that modulates enzyme activity, regulates protein conformation, and influences gene expression.^19–21^ Moreover, disrupted chloride transport activity underlies a wide range of disease phenotypes, motivating growing efforts toward pharmacological intervention.^7^

Our understanding of chloride-transporting proteins has been shaped by decades of genetic, structural, and biochemical studies.^5,6,22–25^ At the cellular level, chloride transport has traditionally been interrogated with electrophysiology.^26–29^ To extend this across larger populations of cells, *Aequorea victoria* green fluorescent protein (GFP)-derived indicators emerged as optical complements nearly three decades ago.^30–33^ Leveraging the greater tunability of GFP, a diverse lineage of fluorescent protein indicators has since been developed for a range of biological applications.^34–36^ Among these, the pioneering variant YFP-H148Q has stood the test of time.^30^

In YFP-H148Q, halides and nitrate permeate a water-filled channel between the β7 and β10 strands to the anion binding pocket adjacent to the chromophore (Figure 1A).^37^ This shifts the chromophore equilibrium from the fluorescent phenolate form to the non-fluorescent phenol form.^37,38^ Given its preferential affinity for iodide over chloride, YFP-H148Q is commonly used to indirectly report on chloride in cells via anion exchange with iodide.^38,39^ Beyond fluorescence imaging, high-throughput screening with YFP-H148Q and its variants has enabled the discovery of small-molecule drugs and chloride-transporting proteins.^40–43^

**Figure 1.**
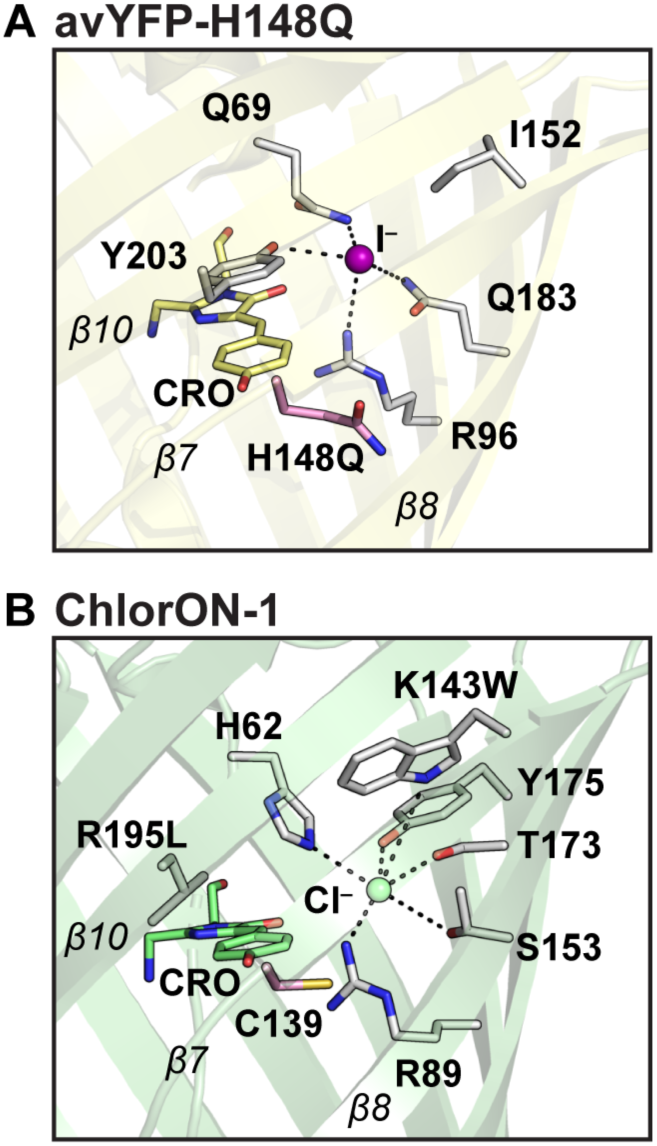
Structural comparison of YFP-H148Q and ChlorON-1 to identify a mutational hotspot in the gatepost region. (A) Anion binding pocket of YFP-H148Q with bound iodide (PDB ID: 1F09). (B) Anion binding pocket of ChlorON-1 with chloride, modeled by structural alignment to mNeonGreen (PDB ID: 5LTP). All residues within 4 Å of the anion are shown with coordinating interactions depicted as dashed lines. Residue C139 in the gatepost region, analogous to H148 in YFP, was targeted for mutagenesis (pink sticks). Key β-strands are labeled. Abbreviation: CRO, chromophore.

Despite their impact, GFP-derived indicators undergo fluorescence quenching upon anion binding, which can reduce signal contrast and complicate spatial and temporal interpretation of anion dynamics.^35,36^ To access new sensing features and biological insights, recent efforts have explored alternative fluorescent protein scaffolds and engineering strategies.^44–51^ In 2024, we introduced the ChlorON family of indicators, which provide a direct readout of chloride at physiological pH through a turn-on sensing mechanism.^52^ This unprecedented function was unlocked in the bright GFP mNeonGreen through targeted mutagenesis within the anion binding pocket at K143 and R195, albeit at the cost of overall brightness – a critical imaging parameter (Figure 1B).^52^

Motivated by the goal to overcome this limitation, we examined the microenvironment around the chromophore and focused on position 139, a conserved gatepost residue along the β7 strand of the β-barrel that is structurally analogous to position 148 in the GFP-derived indicators (Figure 1).^30,53^ The H148Q mutation in YFP increases anion accessibility and strengthens the coupling between binding and chromophore protonation, resulting in greater anion sensitivity (Figure S1).^37^ Here, we reasoned that mutagenesis at C139 could enhance chloride-dependent optical responses in the ChlorON scaffold while preserving the unique turn-on sensing mechanism for imaging cellular chloride.

## Results and Discussion

### Engineering ChlorON-1

To explore the role of position 139 in tuning ChlorON function, a library encoding all possible amino acid substitutions was generated by site-saturation mutagenesis using the 22c-trick (Figure S2, Table S1).^52,54^ Among the ChlorON indicators, ChlorON-1 was chosen because it has the largest dynamic range but weakest affinity for chloride and lowest overall brightness, providing clear opportunities for further optimization.^52^ The resulting library of C139 variants was transformed into *E. coli* and 88 colonies (>95% theoretical coverage) were randomly selected for expression and screening in a 96-well plate format (Figure S3).^54^

Since ChlorON-1 was originally identified based on its response to bromide in *E. coli* lysate at physiological pH, we extended our anion-walking strategy to chloride.^52^ From 100 mM bromide at pH 8, the selection pressure was shifted to 25 mM chloride at pH 7.4 to favor the identification of variants that maintain a large dynamic range with higher affinity at physiological pH.^52^ Variants were screened in the absence (F_i_) and presence (F_f_) of chloride and evaluated based on the turn-on response (F_f_/F_i_) alongside the overall emission intensity relative to the parent in the bound form.

Using these criteria, the top two-performing variants were identified and re-screened in *E. coli* lysate at pH 7.0 to confirm their response to 100 mM chloride and determine apparent dissociation constants (*K*_d_). Sequencing identified two distinct mutations: C139N and C139H. Relative to ChlorON-1 (F_f_/F_i_ = 8.2 ± 2.8; *K*_d_ = 205 ± 34 mM (average ± standard deviation)), the C139N variant (F_f_/F_i_ = 15.8 ± 3.1; *K*_d_ = 31.5 ± 1.2 mM) outperformed the C139H variant (F_f_/F_i_ = 3.8 ± 0.4; *K*_d_ = 4.7 ± 0.4 mM) across both metrics (Figure S4, Table S2). Based on the larger dynamic range, the C139N variant, hereafter referred to as ChlorON-1-PRO, was selected for further characterization.

### Spectroscopic characterization of ChlorON-1-PRO

For spectroscopic characterization, ChlorON-1-PRO was recombinantly expressed in *E. coli* with an N-terminal poly-histidine tag and purified via affinity and size-exclusion chromatography as a monomer (Figure S2, Table S2, Figure S5–S6). At pH 7.0, apo ChlorON-1-PRO formulated with 1 mM chloride (F_i_) displayed an absorption maximum centered at 490 nm (ε = 26,300 ± 400 M⁻¹ cm⁻¹) that red-shifted to 495 nm (ε = 36,800 ± 900 M^−1^ cm^−1^) upon the addition of 200 mM chloride (F_f_) (Figure 2A, Figure S7–S8, Table 1). This ground-state behavior is consistent with vibrational tuning of the phenolate form of the chromophore.^55^ Excitation at 485 nm produced a weak emission at 512 nm (Φ = 0.03 ± 0.01) (Figure 2B, Figure S7–S8, Table 1). The addition of chloride did not shift this maximum but triggered a ∼14-fold increase (Φ = 0.37 ± 0.01), with an apparent dissociation constant of 47.4 ± 7.4 mM (Figure 2B, Figure S7–S8, Table 1).

**Figure 2.**
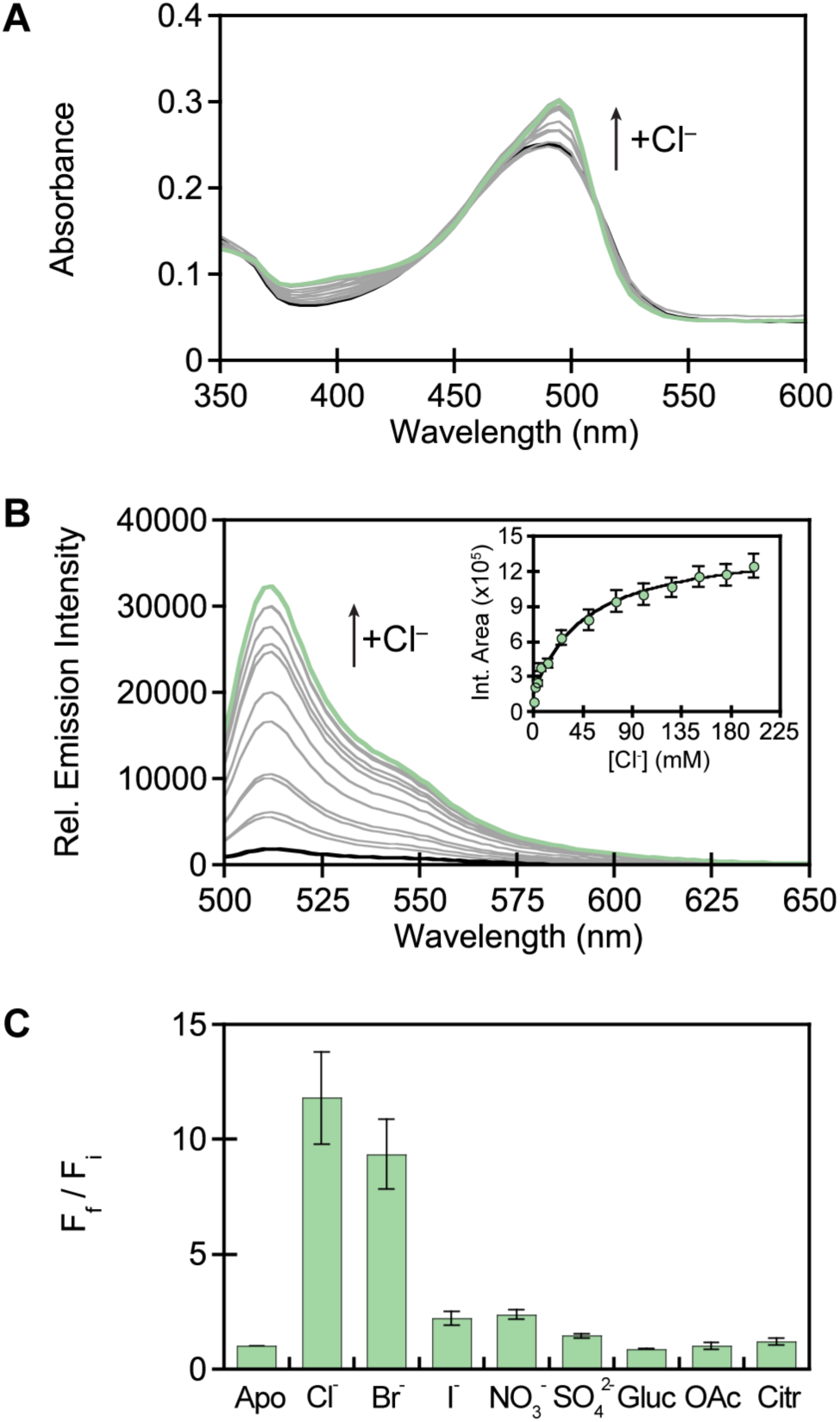
Spectroscopic characterization of ChlorON-1-PRO. (A) Absorbance and (B) emission spectra (λ_ex_ = 485 nm) of ChlorON-1-PRO with 1 (black), 2, 4.1, 7.3, 13.5, 26, 51, 76, 101, 126, 151, 176 (gray), and 201 (green) mM chloride. Inset: Integrated emission response from 500–650 nm. (C) The integrated emission response (F_f_/F_i_) of ChlorON-1-PRO from 500–650 nm in the presence of 1 mM chloride (F_i_), 201 mM chloride (Cl^-^), and 200 mM bromide (Br^-^), iodide (I^-^), nitrate (NO_3_^-^), sulfate (SO_4_^2-^), gluconate (Gluc), acetate (OAc), or citrate (Citr) (F_f_). All experiments were carried out with ∼10 µM protein in 50 mM sodium phosphate buffer at pH 7. The sodium counterion was used for all anions. The average of four technical replicates across two protein batches is shown. For clarity, the standard deviations for panels A and B are shown in Figure S8.

**Table 1.** Summary of the spectroscopic properties for ChlorON-1 and ChlorON-1-PRO at pH 7. The fluorescence response (F_f_/F_i_) in the presence of 1 mM NaCl (F_i_, -Cl^-^) and 197 mM NaCl for ChlorON-1 or 201 mM NaCl (F_f_, +Cl^-^) for ChlorON-1-PRO, the apparent dissociation constants (*K*_d_), and chromophore p*K*_a_ values are shown. Values are reported as the average ± standard deviation of four technical replicates across two protein batches.

|  |  | **ChlorON-1** | **ChlorON-1-PRO** |
| --- | --- | --- | --- |
| **Response ($F_f/F_i$)** | | $45 \pm 3$ | $13.9 \pm 2.6$ |
| **Apparent dissociation constant ($K_d$, mM)** | | $285 \pm 59$ | $47.4 \pm 7.4$ |
| **Chromophore $pK_a$** | $pK_a -Cl^-$ | N.D.<sup>a</sup> | N.D.<sup>a</sup> |
| | $pK_a +Cl^-$ | $6.0 \pm 0.1$ | $6.3 \pm 0.4$ |
| **Absorption and Emission Maximum (nm)** | $-Cl^-$ | 480, 515 | 490, 512 |
| | $+Cl^-$ | 480, 515 | 495, 512 |
| **Extinction coefficient ($\times 10^3 (M^{-1} * cm^{-1})$)** | $\epsilon -Cl^-$ | $27.4 \pm 4^b$ | $26.3 \pm 0.4^c$ |
| | $\epsilon +Cl^-$ | $24.2 \pm 3^b$ | $36.8 \pm 0.9^d$ |
| **Quantum yield** | $\Phi -Cl^-$ | $< 0.01$ | $0.03 \pm 0.01$ |
| | $\Phi +Cl^-$ | $0.027 \pm 0.007$ | $0.37 \pm 0.01$ |
| **Molar brightness** | $-Cl^-$ | $0.01^b$ | $0.08^c$ |
| | $+Cl^-$ | $0.65^b$ | $13.6^d$ |

<sup>a</sup> Abbreviation: N.D., not determined.
<sup>b</sup> Determined with the ChlorON-1 absorbance at 480 nm.
<sup>c</sup> Determined with the ChlorON-1-PRO absorbance at 490 nm.
<sup>d</sup> Determined with the ChlorON-1-PRO absorbance at 494 nm.

ChlorON-1-PRO remains responsive to chloride over a broad pH range, with the largest variations in signal observed near and below the chromophore p*K*_a_ (Figure S9). Based on these data, the chromophore p*K*_a_ in the presence of chloride was determined to be 6.3 ± 0.4, whereas the apo form could not be determined due to low signal intensity (Table 1, Figure S9). Further evaluation of the anion sensitivity revealed that ChlorON-1-PRO has a comparable response to bromide (F_f_/F_i_ = 11.6 ± 2.8; *K*_d_ = 41.3 ± 2.2 mM) but shows little to no response to iodide or the other oxyanions nitrate, sulfate, gluconate, acetate, citrate, and phosphate (F_f_/F_i_ < 2.5; *K*_d_ = not determined) (Figure 2C, Figure S10–S11, Table S3). Overall, the major spectral features of the parent ChlorON-1 are preserved in ChlorON-1-PRO, and no significant change in either the p*K*_a_ or anion preference is observed (Table 1, Figure S12).^52^ However, the C139N mutation reduced the dynamic range for chloride by 70% but increased the affinity by ∼6-fold and bound-state brightness by ∼21-fold (Table 1).^52^

### In silico analysis of ChlorON-1-PRO

To investigate how the C139N mutation enhances chloride sensing, molecular dynamics (MD) simulations were performed for ChlorON-1 and ChlorON-1-PRO in both apo and chloride-bound states at pH 7. In general, ChlorON-1-PRO had a more rigid protein backbone in the apo form as compared to ChlorON-1. Quantitative comparison of per-residue root-mean-square fluctuation (RMSF) differences showed that flexibility changes between the apo and bound states were substantially lower in ChlorON-1-PRO, particularly for the β7 and β10 strands, the likely entry and exit point of ions as shown in previous work (Figure 3).^55^ The increase in rigidity and reduced RMSF difference between the bound and apo states correlated with the observed decrease in dynamic range. Upon chloride binding, ChlorON-1 underwent a larger change in overall protein motion and had a more flexible chromophore, whereas ChlorON-1-PRO had a smaller change in overall protein motion and a more rigid chromophore with more substantial movement localized to residues within the binding pocket in the presence of chloride (Figure 3). In line with this, dynamic cross-correlation (DCC) and normal-mode analyses revealed more positively correlated motions, albeit subtly, and reduced breathing motions, particularly near the chromophore (Figure S13–S14). Similarly, the water volume within the barrel showed a small difference between apo and bound states equivalent to about one water for ChlorON-1-PRO but two to three waters for ChlorON-1 (Figure S15). Overall, ChlorON-1-PRO adopts a more structurally rigid framework that supports a molecular model for its altered affinity, brightness, and sensing performance.

**Figure 3.**
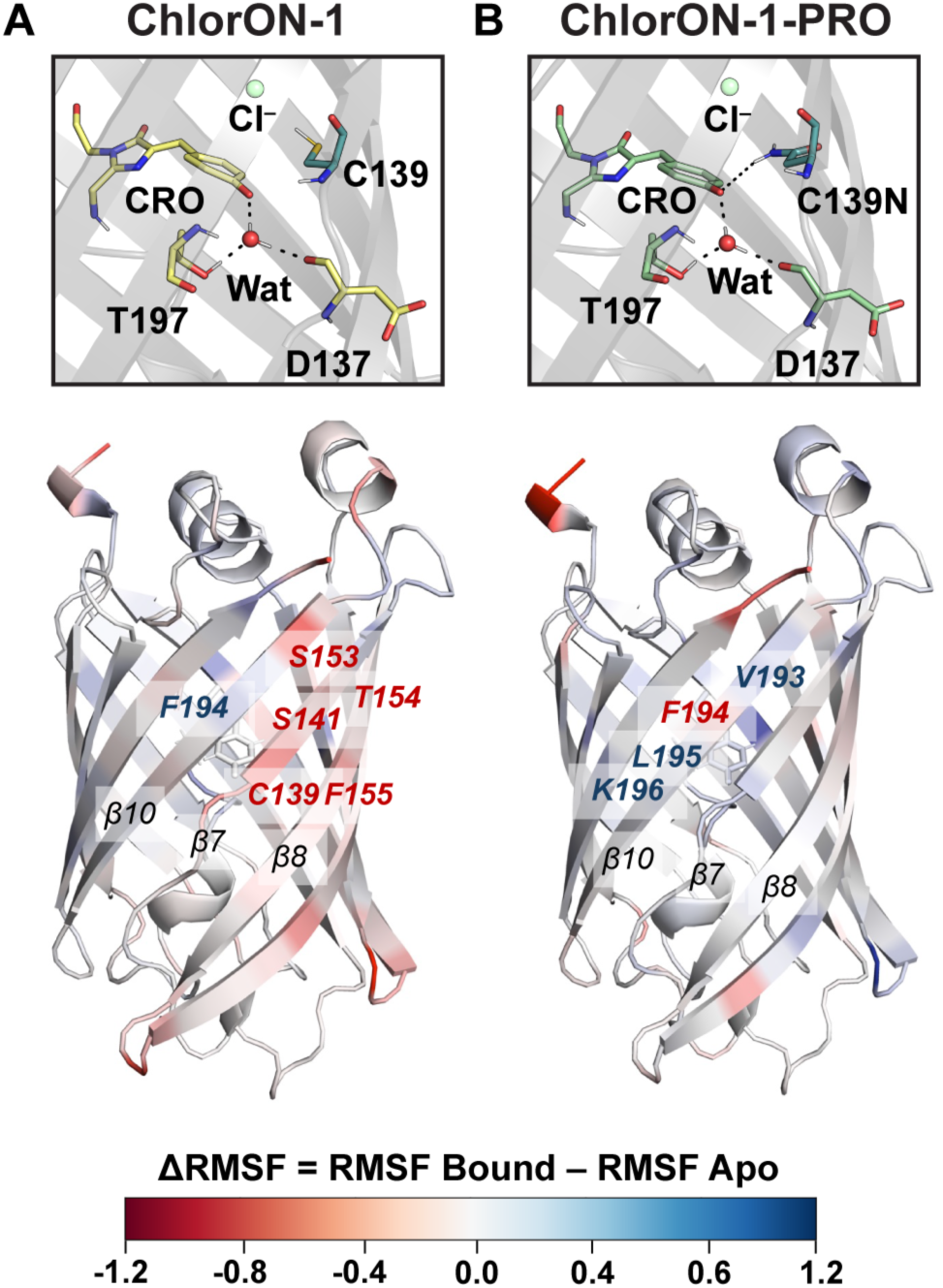
Molecular dynamics (MD) simulations reveal differences between (A) ChlorON-1 and (B) ChlorON-1-PRO. (Top) Representative snapshots of the hydrogen-bonding interactions around the chromophore. (Bottom) Differences in the RMSF between the apo and chloride-bound states with increased (blue) or decreased (red) flexibility in the bound state relative to the apo state. Abbreviations: CRO, chromophore; RMSF, root-mean-square fluctuation.

In the chloride-bound states, as expected, the chloride binding pocket is positioned adjacent to the chromophore (Figure S16) where position 139 lies along the β7 strand, with its side chain facing inward. In both ChlorON-1 and ChlorON-1-PRO, the side chain of H62 rotates to directly coordinate chloride, indicating its key role in anion recognition. More pronounced rearrangements of the S141 and W143 side chains are observed in ChlorON-1-PRO than ChlorON-1 upon chloride binding, which coincides with a greater twisting of the phenolate ring relative to the imidazolinone core of the chromophore (Figure S16). Thus, ChlorON-1-PRO is generally more rigid, but the conformational changes we observe are more specific to the sensing mechanism and motion of the chromophore. This conformation mimics the bound state observed in ChlorON-3 in our previous QM/MM and spectroscopic work on the ChlorON series, where chloride binding was shown to tilt and lock the chromophore into a highly emissive and rigidified position.^55,56^

In the bound state, chromophore stabilization in ChlorON-1 appears to rely on a front-facing water molecule that interacts with D137 (37% occupancy) and T197 (51% occupancy). These interactions are less prevalent in ChlorON-1-PRO (17% for D137 and 31% for T197). Akin to YFP-H148Q, ChlorON-1-PRO exhibits enhanced hydrogen bonding at the phenolate end of the chromophore, involving both a water molecule (93% occupancy) and the side chain of N139 (33% occupancy), thereby stabilizing the chromophore in a single, well-defined conformation (Figure 3, Figure S16, Table S4). This effect is connected to a narrower distribution of chloride positional shifts relative to its initial position (∼0-1.1 Å) as compared to ChlorON-1 (∼0-1.5 Å) (Figure S17). While our classical MD simulations here do not model the excited state directly, rigidification of the chromophore in the ground state has been previously associated with improved fluorescence quantum yield.^56^ We therefore associate these alterations in the binding pocket with improved spectroscopic features. Together, these data support a multifactorial model in which distributed changes in protein rigidity, binding-pocket organization, and chromophore stabilization collectively contribute to the sensing properties of ChlorON-1-PRO.

### Live-cell fluorescence microscopy with ChlorON-1-PRO

From these insights, ChlorON-1-PRO was applied to monitor chloride transport in living cells using fluorescence microscopy. For this, we selected the human osteosarcoma cell line U-2 OS, which endogenously expresses chloride-transporting proteins, including calcium-activated chloride channels (CaCCs) — such as anoctamin-1 (ANO1), bestrophins (BEST), and tweety (TTYH) — chloride intracellular channel proteins (CLICs), chloride channels and exchangers (CLCs), proton-activated chloride channels (PACs), sodium- and chloride-dependent GABA and glycine transporters, and volume-regulated anion channels (VRACs).^57–61^ To streamline the imaging workflow, fluorescence-activated cell sorting (FACS) was used to generate a U-2 OS cell line stably expressing ChlorON-1-PRO. When imaged in buffer containing 137 mM chloride, cells displayed bright, evenly distributed fluorescence throughout the cytosol and nucleus, consistent with the chloride-bound form of the indicator (Figure S18–S19). To confirm this, a chloride influx assay was performed in which cells were acutely incubated in buffer containing 137 mM gluconate to deplete the labile chloride pool (F_i_), followed by reperfusion with chloride-containing buffers (F_f_). At the start of the titration, little to no signal was observed above the background, but the fluorescence intensity progressively increased with higher chloride concentrations (Figure 4). ChlorON-1-PRO responded to extracellular chloride concentrations as low as 5 mM (F_f_/F_i_ = 2.4 ± 0.5, *p*-value < 0.05) and up to 137 mM (F_f_/F_i_ = 6.9 ± 1.4, *p*-value < 0.05) (Figure 4). Under matched experimental conditions, the parent ChlorON-1 responses were not statistically distinguishable from 0 mM chloride across the concentrations tested (*p*-value > 0.05) (Figure 4). No significant change was observed in cells expressing mNeonGreen (F_f_/F_i_ = 1.1 ± 0.3, *p*-value > 0.05) (Figure S20). The observed cell-to-cell variability could arise from indicator expression and cell morphology, as well as heterogeneity in labile chloride and its handling. The time-resolved imaging workflow captures these differences across many individual cells (600−1200 regions of interest per biological replicate) throughout the assay, with each cell serving as its own reference.

**Figure 4.**
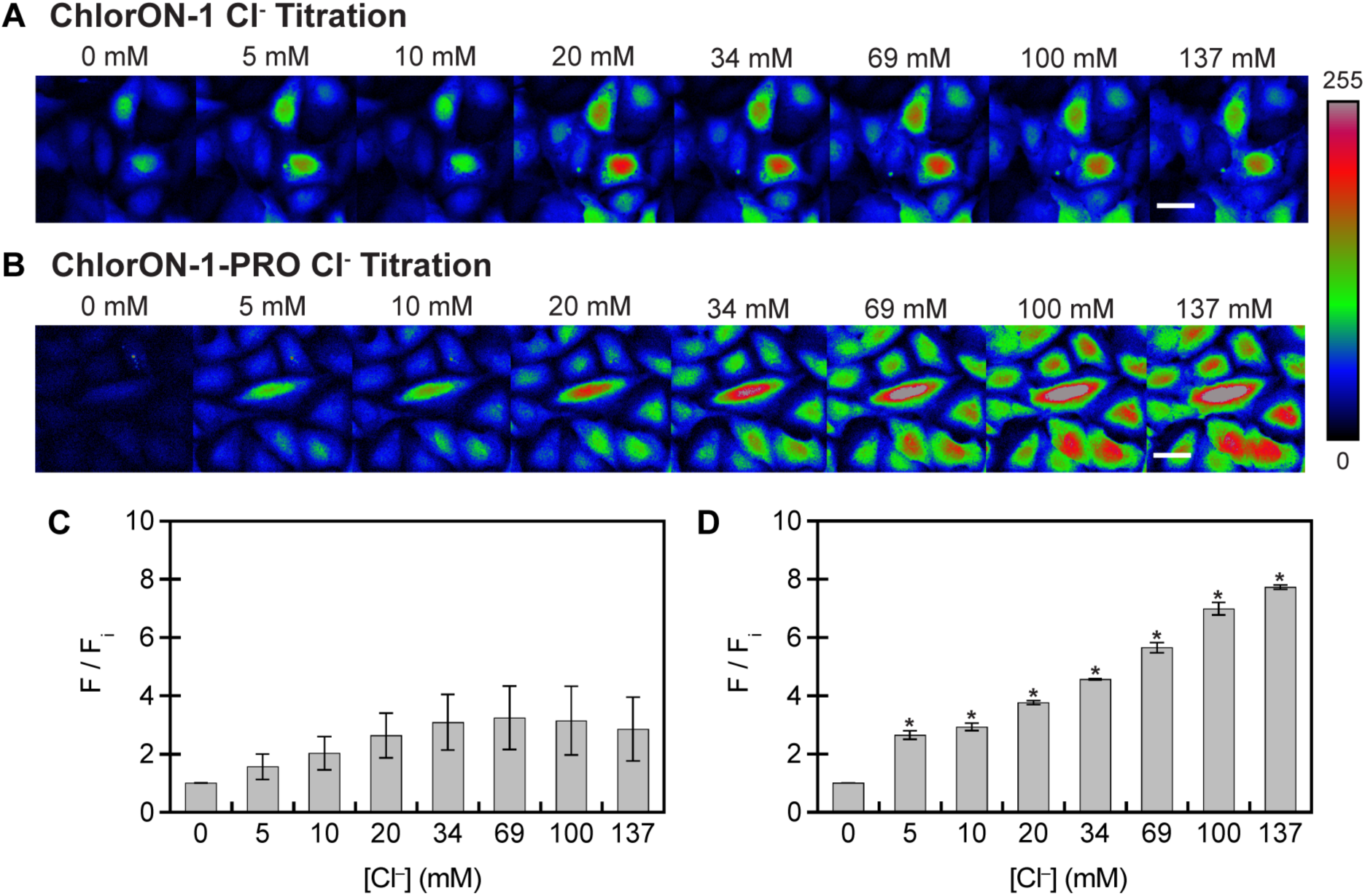
In-cell chloride titration with ChlorON-1 and ChlorON-1-PRO. Representative fluorescence images of (A) ChlorON-1 and (B) ChlorON-1-PRO expressed in U-2 OS cells in 0 mM (F_i_) and following exposure to 5, 10, 20, 34, 69, 100, and 137 mM (F) chloride. Scale bar = 30 µm. Bar graphs showing the turn-on fluorescence response (F/F_i_) for (C) ChlorON-1 (*N* = 2) and (D) ChlorON-1-PRO (*N* = 3) at each chloride concentration. Statistical comparisons at each chloride concentration were made relative to F_i_ for ChlorON-1 (*p*-value > 0.05) and ChlorON-1-PRO (* = *p*-value < 0.05). Values are reported as the average ± standard deviation of three fields from each biological replicate.

Given these robust properties, ChlorON-1-PRO was used to assess pharmacological modulation of chloride transport in live cells using the chloride influx assay. For these experiments, cells treated with 137 mM gluconate (F_i_) were perfused with buffer containing 68.5 mM gluconate and 68.5 mM chloride (F_f_) under conditions optimized to maximize the fluorescence response while maintaining modulator efficacy. All treatments contained either DMSO alone (co-solvent control, ≤ 0.2% v/v) or the specified modulator prepared in DMSO. We first tested the non-selective chloride channel inhibitor indanyloxyacetic acid-94 (IAA-94), which potently inhibits CLC family members and VRACs, and to a lesser extent CaCCs.^62–65^ Relative to the DMSO control (F_f_/F_i_ = 10.5 ± 3.9), IAA-94 (100 µM) attenuated the chloride-triggered fluorescence response (F_f_/F_i_ = 2.6 ± 2.2), demonstrating that ChlorON-1-PRO reliably reports on inhibition of chloride transport under these conditions (Figure 5, Figure S21–S22).

**Figure 5.**
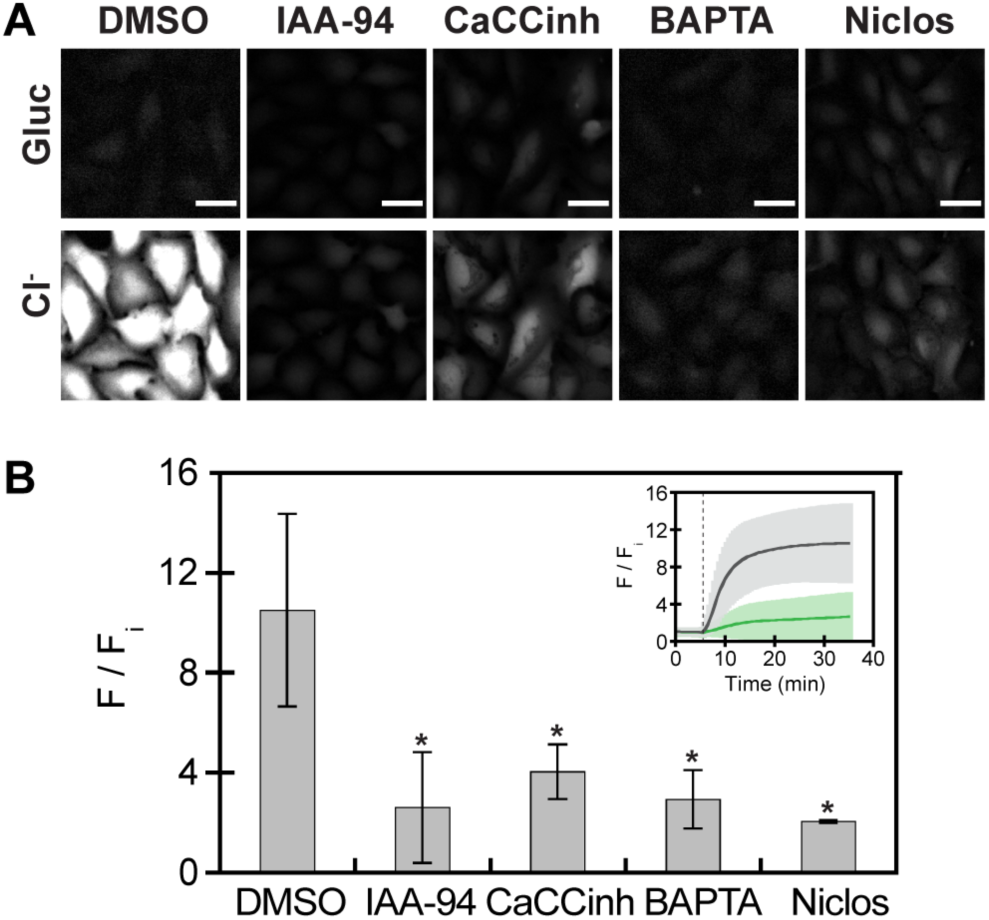
ChlorON-1-PRO provides a direct readout of chloride transport inhibition. (A) Representative fluorescence images of ChlorON-1-PRO expressed in U-2 OS cells in the presence of 137 mM gluconate (F_i_, top row) and 68.5 mM gluconate/68.5 mM chloride (F, bottom row). Cells were treated with 0.2% DMSO (vehicle control), 100 µM IAA-94, 20 µM CaCCinh-A01, 20 µM BAPTA as the AM ester, or 5 µM niclosamide. Scale bar = 30 µm. (B) Bar graph showing the turn-on fluorescence response (F/F_i_) for each treatment in panel A. Statistical comparisons for each treatment were made relative to the DMSO control (* = *p*-value < 0.05). Inset: Representative time-lapse trace of the ChlorON-1-PRO turn-on response (F/F_i_) in the presence of 137 mM gluconate (F_i_) and 68.5 mM gluconate/68.5 mM chloride (F) with DMSO (gray trace, *N* = 6) and IAA-94 (green trace, *N* = 3). The dashed line represents the start of the chloride buffer perfusion. Values are reported as the average ± standard deviation of three fields from each biological replicate. Abbreviations: AM, acetoxymethyl; CaCCinh, CaCCinh-A01; Niclos, niclosamide.

**Figure 6.**
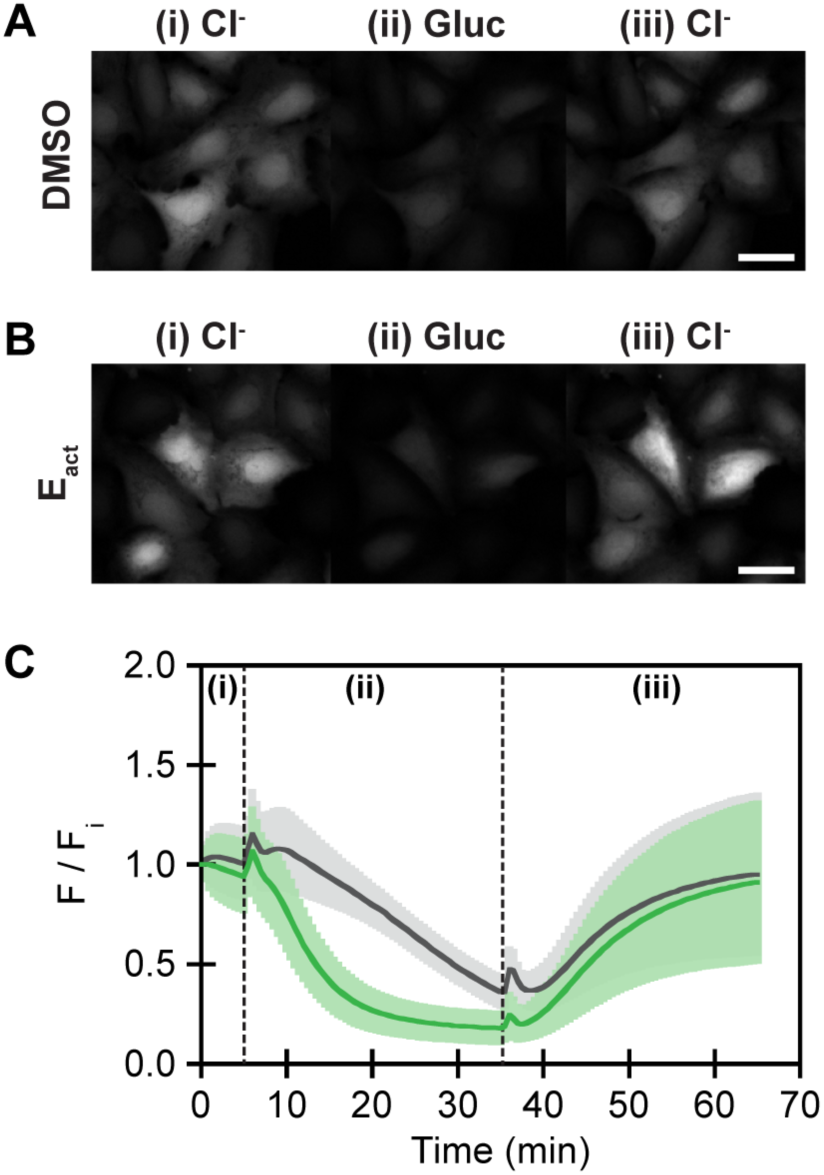
ChlorON-1-PRO provides a direct readout of chloride transport activation. Representative fluorescence images of ChlorON-1-PRO expressed in U-2 OS cells treated with (A) 0.1% DMSO (vehicle control) or (B) 10 µM E_act_ in the presence of (i) 137 mM chloride at *t =* 0 min (F_i_), (ii) 137 mM gluconate at *t =* 35 min, and (iii) 137 mM chloride at *t =* 65 min. Scale bar = 30 µm. (C) Time-lapse traces of the fluorescence response (F/F_i_) at each time point (F) relative to *t =* 0 min (F_i_) with DMSO (gray, *N* = 3) and E_act_ (green, *N* = 3). The first dashed line represents the start of the gluconate buffer perfusion, and the second dashed line represents the start of the chloride buffer perfusion. Traces are reported as the average ± standard deviation of three fields from each biological replicate.

Building from this, we focused on CaCC-mediated chloride transport, which has been well-characterized through structural, electrophysiological, and pharmacological studies.^17,66,67^ To probe CaCC-mediated transport, we selected four modulators with distinct mechanisms of action: CaCCinh-A01, the intracellular calcium chelator BAPTA delivered as the acetoxymethyl ester, niclosamide, and E_act_. CaCCinh-A01 inhibits multiple CaCCs, including ANO1 and BEST.^68,69^ For ANO1, inhibition has been attributed to binding within the ion-conduction pore near the extracellular face and longer exposure can also promote proteasomal degradation of the channel.^67,70^ Esterase-released BAPTA reduces cytosolic calcium, thereby limiting calcium-dependent gating of CaCCs and indirectly suppressing chloride transport.^12^ Niclosamide, an anthelmintic drug, is a less specific and mechanistically complex inhibitor.^71^ For ANO1, it can inhibit, and in certain cellular contexts, potentiate channel activity.^72,73^ These effects may arise through interactions with a hydrophobic pocket on the extracellular face or indirectly by altering intracellular calcium dynamics, among other mechanisms.^72–74^ E_act_ is a reported allosteric activator of ANO1 that can stimulate chloride transport independently of increases in intracellular calcium, with reported off-target activation of transient receptor potential vanilloid 1 (TRPV1).^75,76^

All three inhibitory modulators attenuated the ChlorON-1-PRO response to a comparable extent: CaCCinh-A01 (20 µM; F_f_/F_i_ = 4.1 ± 1.1), BAPTA (20 µM; F_f_/F_i_ = 2.9 ± 1.2), and niclosamide (5 µM; F_f_/F_i_ = 2.0 ± 0.1) (Figure 5, Figure S23–S25). These data point to effective, but not complete, inhibition of CaCC-mediated chloride transport, as the fluorescence signal with chloride did not reach the background level observed with gluconate. This outcome is not unexpected in cells where multiple chloride-transporting proteins at the plasma and subcellular membranes operate in parallel with varying expression levels and regulatory controls. Irrespective of potential differences in inhibitor potency, mechanism of action, or selectivity, ChlorON-1-PRO reveals a basal labile chloride pool that persists under inhibitory conditions.

In contrast, the ChlorON-1-PRO turn-on response with E_act_ was comparable to the DMSO control in the chloride influx assay used for the inhibitors (F_f_/F_i_ = 8.8 ± 2.6) (Figure S26). We hypothesized that the strong driving force for chloride influx under these conditions may limit the ability to resolve additional E_act_-mediated increases in chloride transport. The assay was therefore reconfigured to monitor chloride efflux. For these experiments, cells treated with 137 mM chloride (F_i-Cl_) were perfused with buffer containing 137 mM gluconate (F_f-Gluc_) to initiate efflux followed by reperfusion with 137 mM chloride (F_f-Cl_) to initiate influx, with either DMSO or 10 µM E_act_ present throughout. During the efflux phase, E_act_ potentiated chloride transport, resulting in a more rapid and larger decrease in the ChlorON-1-PRO fluorescence signal (F_f-Gluc_/F_i-Cl_ = 0.18 ± 0.02) relative to the DMSO control (F_f-Gluc_/F_i-Cl_ = 0.4 ± 0.1). However, during the influx phase, no difference was observed (F_f-Cl_/F_i-Cl_ DMSO = 0.9 ± 0.1; F_f-Cl_/F_i-Cl_ E_act_ = 0.9 ± 0.1). Together, these results demonstrate that ChlorON-1-PRO can report changes in endogenous chloride transport associated with channel activation, highlighting its expanded utility beyond inhibitors for monitoring chloride transport modulation.

Finally, because chloride transport can be coupled to pH, we evaluated the sensing performance of ChlorON-1-PRO across physiological pH values. To manipulate intracellular pH, cells were incubated with gluconate-to-chloride exchange assay buffers ranging from pH 6.5 to 8.0 in the presence of ionophores.^77^ In 140 mM gluconate, little to no fluorescence signal was observed, whereas reperfusion with 70 mM gluconate and 70 mM chloride produced a measurable turn-on fluorescence response that followed the trend: pH 6.5 (F_f_/F_i_ = 7.4 ± 1.3) < pH 7.0 (F_f_/F_i_ = 8.5 ± 2.0) ≈ pH 7.5 (F_f_/F_i_ = 8.9 ± 2.0) > pH 8.0 (F_f_/F_i_ = 7.9 ± 1.6) (Figure S27–S30).

To confirm the effectiveness of the pH clamping, parallel experiments were performed using the ratiometric dye SNARF-1.^78^ In the chloride influx assay, the SNARF-1 fluorescence ratio decreased (F_f_/F_i_ = 0.8 ± 0.1), consistent with a mild intracellular acidification where the pH remained above 7 (Figure S31).

Similar responses were observed with DMSO (F_f_/F_i_ = 0.8 ± 0.1) and BAPTA (F_f_/F_i_ = 0.9 ± 0.1), whereas the ratio did not change with IAA-94 (Ff/Fi = 1.04 ± 0.02), CaCCinh-A01 (Ff/Fi = 1.0 ± 0.1), and niclosamide (Ff/Fi = 1.05 ± 0.01) (Figure S32–S36). In the efflux assay, the ratio remained unchanged with DMSO (Ff/Fi = 0.9 ± 0.1) and E_act_ (Ff/Fi = 1.0 ± 0.1) (Figure S37). Within the physiological pH range tested, ChlorON-1-PRO maintained strong fluorescence (Figure S27–S30). Together, these results indicate that changes in ChlorON-1-PRO fluorescence reflect chloride binding, with minimal interference from intracellular pH fluctuations under the conditions tested.

## Conclusion

In this work, we identify a single, evolutionarily conserved gatepost residue – position 139 – as a powerful handle to engineer the next-generation ChlorON, ChlorON-1-PRO. In purified form, ChlorON-1-PRO retains the turn-on sensing mechanism of the ChlorON-1 parent but exhibits markedly enhanced chloride affinity and bound-state brightness, albeit with an attenuated response. MD simulations support a model in which the C139N mutation introduces hydrogen-bonding capacity that globally rigidifies the β-barrel and locally alters the binding pocket while stabilizing the chromophore. The multifactorial nature of these changes highlights the value of MD for resolving integrated dynamic features and guiding future mechanistic investigations. Together, the spectroscopic and computational findings echo the functional role of the analogous 148 position in YFP-derived indicators and support a general role for the gatepost position in tuning anion-sensing potential across fluorescent proteins. These molecular advantages translate into robust utility of ChlorON-1-PRO for probing chloride transport under basal and pharmacologically modulated conditions in the U-2 OS cell model. Under matched experimental conditions, ChlorON-1-PRO produced a dose-dependent turn-on fluorescence response to extracellular chloride concentrations as low as 5 mM, outperforming the ChlorON-1 parent. Notably, the attenuated turn-on response observed in purified form was not reflected in the cellular response of ChlorON-1-PRO. Thus, individual *in vitro* properties do not necessarily translate to or predict indicator performance in cells and must be considered collectively.

Our anion exchange assays with ChlorON-1-PRO provide an orthogonal complement to traditional electrophysiological methods and a modern alternative to historically used GFP-derived indicators. Across inhibitory and activating modulators with differing specificities and mechanisms of action, the indicator reports the net outcome of endogenous chloride transport by directly following changes in the labile chloride pool with spatial and temporal resolution. Because ChlorON-1-PRO directly reports chloride, the assay configuration must account for the chloride gradient and direction of transport to ensure that transport modulation generates a resolvable change. While we focused on CaCC-linked pathways as an initial application, the framework established here can be extended to diverse transport mechanisms and cellular contexts. More broadly, this work positions ChlorON-1-PRO as a next-generation platform technology, and with continued application and refinement, it is poised to advance chloride biology.

## Methods

The methods and data analysis for this manuscript are provided in the Supporting Information.

## Data Availability Statement

All the data in this manuscript are provided in the Main Text and Supporting Information. The corresponding author can be contacted for additional requests.

## Supporting Information

Experimental methods and data (PDF).

## Author contributions

S.C.D. designed and supervised the research project with support from A.R.W. for the computational modeling. J.N.T. carried out wet-lab experiments and data analysis. S.M.P. carried out wet-lab experiments and data analysis for the in-cell characterization of ChlorON-1. V.P. carried out computational experiments and analysis. J.N.T., V.P., A.R.W., and S.C.D. wrote the manuscript with input from S.M.P.

## Acknowledgements

J.N.T. acknowledges the Irving S. Sigal Postdoctoral Fellowship from the American Chemical Society. V.P. acknowledges Wayne State University and the Department of Chemistry for the Thomas C. Rumble University Graduate Fellowship. A.R.W. acknowledges the National Science Foundation (NSF CHE2338804), computational support from the Wayne State University Supercomputing Grid, and donations from Arthur D. Barondes. S.C.D. acknowledges the National Institute of General Medical Sciences of the National Institutes of Health (R35GM128923) and the Eugene McDermott Professorship from UT Dallas. We thank Dr. Ke Ji from the Dodani Lab for helpful discussions. This study does not represent the views of the funding agencies and is the sole responsibility of the authors.

## References

(1) Berend, K.; Van Hulsteijn, L. H.; Gans, R. O. B. Chloride: the queen of electrolytes? Eur. J. Intern. Med. 2012, 23, 203–211. DOI: 10.1016/j.ejim.2011.11.013.

(2) Valdivieso, Á. G.; Santa-Coloma, T. A. The chloride anion as a signalling effector. Biol. Rev. 2019, 94, 1839–1856. DOI: 10.1111/brv.12536.

(3) Raut, S. K.; Singh, K.; Sanghvi, S.; Loyo-Celis, V.; Varghese, L.; Singh, E. R.; Rao, S. G.; Singh, H. Chloride ions in health and disease. Biosci. Rep. 2024, 44, BSR20240029. DOI: 10.1042/bsr20240029.

(4) Edwards, J. C.; Kahl, C. R. Chloride channels of intracellular membranes. FEBS Lett. 2010, 584, 2111. DOI: 10.1016/j.febslet.2010.01.037.

(5) Kahle, K. T.; Khanna, A. R.; Alper, S. L.; Adragna, N. C.; Lauf, P. K.; Sun, D.; Delpire, E. K-Cl cotransporters, cell volume homeostasis, and neurological disease. Trends Mol. Med. 2015, 21, 513–523. DOI: 10.1016/j.molmed.2015.05.008.

(6) Jentsch, T. J.; Pusch, M. CLC chloride channels and transporters: structure, function, physiology, and disease. Physiol. Rev. 2018, 98, 1493–1590. DOI: 10.1152/physrev.00047.2017.

(7) Verkman, A. S.; Galietta, L. J. V. Chloride transport modulators as drug candidates. Am. J. Physiol. Cell Physiol. 2021, 321, C932–C946. DOI: 10.1152/ajpcell.00334.2021.

(8) Skov, M.; Ruijs, T. Q.; Grønnebæk, T. S.; Skals, M.; Riisager, A.; Winther, J. B.; Dybdahl, K. L. T.; Findsen, A.; Morgen, J. J.; Huus, N.;, et al. The CLC-1 chloride channel inhibitor NMD670 improves skeletal muscle function in rat models and patients with myasthenia gravis. Sci. Transl. Med. 2024, 16, eadk9109. DOI: 10.1126/scitranslmed.adk9109.

(9) Levin-Konigsberg, R.; Mitra, K.; Spees, K.; Nigam, A. K.; Liu, K.; Januel, C.; Hivare, P.; Arana, S. M.; Prolo, L. M.; Kundaje, A.;, et al. An SLC12A9-dependent ion transport mechanism maintains lysosomal osmolarity. Dev. Cell 2025, 60, 220–235. DOI: 10.1016/j.devcel.2024.10.003.

(10) Jentsch, T. J.; Stein, V.; Weinreich, F.; Zdebik, A. A. Molecular structure and physiological function of chloride channels. Physiol. Rev. 2002, 82, 503–568. DOI: 10.1152/physrev.00029.2001.

(11) Littler, D. R.; Harrop, S. J.; Fairlie, W. D.; Brown, L. J.; Pankhurst, G. J.; Pankhurst, S.; DeMaere, M. Z.; Campbell, T. J.; Bauskin, A. R.; Tonini, R.;, et al. The intracellular chloride ion channel protein CLIC1 undergoes a redox-controlled structural transition. J. Biol. Chem. 2003, 279, 9298–9305. DOI: 10.1074/jbc.m308444200.

(12) Xiao, Q.; Yu, K.; Perez-Cornejo, P.; Cui, Y.; Arreola, J.; Hartzell, H. C. Voltage- and calcium-dependent gating of TMEM16A/ANO1 chloride channels are physically coupled by the first intracellular loop. Proc. Natl. Acad. Sci. U. S. A. 2011, 108, 8891–8896. DOI: 10.1073/pnas.1102147108.

(13) Sala-Rabanal, M.; Yurtsever, Z.; Nichols, C. G.; Brett, T. J. Secreted CLCA1 modulates TMEM16A to activate Ca^2+^-dependent chloride currents in human cells. eLife 2015, 4, e05875. DOI: 10.7554/elife.05875.

(14) Ruan, Z.; Osei-Owusu, J.; Du, J.; Qiu, Z.; Lü, W. Structures and pH-sensing mechanism of the proton-activated chloride channel. Nature 2020, 588, 350–354. DOI: 10.1038/s41586-020-2875-7.

(15) Della Sala, A.; Prono, G.; Hirsch, E.; Ghigo, A. Role of protein kinase A-mediated phosphorylation in CFTR channel activity regulation. Front. Physiol. 2021, 12, 690247. DOI: 10.3389/fphys.2021.690247.

(16) Varela, L.; Hendry, A. C.; Cassar, J.; Martin-Escolano, R.; Cantoni, D.; Ossa, F.; Edwards, J. C.; Abdul-Salam, V.; Ortega-Roldan, J. L. A Zn^2+^-triggered two-step mechanism of CLIC1 membrane insertion and activation into chloride channels. J. Cell Sci. 2022, 135, jcs259704. DOI: 10.1242/jcs.259704.

(17) Pant, S.; Tam, S. W.; Long, S. B. The pentameric chloride channel BEST1 is activated by extracellular GABA. Proc. Natl. Acad. Sci. U. S. A. 2025, 122, e2424474122. DOI: 10.1073/pnas.2424474122.

(18) Lara-Santos, C.; Tembo, M.; Miller, J. M.; Rosenbaum, J. C.; Carlson, A. E. Interplay of Ca^2+^ and PIP2 in TMEM16A channel function. J. Physiol. 2026, 604, 366–378. DOI: 10.1113/jp289257.

(19) Piala, A. T.; Moon, T. M.; Akella, R.; He, H.; Cobb, M. H.; Goldsmith, E. J. Chloride sensing by WNK1 involves inhibition of autophosphorylation. Sci. Signal. 2014, 7, ra41. DOI: 10.1126/scisignal.2005050.

(20) Valdivieso, Á. G.; Clauzure, M.; Massip-Copiz, M.; Santa-Coloma, T. A. The chloride anion acts as a second messenger in mammalian cells - modifying the expression of specific genes. Cell. Physiol. Biochem. 2016, 38, 49–64. DOI: 10.1159/000438608.

(21) Lüscher, B. P.; Vachel, L.; Ohana, E.; Muallem, S. Cl^-^ as a bona fide signaling ion. Am. J. Physiol. Cell Physiol. 2020, 318, C125–C136. DOI: 10.1152/ajpcell.00354.2019.

(22) Zhang, Z.; Liu, F.; Chen, J. Conformational changes of CFTR upon phosphorylation and ATP binding. Cell 2017, 170, 483–491. DOI: 10.1016/j.cell.2017.06.041.

(23) Palmer, E. E.; Pusch, M.; Picollo, A.; Forwood, C.; Nguyen, M. H.; Suckow, V.; Gibbons, J.; Hoff, A.; Sigfrid, L.; Megarbane, A.; Nizon, M.;, et al. Functional and clinical studies reveal pathophysiological complexity of CLCN4-related neurodevelopmental condition. Mol. Psychiatry 2022, 28, 668–697. DOI: 10.1038/s41380-022-01852-9.

(24) Tippett, D. N.; Breen, C.; Butler, S. J.; Sawicka, M.; Dutzler, R. Structural and functional properties of the transporter SLC26A6 reveal mechanism of coupled anion exchange. eLife 2023, 12, RP87178. DOI: 10.7554/elife.87178.

(25) Davis, O. C.; Ferland, S.; Lorenzo, L. E.; Murray-Lawson, C.; Shiers, S.; Yousuf, M. S.; Dedek, A.; Tsai, E. C.; Vines, E.; Horton, P.;, et al. Decreased KCC2 expression in the human spinal dorsal horn associated with chronic pain and long-term opioid use. Pain 2025, 166, e665–e673. DOI: 10.1097/j.pain.0000000000003700.

(26) Verkman, A. S.; Galietta, L. J. V. Chloride channels as drug targets. Nat. Rev. Drug Discov. 2009, 8, 153–171. DOI: 10.1038/nrd2780.

(27) Drummond-Main, C. D.; Rho, J. M. Electrophysiological characterization of a mitochondrial inner membrane chloride channel in rat brain. FEBS Lett. 2018, 592, 1545–1553. DOI: 10.1002/1873-3468.13042.

(28) Pressey, J. C.; de Saint-Rome, M.; Raveendran, V. A.; Woodin, M. A. Chloride transporters controlling neuronal excitability. Physiol. Rev. 2023, 103, 1095–1135. DOI: 10.1152/physrev.00025.2021.

(29) Xu, M.; Neelands, T.; Powers, A. S.; Liu, Y.; Miller, S. D.; Pintilie, G. D.; Du Bois, J.; Dror, R. O.; Chiu, W.; Maduke, M. CryoEM structures of the human CLC-2 voltage-gated chloride channel reveal a ball-and-chain gating mechanism. eLife 2024, 12, RP90648. DOI: 10.7554/elife.90648.

(30) Wachter, R. M.; Remington, S. J. Sensitivity of the yellow variant of green fluorescent protein to halides and nitrate. Curr. Biol. 1999, 9, R628–R629. DOI: 10.1016/s0960-9822(99)80408-4.

(31) Kuner, T.; Augustine, G. J. A genetically encoded ratiometric indicator for chloride: capturing chloride transients in cultured hippocampal neurons. Neuron 2000, 27, 447–459. DOI: 10.1016/s0896-6273(00)00056-8.

(32) Arosio, D.; Garau, G.; Ricci, F.; Marchetti, L.; Bizzarri, R.; Nifosì, R.; Beltram, F. Spectroscopic and structural study of proton and halide ion cooperative binding to GFP. Biophys. J. 2007, 93, 232–244. DOI: 10.1529/biophysj.106.102319.

(33) Arosio, D.; Ratto, G. M. Twenty years of fluorescence imaging of intracellular chloride. Front. Cell. Neurosci. 2014, 8, 258. DOI: 10.3389/fncel.2014.00258.

(34) Chudakov, D. M.; Matz, M. V.; Lukyanov, S.; Lukyanov, K. A. Fluorescent proteins and their applications in imaging living cells and tissues. Physiol. Rev. 2010, 90, 1103–1163. DOI: 10.1152/physrev.00038.2009.

(35) Zajac, M.; Chakraborty, K.; Saha, S.; Mahadevan, V.; Infield, D. T.; Accardi, A.; Qiu, Z.; Krishnan, Y. What biologists want from their chloride reporters — a conversation between chemists and biologists. J. Cell Sci. 2020, 133, jcs240390. DOI: 10.1242/jcs.240390.

(36) Lodovichi, C.; Ratto, G. M.; Trevelyan, A. J.; Arosio, D. Genetically encoded sensors for chloride concentration. J. Neurosci. Methods 2022, 368, 109455. DOI: 10.1016/j.jneumeth.2021.109455.

(37) Wachter, R. M.; Yarbrough, D.; Kallio, K.; Remington, S. J. Crystallographic and energetic analysis of binding of selected anions to the yellow variants of green fluorescent protein. J. Mol. Biol. 2000, 301, 157–171. DOI: 10.1006/jmbi.2000.3905.

(38) Jayaraman, S.; Haggie, P.; Wachter, R. M.; Remington, S. J.; Verkman, A. S. Mechanism and cellular applications of a green fluorescent protein-based halide sensor. J. Biol. Chem. 2000, 275, 6047–6050. DOI: 10.1074/jbc.275.9.6047.

(39) Galietta, L. J. V; Haggie, P. M.; Verkman, A. S. Green fluorescent protein-based halide indicators with improved chloride and iodide affinities. FEBS Lett. 2001, 499, 220–224. DOI: 10.1016/s0014-5793(01)02561-3.

(40) Galietta, L. J. V; Springsteel, M. F.; Eda, M.; Niedzinski, E. J.; By, K.; Haddadin, M. J.; Kurth, M. J.; Nantz, M. H.; Verkman, A. S. Novel CFTR chloride channel activators identified by screening of combinatorial libraries based on flavone and benzoquinolizinium lead compounds. J. Biol. Chem. 2001, 276, 19723–19728. DOI: 10.1074/jbc.m101892200.

(41) Caputo, A.; Caci, E.; Ferrera, L.; Pedemonte, N.; Barsanti, C.; Sondo, E.; Pfeffer, U.; Ravazzolo, R.; Zegarra-Moran, O.; Galietta, L. J. V. TMEM16A, a membrane protein associated with calcium-dependent chloride channel activity. Science 2008, 322, 590–594. DOI: 10.1126/science.1163518.

(42) Qiu, Z.; Dubin, A. E.; Mathur, J.; Tu, B.; Reddy, K.; Miraglia, L. J.; Reinhardt, J.; Orth, A. P.; Patapoutian, A. SWELL1, a plasma membrane protein, is an essential component of volume-regulated anion channel. Cell 2014, 157, 447–458. DOI: 10.1016/j.cell.2014.03.024.

(43) Fiore, M.; Cossu, C.; Capurro, V.; Picco, C.; Ludovico, A.; Mielczarek, M.; Carreira-Barral, I.; Caci, E.; Baroni, D.; Quesada, R.; Moran, O. Small molecule-facilitated anion transporters in cells for a novel therapeutic approach to cystic fibrosis. Br. J. Pharmacol. 2019, 176, 1764–1779. DOI: 10.1111/bph.14649.

(44) Tutol, J. N.; Peng, W.; Dodani, S. C. Discovery and characterization of a naturally occurring, turn-on yellow fluorescent protein sensor for chloride. Biochemistry 2019, 58, 31–35. DOI: 10.1021/acs.biochem.8b00928.

(45) Tutol, J. N.; Kam, H. C.; Dodani, S. C. Identification of mNeonGreen as a pH-dependent, turn-on fluorescent protein sensor for chloride. ChemBioChem 2019, 20, 1759–1765. DOI: 10.1002/cbic.201900147.

(46) Salto, R.; Giron, M. D.; Puente-Muñ Oz, V.; Vilchez, J. D.; Espinar-Barranco, L.; Valverde-Pozo, J.; Arosio, D.; Paredes, J. M. New red-emitting chloride-sensitive fluorescent protein with biological uses. ACS Sens. 2021, 6, 2563–2573. DOI: 10.1021/acssensors.1c00094.

(47) Shariati, K.; Zhang, Y.; Giubbolini, S.; Parra, R.; Liang, S.; Edwards, A.; Hejtmancik, J. F.; Ratto, G. M.; Arosio, D.; Ku, G. A superfolder green fluorescent protein-based biosensor allows monitoring of chloride in the endoplasmic reticulum. ACS Sens. 2022, 7, 2218–2224. DOI: 10.1021/acssensors.2c00626.

(48) Peng, W.; Maydew, C. C.; Kam, H.; Lynd, J. K.; Tutol, J. N.; Phelps, S. M.; Abeyrathna, S.; Meloni, G.; Dodani, S. C. Discovery of a monomeric green fluorescent protein sensor for chloride by structure-guided bioinformatics. Chem. Sci. 2022, 13, 12659–12672. DOI: 10.1039/d2sc03903f.

(49) Ong, W. S. Y.; Ji, K.; Pathiranage, V.; Maydew, C.; Baek, K.; Villones, R. L. E.; Meloni, G.; Walker, A. R.; Dodani, S. C. Rational design of the β-bulge gate in a green fluorescent protein accelerates the kinetics of sulfate sensing. Angew. Chem. Int. Ed. 2023, 62, e202302304. DOI: 10.1002/anie.202302304.

(50) Peng, W.; Tutol, J. N.; Phelps, S. M.; Kam, H.; Lynd, J. K.; Dodani, S. C. Directed evolution of a genetically encoded indicator for chloride. ACS Synth. Biol. 2025, 14, 1009–1013. DOI: 10.1021/acssynbio.4c00818.

(51) Cook, M. A.; Smailys, J. D.; Ji, K.; Phelps, S. M.; Tutol, J. N.; Kim, W.; Ong, W. S. Y.; Peng, W.; Maydew, C.; Zhang, Y. J.; Dodani, S. C. NitrOFF: An engineered fluorescent biosensor to illuminate nitrate transport in living cells. Angew. Chem. Int. Ed. 2025, 64, e202508058. DOI: 10.1002/anie.202508058.

(52) Tutol, J. N.; Ong, W. S. Y.; Phelps, S. M.; Peng, W.; Goenawan, H.; Dodani, S. C. Engineering the ChlorON series: Turn-on fluorescent protein sensors for imaging labile chloride in living cells. ACS Cent. Sci. 2024, 10, 77–86. DOI: 10.1021/acscentsci.3c01088.

(53) Lai, C.; Yang, L.; Pathiranage, V.; Wang, R.; Subach, F. V.; Walker, A. R.; Piatkevich, K. D. Genetically encoded green fluorescent sensor for probing sulfate transport activity of solute carrier family 26 member a2 (SLC26A2) protein. *Commun*. Biol. 2024, 7, 1375. DOI: 10.1038/s42003-024-07020-9.

(54) Kille, S.; Acevedo-Rocha, C. G.; Parra, L. P.; Zhang, Z.-G.; Opperman, D. J.; Reetz, M. T.; Acevedo, J. P. Reducing codon redundancy and screening effort of combinatorial protein libraries created by saturation mutagenesis. ACS Synth. Biol. 2013, 2, 83–92. DOI: 10.1021/sb300037w.

(55) Chen, C.; Pathiranage, V.; Ong, W. S. Y.; Dodani, S. C.; Walker, A. R.; Fang, C. A twisted chromophore powers a turn-on fluorescent protein chloride sensor. Proc. Natl. Acad. Sci. U. S. A. 2025, 122, e2508094122. DOI: 10.1073/pnas.2508094122.

(56) Mukherjee, S.; Manna, P.; Douglas, N.; Chapagain, P. P.; Jimenez, R. Conformational dynamics of mCherry variants: a link between side-chain motions and fluorescence brightness. J. Phys. Chem. B 2023, 127, 52–61. DOI: 10.1021/acs.jpcb.2c05584.

(57) Niforou, K. N.; Anagnostopoulos, A. K.; Vougas, K.; Kittas, C.; Gorgoulis, V. G.; Tsangaris, G. T. The proteome profile of the human osteosarcoma U2OS cell line. CGP. 2008, 5, 63–77. PMID: 18359981.

(58) Beck, M.; Schmidt, A.; Malmstroem, J.; Claassen, M.; Ori, A.; Szymborska, A.; Herzog, F.; Rinner, O.; Ellenberg, J.; Aebersold, R. The quantitative proteome of a human cell line. Mol. Syst. Biol. 2011, 7, 549. DOI: 10.1038/msb.2011.82.

(59) Karlsson, M.; Zhang, C.; Méar, L.; Zhong, W.; Digre, A.; Katona, B.; Sjöstedt, E.; Butler, L.; Odeberg, J.; Dusart, P.;, et al. A single–cell type transcriptomics map of human tissues. Sci. Adv. 2021, 7, eabh2169. DOI: 10.1126/sciadv.abh2169.

(60) Müller, S. Whole cell proteome of U2OS cells or Saos-2 cells depleted for SENP3. Proteomics Identifications Database 2023, https://www.ebi.ac.uk/pride/archive/projects/PXD037800.

61. (61) *The Human Protein Atlas*, v.25.1. https://www.proteinatlas.org (accessed 2026-08-20).

(62) Reeves, W. B.; Gurich, R. W. Calcium-dependent chloride channels in endosomes from rabbit kidney cortex. Am. J. Physiol. Cell Physiol. 1994, 266, C741–C750. DOI: 10.1152/ajpcell.1994.266.3.c741.

(63) Nelson, M. T.; Conway, M. A.; Knot, H. J.; Brayden, J. E. Chloride channel blockers inhibit myogenic tone in rat cerebral arteries. J. Physiol. 1997, 502, 259–264. DOI: 10.1111/j.1469-7793.1997.259bk.x.

(64) Cheng, G.; Kim, M. J.; Jia, G.; Agrawal, D. K. Involvement of chloride channels in IGF-I-induced proliferation of porcine arterial smooth muscle cells. Cardiovasc. Res. 2006, 73, 198–207. DOI: 10.1016/j.cardiores.2006.10.012.

(65) Zhang, H.; Cao, H. J.; Kimelberg, H. K.; Zhou, M. Volume regulated anion channel currents of rat hippocampal neurons and their contribution to oxygen-and-glucose deprivation induced neuronal death. PLoS One 2011, 6, e16803. DOI: 10.1371/journal.pone.0016803.

(66) Dickson, V. K.; Pedi, L.; Long, S. B. Structure and insights into the function of a Ca^2+^-activated Cl^-^ channel. Nature 2014, 516, 218. DOI: 10.1038/nature13913.

(67) Shi, S.; Guo, S.; Chen, Y.; Sun, F.; Pang, C.; Ma, B.; Qu, C.; An, H. Molecular mechanism of CaCCinh-A01 inhibiting TMEM16A channel. Arch. Biochem. Biophys. 2020, 695, 108650. DOI: 10.1016/j.abb.2020.108650.

(68) De La Fuente, R.; Namkung, W.; Mills, A.; Verkman, A. S. Small-molecule screen identifies inhibitors of a human intestinal calcium-activated chloride channel. Mol. Pharmacol. 2008, 73, 758–768. DOI: 10.1124/mol.107.043208.

(69) Liu, Y.; Zhang, H.; Huang, D.; Qi, J.; Xu, J.; Gao, H.; Du, X.; Gamper, N.; Zhang, H. Characterization of the effects of Cl^−^ channel modulators on TMEM16A and bestrophin-1 Ca^2+^ activated Cl^−^ channels. Pflugers Arch. 2015, 467, 1417–1430. DOI: 10.1007/s00424-014-1572-5.

(70) Bill, A.; Hall, M. L.; Borawski, J.; Hodgson, C.; Jenkins, J.; Piechon, P.; Popa, O.; Rothwell, C.; Tranter, P.; Tria, S.; Wagner, T.; Whitehead, L.; Gaither, L. A. Small molecule-facilitated degradation of ANO1 protein. J. Biol. Chem. 2014, 289, 11029–11041. DOI: 10.1074/jbc.m114.549188.

(71) Chen, W.; Mook, R. A.; Premont, R. T.; Wang, J. Niclosamide: beyond an antihelminthic drug. Cell Signal. 2017, 41, 89–96. DOI: 10.1016/j.cellsig.2017.04.001.

(72) Liang, P.; Wan, Y. C. S.; Yu, K.; Hartzell, H. C.; Yang, H. Niclosamide potentiates TMEM16A and induces vasoconstriction. J. Gen. Physiol. 2024, 156, e202313460. DOI: 10.1085/jgp.202313460/276776.

(73) Ousingsawat, J.; Centeio, R.; Reyne, N.; McCarron, A.; Cmielewski, P.; Schreiber, R.; diStefano, G.; Römermann, D.; Seidler, U.; Donnelley, M.; Kunzelmann, K. Inhibition of mucus secretion by niclosamide and benzbromarone in airways and intestine. Sci. Rep. 2024, 14, 1464. DOI: 10.1038/s41598-024-51397-w.

(74) Ousingsawat, J.; Centeio, R.; Schreiber, R.; Kunzelmann, K. Niclosamide, but not ivermectin, inhibits anoctamin 1 and 6 and attenuates inflammation of the respiratory tract. Pflugers Arch. 2023, 476, 211–227. DOI: 10.1007/s00424-023-02878-w.

(75) Namkung, W.; Yao, Z.; Finkbeiner, W. E.; Verkman, A. S. Small-molecule activators of TMEM16A, a calcium-activated chloride channel, stimulate epithelial chloride secretion and intestinal contraction. The FASEB Journal 2011, 25, 4048–4062. DOI: 10.1096/fj.11-191627.

(76) Liu, S.; Feng, J.; Luo, J.; Yang, P.; Brett, T. J.; Hu, H. Eact, a small molecule activator of TMEM16A, activates trpv1 and elicits pain- and itch-related behaviours. Br. J. Pharmacol. 2016, 173, 1218. DOI: 10.1111/bph.13420.

(77) Rhoden, K. J.; Cianchetta, S.; Duchi, S.; Romeo, G. Fluorescence quantitation of thyrocyte iodide accumulation with the yellow fluorescent protein variant YFP-H148Q/I152L. Anal. Biochem. 2008, 373, 239–246. DOI: 10.1016/j.ab.2007.10.020.

(78) Seksek, O.; Henry-Toulmé, N.; Sureau, F.; Bolard, J. SNARF-1 as an intracellular pH indicator in laser microspectrofluorometry: a critical assessment. Anal. Biochem. 1991, 193, 49–54. DOI: 10.1016/0003-2697(91)90042-r.

