## Supplementary Information for "An All-Optical Approach to Probe Chloride Transport with a Bright ChlorON"

### Table of Contents

**Methods**……S3

**Figures and Tables**……S13
Crystal structures of YFP and YFP-H148Q……S13
ChlorON-1 plasmid design, mutagenesis, library screen, and rescreen
ChlorON-1 nucleotide and amino acid sequences for expression in bacteria cells……S14
C139 site-saturation mutagenesis (SSM) primers and PCR conditions……S15
C139 SSM library results……S16
Rescreen results for the C139 SSM library variants……S17
Summary table of the SSM library and rescreen……S18
Purification and spectroscopic characterization of ChlorON-1-PRO
Size-exclusion chromatogram……S19
SDS-PAGE gel of purified ChlorON-1-PRO……S20
Standard curves for extinction coefficient and quantum yield……S21
Chloride titration at pH 7……S22
pH profile and chromophore p$K_a$ spectra and fits……S23
Anion selectivity bar graph at pH 7……S24
Bromide titration at pH 7……S25
Anion selectivity summary table ……S26
ChlorON-1 chloride titration at pH 7……S27
Computational analysis of ChlorON-1 and ChlorON-1-PRO
Dynamic cross correlation analysis……S28
Normal mode analysis……S29
Water volume calculations……S30
Snapshots of the molecular dynamics simulations……S31
Hydrogen-bond analysis……S32
Spatial distribution of chloride in the binding pocket……S33
Fluorescence microscopy of U-2 OS cells
ChlorON-1-PRO nucleotide and amino acid sequences for expression in mammalian cells……S34
ChlorON-1-PRO with chloride……S35
mNeonGreen anion exchange assay……S36
ChlorON-1-PRO anion exchange assays with modulators……S37
ChlorON-1-PRO pH clamp……S43
SNARF-1 anion exchange microscopy assay and pH clamp……S47
Fluorescence imaging analysis macros……S54

**References**……S56

### Methods

**General.** Except where noted, the supplies, reagents, and equipment used for this manuscript were purchased from Beckman Coulter, Bio-Rad Laboratories, Eppendorf, Fisher Scientific, Gold Biotechnology, Integra Biosciences, Lucigen, Research Products International, Sigma-Aldrich, USA Scientific, VWR, and Zymo Research. The figures with crystal structure and homology models were generated using PyMOL (version 3.1) and MODELLER (version 10.6), respectively (Figure 1, Figure S1).[1,2]

**Bacterial plasmids.** The plasmid encoding ChlorON-1 was transformed using electroporation into *E. cloni* 10G Elite Electrocompetent Cells and extracted as described in our previous study (Figure S2).[3]

**Library cloning, expression, screening, and rescreening.** The site-saturation mutagenesis library was generated as follows. The resulting DNA from above was diluted to 10 ng/µL using autoclaved water and combined with Phusion Hot Start Flex 2X Master Mix, three sets of forward primers, and one reverse primer according to the conditions in Table S1.[4] Next, template DNA was removed by adding 1 µL of DpnI to the PCR tube and incubating at 37 °C for 1 h. The DNA was then isolated, assembled, and concentrated using commercial kits and reagents as described in our previous study.[3] The resulting library DNA (1 µL) and the ChlorON-1 parent DNA (1 µL of 5 ng/µL stock solution) were transformed into *E. cloni* EXPRESS BL21(DE3) Electrocompetent Cells and expressed for screening as described in our previous study with the following modifications.[3] In a 96-well deep well plate, 94 colonies from the ChlorON-1 transformation were picked into 300 µL of 2xYT media (without NaCl) supplemented with 50 µg/mL kanamycin. The two remaining wells were left blank with no colonies. The cells were incubated at 37 °C overnight with shaking at 225 rpm (Innova 42R). The following day, 50 µL of the overnight culture from each well was transferred to 950 µL of 2xYT media (without NaCl) supplemented with 50 µg/mL kanamycin. The cells were further incubated at 37 °C for 2.5 h with shaking at 225 rpm.

Prior to induction, the cells were chilled in an ice water bath for 10 min. To induce protein expression, a solution of 21 mM isopropyl β-D-thiogalactopyranoside (IPTG) was prepared in 2xYT media (without NaCl) and supplemented with 50 µg/mL kanamycin, and 50 µL of this solution was added to each well to a final concentration of 1 mM IPTG. The cells were further incubated at 37 °C for 24 h with shaking at 225 rpm and then at 4 °C overnight without shaking. The cells were harvested via centrifugation at 2,500*g* for 15 min at 4 °C (Eppendorf 5810 R) and stored at -20 °C. For the library expression, 88 variant colonies and 6 colonies of the ChlorON-1 parent were picked into 96-well deep well plate and expressed as described above for the parent plate.

On the day of screening, the parent and library plates were subjected to three freeze-thaw cycles between -20 °C and room temperature for at least 10 min to improve cell lysis. Then, the cell pellets were resuspended in 300 µL of cold lysis buffer (25 mM sodium phosphate buffer at pH 7.4, 2 mM $MgCl_2$, 30 µg/mL DNase I, and 2 mg/mL lysozyme), gently vortexed, and incubated at 37 °C for 1.5 h with shaking at 225 rpm. The cell lysate was clarified via centrifugation at 2,500*g* for 30 min at 4 °C, and 175 µL of lysate from each well was transferred to two transparent 96-well microtiter plates using a liquid handler (Beckman Coulter Biomek NX$^P$). For each well, absorbance spectra were collected from 350–600 nm (3.5 nm bandwidth, 5 nm step-size), and an end-point emission intensity was collected at 515 nm (5 nm bandwidth, gain 100) with excitation provided at 485 nm (5 nm bandwidth) (Figure S3). Next, 25 µL of 25 mM sodium phosphate buffer at pH 7.4 containing 200 mM NaCl was added to each well for a final concentration of 25 mM NaCl. The turn-on fluorescence responses ($F_f/F_i$) in the absence ($F_i$) and presence ($F_f$) of chloride were calculated for each well based on the end-point emission intensity (Figure S3). The fluorescence response of the ChlorON-1 parent plate was used to determine the coefficient of variation which was less than 10%. Given the robustness of the assay, variants with a fluorescence response greater than 2-fold and higher end-point emission intensities than the ChlorON-1 parent on the library plate were selected for sequencing (Eurofins), followed by rescreening (Figure S3). Duplicate sequences were excluded from the rescreening process.

For the rescreen, three colonies of each variant and the ChlorON-1 parent were picked into 5 mL of 2xYT media (without NaCl) and supplemented with 50 µg/mL kanamycin and incubated at 37 °C for 16 h with shaking at 250 rpm. The following day, 200 µL of each overnight culture was added to 5 mL of 2xYT media (without NaCl) supplemented with 50 µg/mL kanamycin in 14 mL culture tubes. The cells were incubated at 37 °C for 2.5 h with shaking at 250 rpm. Prior to induction, the cells were chilled on ice for 10 min. Protein expression was induced with 1 mM IPTG (5 µL from a 1 M stock), and the cells were further incubated at 37 °C for 24 h with shaking at 250 rpm and then at 4 °C overnight without shaking.

The cells were harvested via centrifugation at 2,500*g* for 5 min at 4 °C and stored as pellets at -20 °C. On the day of the rescreen, the cell pellets were resuspended in 100 µL of Bacterial Protein Extraction Reagent (B-PER), vortexed, transferred to 2 mL microcentrifuge tubes, and incubated at room temperature for 15 min. The cell lysate was diluted with 1.1 mL of 50 mM sodium phosphate buffer at pH 7 and clarified via centrifugation at 18,000*g* for 15 min at 4 °C (Eppendorf 5424 R). In a transparent 96-well microtiter plate, 175 µL of each clarified cell lysate was diluted with 25 µL of 50 mM sodium phosphate buffer at pH 7 containing 0, 50, 100, 200, 400, or 800 mM NaCl to a final concentration of 0, 6.3, 12.5, 25, 50, or 100 mM NaCl. For each well, absorbance spectra were collected from 350–600 nm (3.5 nm bandwidth, 5 nm step-size), and excitation was provided at 485 nm (5 nm bandwidth) to collect emission spectra from 505–650 nm (5 nm bandwidth, 5 nm step-size, gain 100) (Figure S4). For the ChlorON-1 parent and each variant, the average fluorescence response at 515 nm along with the apparent dissociation constants from three biological replicates with standard deviation is reported (Table S2).[3]

**ChlorON-1-PRO large-scale expression in bacteria and purification.** The plasmid encoding ChlorON-1-PRO was transformed into *E. cloni* 10G Elite Electrocompetent Cells and extracted for transformation into *E. cloni* EXPRESS BL21(DE3) Electrocompetent Cells as previously described (Figure S2, Table S2).[3] The large-scale expression of ChlorON-1-PRO was carried out as previously described with the following modifications.[3] Two colonies were picked into 30–50 mL of 2xYT media supplemented with 50 µg/mL kanamycin in 250 mL baffled flasks and incubated at 37 °C overnight with shaking at 225 rpm. The next day, ~12 mL of each overnight culture was added to two 2 L baffled flasks containing 600 mL of 2xYT media supplemented with 50 µg/mL kanamycin. The cultures were incubated at 37 °C for 2.5 h with shaking at 200 rpm. Once the optical density at 600 nm reached ~1.2–1.5, protein expression was induced with 600 µL of 1 M IPTG to a final concentration of 1 mM IPTG. The cultures were further incubated at 37 °C for 22 h with shaking at 225 rpm. When this incubation period was completed, the cultures were maintained at 10 °C for at least 8 h without shaking prior to harvesting by centrifugation at

2,500*g* for 30 min at 4 °C. The resulting cell pellets were resuspended in cold 20 mM Tris buffer at pH 7.5 containing 200 mM NaCl, 5 mM $MgCl_2$, 30 µg/mL DNase I, and one XL Protease Inhibitor Tablet for 250 mL of buffer at a ratio of 4 mL of buffer per 1 g of cell mass and stored at -20 °C.

On the day of purification, ~35 mL of the resuspended cells was sonicated on ice for 5 min with an amplitude of 30% and 15 s/45 s on/off cycles (QSonica). The cell lysate was clarified via ultracentrifugation at 18,000*g* for 30 min at 4 °C (Optima XPN-80), and the supernatant was combined for each protein batch. A 5 mL HP HisTrap column was equilibrated with 3 column volumes (CVs) of running buffer composed of 20 mM Tris buffer at pH 7.5 with 200 mM NaCl and 30 mM imidazole. The clarified lysate was loaded onto the HisTrap column at a flow rate of 1 mL/min. The column was then washed with 10 CVs of the running buffer and eluted with a 0–100% gradient of elution buffer composed of 20 mM Tris buffer at pH 7.5 with 200 mM NaCl and 500 mM imidazole at a flow rate of 5 mL/min for 20 CVs. The eluted fractions with absorbance at 480 nm were combined, concentrated to ~10 mL using Amicon centrifugal filters (10 kDa MWCO), and loaded onto a HiPrep 26/10 Desalting column that was pre-equilibrated with 20 mM Tris buffer at pH 7.5 with 200 mM NaCl. The eluted fractions were combined and concentrated to ~11–13 mL before loading onto a HiLoad 26/60 Superdex 200 pg size exclusion column that was equilibrated with 20 mM Tris buffer at pH 7.5 with 200 mM NaCl. For each protein batch, the eluted fractions from ~0.7 CVs with absorbance at 480 nm were combined and concentrated to ~28 mL before transferring to a 30 mL Slide-A-Lyzer Dialysis Cassette (10 kDa MWCO) (Figure S5). Then, the cassettes were placed in a 1 L beaker filled with 20 mM sodium phosphate buffer at pH 7.4 with 50 mM NaCl and stirred at 10 °C for at least 10 h. This process was repeated twice with fresh buffer. After the dialysis, each protein batch was concentrated using centrifugal filters (10 kDa MWCO) to a final concentration of ~500 µM as determined with the Pierce BCA Protein Assay Kit, aliquoted, and stored at -20 °C.

**SDS-PAGE.** The SDS-PAGE and gel staining and destaining with Coomassie dye was adapted from our previous study with the following modifications.[3] Aliquots of purified ChlorON-1-PRO were diluted 100-fold to 5 µM with 50 mM sodium phosphate buffer at pH 7. In a 1.5 mL centrifuge tube, 18 µL of the diluted protein sample was further diluted with 6 µL of 4X Laemmli sample buffer. The samples were incubated for 5 min at 37 °C, and a 20 µL portion was loaded onto a 12% TGX gel along with 5 µL of the PageRuler Plus Prestained Protein Ladder. The electrophoresis was performed at 200 V for 25 min (Figure S6).

**Determination of the protein concentration, extinction coefficient, and quantum yield of ChlorON-1-PRO.** The protein concentration, extinction coefficient, and quantum yield ($\Phi$) of ChlorON-1-PRO in the absence and presence of chloride at pH 7 were determined as described in our previous study with the following modifications.[3] For each protein batch, purified ChlorON-1-PRO was diluted 25-fold to ~20 µM with 50 mM sodium phosphate buffer at pH 7 in two 1.5 mL microcentrifuge tubes for two technical replicates. For each technical replicate, ChlorON-1-PRO was further diluted 2-fold in two 1.5 mL microcentrifuge tubes with 50 mM sodium phosphate buffer at pH 7 containing 0 mM or 400 mM NaCl. Each of these samples was serially diluted 7 times using 50 mM sodium phosphate buffer at pH 7 containing 0 mM or 200 mM NaCl. For the quantum yield standard, serial dilutions of fluorescein disodium salt ($\Phi_{Fluorescein}$ = 0.92) were prepared in fresh 100 mM NaOH.[5] Two hundred microliters of each sample were transferred to a transparent 96-well UV-Star microtiter plate for plate reader measurements.

The following data were used to calculate the protein concentration, extinction coefficient, and quantum yield of ChlorON-1-PRO as described from our previous study.[3] Absorbance spectra were collected from 250–700 nm (3.5 nm bandwidth, 2 nm step-size). The ChlorON-1-PRO absorbance at 490 nm (for 1 mM NaCl) or 494 nm (for 201 mM NaCl) was plotted versus the dilution factor where the linear slopes were used to calculate the protein concentration and extinction coefficient as previously described.[3] For the quantum yield, fluorescence spectra were collected with excitation provided at 488 nm (5 nm bandwidth) and integrated using Microsoft Excel from 506–750 nm (5 nm bandwidth, 2 nm step-size, gain 75) ($F_{Ex488}$). Fluorescence spectra were also collected with excitation provided at 460 nm (5 nm bandwidth) and integrated using Microsoft Excel from 490–750 nm (5 nm bandwidth, 5 nm step-size, gain 75) ($F_{Ex460}$). The ratio of $F_{Ex488}/F_{Ex460}$ was plotted versus the fluorescein and ChlorON-1-PRO absorbance values at 488 nm, and the slopes of these linear plots were used to calculate the quantum yield of ChlorON-1-PRO in the presence of 1 mM and 201 mM NaCl as described in our previous study.[3] The average of four technical replicates across two protein batches with standard deviation is reported (Figure S7, Table 1).

**Plate reader settings for the chloride titration, p$K_a$, and anion selectivity.** Absorbance spectra were collected from 350–600 nm (3.5 nm bandwidth, 5 nm step-size). Fluorescence spectra were collected from 500–650 nm (5 nm bandwidth, 2 nm step-size, gain 75) with excitation provided at 485 nm (5 nm bandwidth). The emission was integrated from 500–650 nm ($F_{Ex485}$) using Microsoft Excel. Fluorescence spectra were also collected from 415–650 nm (5 nm bandwidth, 5 nm step-size, gain 150) with excitation provided at 400 nm (5 nm bandwidth). All experiments were carried out at room temperature from 21–26 °C.

**Chloride titration.** For each technical replicate and protein batch, purified ChlorON-1-PRO was diluted 25-fold to ~20 μM with 50 mM sodium phosphate buffer at pH 7. In 200 μL PCR tubes, ChlorON-1-PRO was further diluted 2-fold with 50 mM sodium phosphate buffer at pH 7 containing 0, 2, 6.3, 12.5, 25, 50, 100, 150, 200, 250, 300, 350, and 400 mM NaCl to a final concentration of ~10 μM protein and 1, 2, 4.1, 7.3, 13.5, 26, 51, 76, 101, 126, 151, 176, and 201 mM NaCl. A portion of each sample (100 μL) was transferred to a transparent 96-well half-area microtiter plate for plate reader measurements. The integrated emission ($F_{Ex485}$) was plotted versus the NaCl concentration which was used to determine apparent dissociation constant ($K_d$) as described in our previous study.[3] The average of four technical replicates across two protein batches with standard deviation is reported (Figure 2, Figure S8, Table 1).

**Chromophore p$K_a$ determination of ChlorON-1-PRO.** For each technical replicate and protein batch, purified ChlorON-1-PRO was diluted 25-fold to ~20 μM with 50 mM sodium acetate buffer at pH 4, 4.5, 5, and 5.5 and 50 mM sodium phosphate buffer at pH 6, 6.5, 7, 7.5, 8, 8.5, and 9. ChlorON-1-PRO was further diluted 2-fold into two sets of 200 μL PCR tubes containing 0 mM or 400 mM NaCl in the corresponding buffer to a final concentration of ~10 μM protein and 1 mM or 201 mM NaCl. A portion of all samples (100 μL) was transferred to a transparent 96-well half-area microtiter plate for plate reader measurements as described above. The fluorescence response ($F_f/F_i$) was calculated by dividing the integrated emission ($F_{Ex485}$) with 201 mM NaCl ($F_f$) by 1 mM NaCl ($F_i$) in their corresponding buffers, and the apo and bound chromophore p$K_a$s were determined as described in our previous study.[3] The average of four technical replicates across two protein batches with standard deviation is reported (Figure S9, Table 1).

**Anion selectivity of ChlorON-1-PRO.** The anion selectivity assay was conducted as described in our previous study with the following modifications.[3] For each technical replicate and protein batch, purified ChlorON-1-PRO was diluted 25-fold to ~20 μM with 50 mM sodium phosphate buffer at pH 7. In 200 μL PCR tubes, ChlorON-1-PRO was further diluted 2-fold with 50 mM sodium phosphate buffer at pH 7 containing 0 mM or 400 mM NaCl, NaBr, NaI, $NaNO_3$, $NaHSO_4$, NaGluconate, NaOAc, and NaCitrate to a final concentration of ~10 μM protein and 1 mM or 201 mM NaCl or 200 mM NaBr, NaI, $NaNO_3$, $NaHSO_4$, NaGluconate, NaOAc, and NaCitrate. For the phosphate selectivity, purified ChlorON-1-PRO was diluted 25-fold to ~20 μM with 50 mM MOPS buffer at pH 7. In 200 μL PCR tubes, ChlorON-1-PRO was further diluted 2-fold with 50 mM MOPS buffer at pH 7 containing 0 mM or 400 mM $NaH_2PO_4$ to a final concentration of ~10 μM protein and 1 mM NaCl or 200 mM $NaH_2PO_4$. A portion of all samples (100 μL) was transferred to a transparent 96-well half-area microtiter plate for plate reader measurements as described above. The fluorescence response ($F_f/F_i$) was calculated by dividing the integrated emission ($F_{Ex485}$) with 201 mM NaCl or 200 mM anion ($F_f$) by 1 mM NaCl ($F_i$) in the corresponding buffers. The average fluorescence response of four technical replicates across two protein batches with standard deviation is reported (Figure 2, Figure S10, Table S3).

For the bromide titration, purified ChlorON-1-PRO was diluted 25-fold to ~20 μM with 50 mM sodium phosphate buffer at pH 7. In 200 μL PCR tubes, ChlorON-1-PRO was further diluted 2-fold with 50 mM sodium phosphate buffer at pH 7 containing 0, 2, 6.3, 12.5, 25, 50, 100, 150, 200, 250, 300, 350, and 400 mM NaBr to a final concentration of ~10 μM protein, 1 mM NaCl, and 0, 1, 3.1, 6.3, 12.5, 25, 50, 75, 100, 125, 150, 175, and 200 mM NaBr. The solutions were then transferred to a transparent 96-well half-area microtiter plate for plate reader measurements, and the apparent dissociation constant was determined as described above. The average of four technical replicates across two protein batches with standard deviation is reported (Figure S11).

**Molecular dynamics simulations for ChlorON-1 and ChlorON-1-PRO.** Classical molecular dynamics (MD) simulations were performed for both the apo and chloride bound states of ChlorON-1-PRO. For comparison, we also incorporated the previously generated MD trajectories of ChlorON-1 from our initial ChlorON-1/ChlorON-3 study, using identical simulation protocols to ensure consistency across all systems.[6] Replicate simulations of ChlorON-1-PRO using Amber20's constant pH MD module (CpHMD) showed minimal effect and residue protonation state changes, and so we proceeded with standard classical MD from then on.

The chloride bound structure of ChlorON-1-PRO was modeled by starting from the X-ray structure of mNeonGreen with bound chloride (PDB ID: 5LTP). Removal of the chloride ion yielded the apo form. Point mutations required for generating the ChlorON-1-PRO variants were introduced using UCSF ChimeraX 1.2.5 with the Dunbrack rotamer library.[7,8] Protonation states for all residues were subsequently optimized using MolProbity 4.5.2 followed by the PDB2PQR web server (version 3.5.2).[9,10] Each system was neutralized by addition of chloride ions and solvated in a rectangular TIP3P water box extending 15 Å from the protein surface.[11]

System parameterization was carried out in AmberTools20 using tleap with the ff14SB force field for the protein.[12,13] Chromophore parameters were supplied from AutoParams.[14] After system construction, we applied an initial round of minimization and density equilibration in the NPT ensemble. During this stage, restraints of 500 kcal/mol were maintained on the protein and chromophore, while the solvent was allowed to relax freely. Additional density equilibration steps were conducted by gradually reducing these restraints at 10 K. The systems were then heated from low temperature to 300 K in the NVT ensemble, reinstating the 500 kcal/mol restraints. A final equilibration phase in the NPT ensemble at 300 K was performed while again lifting the restraints.

Production level MD simulations were run in the NPT ensemble for 100 ns, using a 1 fs integration time step with pmemd.cuda.[15] A Langevin thermostat with a collision frequency of 5 $ps^{-1}$ was used for temperature control.[16] The SHAKE algorithm was applied to constrain bonds involving hydrogen atoms.[17] Non-bonded interactions employed a 10 Å cutoff with smooth particle mesh Ewald treatment for long range electrostatics.[18]

Trajectory analyses including root mean square fluctuation (RMSF), normal mode analysis, and hydrogen bond occupancies were performed with CPPTRAJ in AmberTools20 (Figure 3, Figure S13–S17, Table S4).[19] Structural visualization and figure preparation were carried out with PyMOL (version 3.1).[1] Empty cavity volumes were estimated using a grid-based method inspired by the POVME algorithm.[20] A 3D grid (0.5 Å spacing) was generated around the chromophore, and grid points within a van der Waals radius of any protein atom were removed using SciPy's cKDTree spatial queries.[21,22] The remaining unoccupied grid points were summed to obtain the empty volume. The surface charge distribution was plotted using ChimeraX 1.2.5.

**Mammalian plasmids.** The ChlorON-1-PRO gene was codon optimized for expression in mammalian cells, synthesized, and cloned into the pcDNA-3.1(+)-N-6His vector between the BamHI and EcoRI restriction sites (GenScript, Figure S18).

**U-2 OS cell culture and stable cell line generation.** The U-2 OS cells (HTB-96) used in this study were purchased from the American Type Culture Collection and cultured in McCoy's 5A media supplemented with 10% fetal bovine serum (FBS) and 1% Penicillin-Streptomycin (PS) in a T75 flask. The U-2 OS cell culture and stable cell line generation were conducted as described in our previous study with the following modifications.[23] When the cells were ~100% confluent, the T75 flask was washed with 5 mL Phosphate Buffered Saline (PBS) and incubated with 5 mL of 0.05% trypsin-EDTA at 37 °C, 5% $CO_2$ for 5–7 min, followed by 10 mL of McCoy's 5A media supplemented with 10% FBS and 1% PS to quench the trypsin. After this, the cells were harvested via centrifugation at 300*g* for 5 min at room temperature (Eppendorf 5702) and resuspended in 5 mL of media.

To generate the ChlorON-1-PRO stable cell line, U-2 OS cells were plated at a seeding density of ~6 x $10^5$ cells in 2 mL of McCoy's 5A media supplemented with 10% FBS and 1% PS. One well was seeded at ~3 x $10^5$ cells in 2 mL of media to serve as a negative control. The cells were incubated at 37 °C, 5% $CO_2$ until the next day. When the cells were ~70% confluent, each well was transfected with 2.25 µL ChlorON-1-PRO pcDNA, 4.5 µL P3000 reagent, and 3.75 µL Lipofectamine 3000 in 250 µL Reduced Serum Opti-MEM and incubated at 37 °C, 5% $CO_2$ for 72 h. On the day of sorting, the cells were washed with 2 mL of PBS and treated with 1 mL of 0.05% trypsin-EDTA for 5–7 min and then 2 mL of McCoy's 5A media supplemented with 10% FBS to quench the trypsin. The transfected cells were combined and harvested via centrifugation at 300*g* for 5 min, washed with 5 mL of PBS supplemented with 2% FBS, resuspended in 700 µL of PBS supplemented with 2% FBS, and passed through the strainer cap of a 5 mL flow cytometry tube for single cell sorting. The cells were sorted using the FACSAria Fusion Flow

Cytometer (BD Biosciences) with the GFP emission filter at the UT Dallas Flow Cytometry Core. Non-transfected U-2 OS cells were used to draw gates for the negative population and set the threshold for the positively transfected and emissive population. All cells expressing ChlorON-1-PRO were collected in a 14 mL centrifuge tube containing McCoy's 5A media supplemented with 10% FBS. Next, the cells were harvested via centrifugation at 200*g* for 5 min, resuspended in McCoy's 5A media supplemented with 10% FBS and 500 µg/mL G418 to select for cells with the ChlorON-1-PRO plasmid, transferred to a T25 flask, and incubated at 37 °C, 5% $CO_2$. Once the cells reached 100% confluency, the cells were passaged and sorted as described above. After three total rounds of sorting and when ~95% of the cell population emitted green fluorescence, U-2 OS cells expressing ChlorON-1-PRO were sorted a final time based on populations with the top 25%, middle 25%, and bottom 25% having high, medium, and low fluorescence emission intensities, respectively. The sorted populations were then grown in three T25 flasks containing McCoy's 5A media supplemented with 10% FBS and 500 µg/mL G418 as described above. The ChlorON-1-PRO stable cell line from the medium fluorescence population was used for the imaging experiments described below. U-2 OS cells stably expressing ChlorON-1 were generated under the same conditions as described with the following modification. Cells expressing ChlorON-1 were sorted a final time based on the top 50% of fluorescent population.

**U-2 OS cell plating for microscopy assays.** For the microscopy assays, 35 mm imaging dishes with a 14 mm glass-like polymer and #1.5 cover slip were seeded with ~5–9.5 x $10^5$ U-2 OS cells in 2 mL of McCoy's 5A media supplemented with 10% FBS and 1% PS and cells stably expressing ChlorON-1-PRO in 2 mL of McCoy's 5A media supplemented with 10% FBS and 500 µg/mL G418. The cells were incubated at 37 °C, 5% $CO_2$ for 24–72 h until at least 80% confluent.

**U-2 OS cell plating and transient transfection with mNeonGreen for microscopy assays.** The mNeonGreen pcDNA-3.1(+)-N-6His plasmid was generated and used to transfect U-2 OS cells as previously described with the following modifications.[3] Imaging dishes were seeded with ~2.5 x $10^5$ cells in 2 mL of McCoy's 5A media supplemented with 10% FBS and 1% PS as described above. After 48 h, the cells were transfected with 100 µL EC Buffer, 3.2 µL Enhancer reagent, 0.4 µg plasmid, and 10 µL Effectene (Qiagen) according to the manufacturer's instructions. The cells were incubated at 37 °C, 5% $CO_2$ for 48 h prior to imaging.

**General microscope specifications and settings.** The microscopy images were captured using an inverted fluorescence microscope (IX83, Olympus) with a 20X air objective (numerical aperture: 0.7; working distance: 1.6 mm), a light engine (Spectra X, Lumencor), and a stage incubator and automated perfusion system (Tokai Hit) set to 37 °C as described in our previous study.[23] For all microscopy assays, differential interference contrast (DIC) and fluorescence images were captured with 25% LED power, a resolution of 512 x 512 pixels (2 x 2 binning), and Z-drift compensation. The DIC exposure time ranged from 1–2.4 ms. The position of each field was recorded in the CellSens software to image the same field before and after treatment.

**Anion exchange microscopy assays with ChlorON-1-PRO, ChlorON-1, and mNeonGreen.** Fluorescence images of U-2 OS cells stably expressing ChlorON-1-PRO were collected in a gluconate (Gluc) buffer containing 137 mM NaGluc, 8.1 mM $Na_2HPO_4$, 1.5 mM $KH_2PO_4$, 2.7 mM KGluc, 1.1 mM $MgGluc_2$, 0.7 mM $CaGluc_2$, and 10 mM glucose at pH 7.4 and a chloride ($Cl^-$) buffer containing 137 mM

NaCl, 8.1 mM $Na_2HPO_4$, 1.5 mM $KH_2PO_4$, 2.7 mM KCl, 1.1 mM $MgCl_2$, 0.7 mM $CaCl_2$, and 10 mM glucose at pH 7.4. For the chloride titration assay, the gluconate and chloride buffers were mixed in varying ratios to the final concentrations of 0, 5, 10, 20, 34.3, 68.5, 100 mM and 137 mM NaCl with NaGluc to maintain the ionic strength. For all buffers, the osmolarity was measured to ~250 mOsm (VAPRO, ELITechGroup). Prior to use, the buffers were warmed to 37 °C. The imaging dishes were washed twice with 2 mL of 137 mM NaGluc buffer and incubated at 37 °C for 30 min. Images were then acquired using an EGFP filter set (Chroma) with excitation provided at 488 nm and an exposure time of 75–175 ms. Next, the imaging solution was exchanged with 8 mL of 132 mM NaGluc/5 mM NaCl buffer at a flow rate of 4 mL/min for 2 min. The same fields of cells were imaged after a 15 min incubation period. The perfusion step was then repeated with 127 mM NaGluc/10 mM NaCl buffer, and images were acquired after a 15 min incubation period. This process was repeated for each chloride concentration. The turn-on emission response ($F_f/F_i$) was calculated by dividing the fluorescence emission at each chloride concentration ($F_f$) by the fluorescence emission in the presence of 137 mM NaGluc ($F_i$). Three different fields were sampled for each dish, and the average emission responses ($F_f/F_i$) from three different cell passages with standard deviations are reported (Figure 4). The chloride titration was repeated with U-2 OS cells stably expressing ChlorON-1. Images were acquired using an exposure time of 400 ms. Three different fields were sampled for each dish, and the emission responses ($F_f/F_i$) from two different cell passages with standard deviations are reported (Figure 4).

For the gluconate-to-chloride exchange assay with cells transiently expressing mNeonGreen, the cells were washed twice with 137 mM NaGluc buffer and incubated at 37 °C for 30 min. Images were acquired every 30 s for 5 min with an exposure time of 15 ms using the EGFP filter set. Next, the imaging solution was exchanged with 8 mL of 68.5 mM NaGluc/68.5 mM NaCl buffer at a flow rate of 4 mL/min for 2 min. Images were acquired again every 30 s for 30 min. The average emission responses ($F_f/F_i$) of three different fields from one biological replicate with standard deviation is reported (Figure S20).

For the anion exchange assays with modulators with cells stably expressing ChlorON-1-PRO, each compound was dissolved in DMSO to generate stock solutions of 56 mM IAA-94, 20 mM BAPTA-AM, 10 mM CaCCinh-A01, 5 mM niclosamide, and 10 mM $E_{act}$ and diluted in buffer to the final concentrations as follows. The U-2 OS cells stably expressing ChlorON-1-PRO were washed with 137 mM NaGluc buffer, treated with 137 mM NaGluc buffer containing 0.1–0.2% DMSO (v/v) as a vehicle control, 100 µM IAA-94, 20 µM BAPTA-AM, 20 µM CaCCinh-A01, 5 µM niclosamide, or 10 µM $E_{act}$, and incubated at 37 °C for 30 min. Images were acquired for 5 min as described above. Following this, the imaging solution was exchanged with 8 mL of 68.5 mM NaGluc/68.5 mM NaCl buffer containing the corresponding treatments at a flow rate of 4 mL/min for 2 min. Images were acquired again every 30 s for 30 min. Three different fields were sampled for each dish. The average emission responses with standard deviations from at least three different cell passages are reported (Figure 5, Figure S21–S26).

For the chloride efflux assay, U-2 OS cells stably expressing ChlorON-1-PRO were seeded in imaging dishes as described above. On the day of imaging, each dish was washed twice with 2 mL of 137 mM NaCl buffer and treated with buffer containing 0.1% DMSO or 10 µM $E_{act}$ at 37 °C for 30 min. Fluorescence images were acquired every 30 s for 5 min with an exposure time of 300 ms. The imaging solution was then exchanged with 8 mL of 137 mM NaGluc buffer containing 0.1% DMSO or 10 µM $E_{act}$ as described above. Images were reacquired every 30 s for 30 min. Lastly, 137 mM NaCl buffer containing 0.1% DMSO or 10 μM $E_{act}$ was re-perfused, and images were acquired every 30 s for 30 min. The average emission responses of three different fields from three biological replicates with standard deviation is reported (Figure 6).

**Anion exchange microscopy assays with ChlorON-1-PRO from pH 6.5 to 8.** The intracellular pH of U-2 OS cells stably expressing ChlorON-1-PRO was calibrated to external clamping buffers at pH 6.5, 7, 7.5, and 8. Each buffer consisted of 140 mM Gluc buffer (120 mM KGluc, 20 mM NaGluc, 8.1 mM $Na_2HPO_4$, 1.5 mM $KH_2PO_4$, 1.1 mM $MgGluc_2$, and 0.7 mM $CaGluc_2$) and 70 mM Gluc/70 mM $Cl^-$ buffer (60 mM KGluc/60 mM KCl, 10 mM NaGluc/10 mM NaCl, 8.1 mM $Na_2HPO_4$, 1.5 mM $KH_2PO_4$, 0.55 mM $MgGluc_2$/0.55 mM $MgCl_2$, 0.35 mM $CaGluc_2$/0.35 mM $CaCl_2$). After adjusting the pH, each buffer was supplemented with 5 μM valinomycin and 5 μM nigericin from a 500X stock solution of 2.5 mM valinomycin and 2.5 mM nigericin in DMSO.

On the day of imaging, two imaging dishes were washed with 2 mL of 137 mM NaGluc buffer at pH 7.4 and incubated at 37 °C for 30 min as described above, and baseline images were acquired at $t$ = 0 min and $t$ = 5 min. The imaging solution was manually exchanged on stage with 4 mL of 140 mM Gluc clamping buffer at pH 6.5, 7, 7.5, or 8 containing 5 μM valinomycin and 5 μM nigericin, and the dish was incubated on stage at 37 °C for 15 min prior to image acquisition. Following this, the imaging solution was manually exchanged with 4 mL of 70 mM Gluc/70 mM $Cl^-$ clamping buffer at the corresponding pH. Images were captured after 15 min and 30 min of incubation on the stage with an exposure time of 125 ms. Three different fields were sampled for each dish. The average emission responses with standard deviations from three different cell passages are reported (Figure S27–S30).

**U-2 OS cell staining and microscopy assays with the pH dye SNARF-1.** Anion exchange assays were carried out as described above with the pH dye SNARF-1 as follows.[24] Fifty micrograms of the acetoxymethyl ester form of 5-(and-6) Carboxy SNARF-1-AM was dissolved in 8.8 μL DMSO to a stock concentration of 10 mM. On the day of imaging, the imaging dishes were washed with 2 mL of DMEM containing 1% PS, stained with 5 μM SNARF-1 in DMEM containing 1% PS, and incubated at 37 °C, 5% $CO_2$ for 1 h. The cells were washed twice with 137 mM NaGluc buffer and incubated at 37 °C for 30 min. Images were acquired at $t$ = 0 min and 5 min. Next, the buffer was exchanged with 8 mL of 68.5 mM NaGluc/68.5 mM NaCl or 137 mM NaCl at pH 7.4 at a flow rate of 4 mL/min for 2 min. The dish was incubated at 37 °C for 30 min, and images were acquired at $t$ = 35 min. Next, the pH was clamped by manually exchanging the imaging solution with 4 mL of the 70 mM Gluc/70 mM $Cl^-$ clamping buffer at pH 6 containing 5 μM valinomycin and 5 μM nigericin as described above. The dish was incubated on stage for 15 min, and the images were acquired. This process was repeated two more times for the clamping buffers at pH 7 and pH 8. Images were acquired with an exposure time of 200 ms using the SNARF-1 filter set with the excitation centered at 500 nm (20 nm bandwidth), and the emission centered at 585 nm (20 nm bandwidth) ($F_{Yellow}$) and 640 nm (20 nm bandwidth) ($F_{Red}$) (Chroma). Two different fields were imaged for each dish. The average ratiometric emission responses with standard deviations from three different cell passage numbers are reported (Figure S31). The gluconate-to-chloride anion exchange assay was repeated with the 0.2% DMSO, 100 μM IAA-94, 20 μM BAPTA-AM, 20 μM CaCCinh-A01, or 5 μM niclosamide treatments described above (Figure S32–S36). The chloride efflux assay was repeated with 0.1% DMSO and 10 μM $E_{act}$ as described above (Figure S37).

**Image processing and analysis.** Microscopy data were analyzed using Fiji Is Just ImageJ software (Fiji v2.0) as described in our previous study with the following modifications.[23,25] For all anion exchange assays and ChlorON-1-PRO pH calibrations, the image files for one field were first concatenated into a single file, and then all channels were split. For each fluorescence channel, the StackReg plugin in Translation mode was used to align each frame, and the background was subtracted with a pixel size of 75. Then, the maximum intensity Z-projections for ChlorON-1-PRO, and the $F_{Yellow}$ channel for SNARF-1 were used to create masks with the minimum threshold set to 15 for ChlorON-1-PRO and 200 for SNARF-1 ($F_{Yellow}$). For the chloride efflux assay, the minimum threshold for ChlorON-1-PRO was set to 100. The Watershed and Analyze Particles functions were used to break down the mask into smaller areas and select regions of interest (ROI), respectively. For each resulting ROI, the median fluorescence intensity was recorded (Figure S38–S39).

For the ChlorON-1-PRO emission response ($F/F_i$) in the absence ($F_i$) and presence of chloride (F), outliers were calculated using the following equation where the first quartile (Q1), third quartile (Q3), and the interquartile range (Q3 – Q1) were determined with Microsoft Excel. All data points that fell outside of the lower and upper range were considered outliers and removed from our analysis.

$$\text{Lower range: } Q1 - (IQR*1.5)$$
$$\text{Upper range: } Q3 + (IQR*1.5)$$

For the chloride influx assay with modulators, the statistical significance (*p*-value) of the ChlorON-1-PRO emissions, SNARF-1 ratiometric emissions, and turn-on responses ($F/F_i$) with chloride were determined relative to $F_i$ using the unpaired Student's *t*-test calculator on GraphPad where *N* corresponds to the number of biological replicates for each dataset.[26]

### Figures and Tables

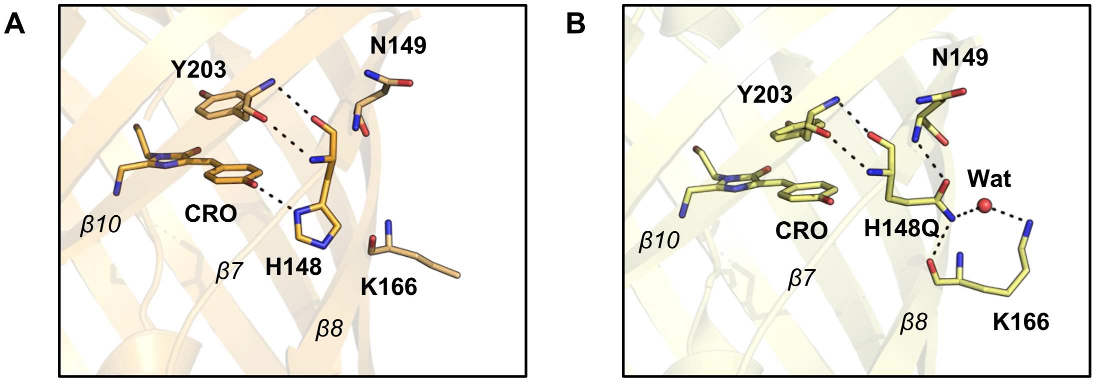


**Figure S1.** Comparison of the hydrogen bonding interactions (dashed lines) in the chromophore region of (A) YFP (PDB ID: 1YFP) and (B) YFP-H148Q (PDB ID: 1F09). Key β-strands are labeled. Abbreviations: CRO, chromophore; Wat, water.

| | | | | | | | | | | | | | | | |
|---|---|---|---|---|---|---|---|---|---|---|---|---|---|---|---|
| | atg | ggc | agc | agc | cat | cat | cat | cat | cat | cac | agc | agc | ggc | ctg | gtg |
| | Met | Gly | Ser | Ser | His | His | His | His | His | His | Ser | Ser | Gly | Leu | Val |
| | ccg | cgc | ggc | agc | cat | atg | gtg | agc | aag | ggc | gag | gaa | gac | aac | atg |
| | Pro | Arg | Gly | Ser | His | Met | Val | Ser | Lys | Gly | Glu | Glu | Asp | Asn | Met |
| | gcg | agc | ctg | ccg | gcg | acc | cat | gag | ctg | cac | atc | ttc | ggc | agc | att |
| 1 | Ala | Ser | Leu | Pro | Ala | Thr | His | Glu | Leu | His | Ile | Phe | Gly | Ser | Ile |
| | aac | ggt | gtg | gac | ttt | gat | atg | gtt | ggt | cag | ggc | acc | ggt | aac | ccg |
| 16 | Asn | Gly | Val | Asp | Phe | Asp | Met | Val | Gly | Gln | Gly | Thr | Gly | Asn | Pro |
| | aac | gac | ggc | tac | gag | gaa | ctg | aac | ctg | aag | agc | acc | aaa | ggt | gat |
| 31 | Asn | Asp | Gly | Tyr | Glu | Glu | Leu | Asn | Leu | Lys | Ser | Thr | Lys | Gly | Asp |
| | ctg | caa | ttc | agc | ccg | tgg | att | ctg | gtg | ccg | cac | att | ggc | tat | ggt |
| 46 | Leu | Gln | Phe | Ser | Pro | Trp | Ile | Leu | Val | Pro | His | Ile | Gly | Tyr | Gly |
| | ttt | cac | cag | tat | ctg | ccg | tat | ccg | gat | ggt | atg | agc | ccg | ttc | caa |
| 61 | Phe | His | Gln | Tyr | Leu | Pro | Tyr | Pro | Asp | Gly | Met | Ser | Pro | Phe | Gln |
| | gcg | gcg | atg | gtg | gat | ggc | agc | ggt | tac | cag | gtt | cac | cgt | acc | atg |
| 76 | Ala | Ala | Met | Val | Asp | Gly | Ser | Gly | Tyr | Gln | Val | His | Arg | Thr | Met |
| | caa | ttt | gaa | gac | ggt | gcg | agc | ctg | acc | gtt | aac | tac | cgt | tat | acc |
| 91 | Gln | Phe | Glu | Asp | Gly | Ala | Ser | Leu | Thr | Val | Asn | Tyr | Arg | Tyr | Thr |
| | tac | gag | ggc | agc | cac | atc | aag | ggt | gaa | gcg | cag | gtg | aag | ggt | acc |
| 106 | Tyr | Glu | Gly | Ser | His | Ile | Lys | Gly | Glu | Ala | Gln | Val | Lys | Gly | Thr |
| | ggt | ttc | ccg | gcg | gat | ggt | ccg | gtt | atg | acc | aac | agc | ctg | acc | gcg |
| 121 | Gly | Phe | Pro | Ala | Asp | Gly | Pro | Val | Met | Thr | Asn | Ser | Leu | Thr | Ala |
| | gcg | gac | tgg | tgc | cgt | agc | aag | tgg | acc | tat | ccg | aac | gat | aag | acc |
| 136 | Ala | Asp | Trp | Cys | Arg | Ser | Lys | Trp | Thr | Tyr | Pro | Asn | Asp | Lys | Thr |
| | atc | att | agc | acc | ttt | aaa | tgg | agc | tat | acc | acc | ggc | aac | ggt | aaa |
| 151 | Ile | Ile | Ser | Thr | Phe | Lys | Trp | Ser | Tyr | Thr | Thr | Gly | Asn | Gly | Lys |
| | cgt | tac | cgt | agc | acc | gcg | cgt | acc | acc | tat | acc | ttt | gcg | aag | ccg |
| 166 | Arg | Tyr | Arg | Ser | Thr | Ala | Arg | Thr | Thr | Tyr | Thr | Phe | Ala | Lys | Pro |
| | atg | gcg | gcg | aac | tat | ctg | aaa | aac | cag | ccg | atg | tac | gtg | ttc | ctg |
| 181 | Met | Ala | Ala | Asn | Tyr | Leu | Lys | Asn | Gln | Pro | Met | Tyr | Val | Phe | Leu |
| | aag | acc | gag | ctg | aag | cac | agc | aaa | acc | gag | ctg | aac | ttc | aag | gaa |
| 196 | Lys | Thr | Glu | Leu | Lys | His | Ser | Lys | Thr | Glu | Leu | Asn | Phe | Lys | Glu |
| | tgg | caa | aaa | gcg | ttt | acc | gac | gtt | atg | ggt | atg | gat | gaa | ctg | tac |
| 211 | Trp | Gln | Lys | Ala | Phe | Thr | Asp | Val | Met | Gly | Met | Asp | Glu | Leu | Tyr |
| | aaa | tga | gga | tcc | | | | | | | | | | | |
| 226 | Lys | * | Gly | Ser | | | | | | | | | | | |

**Figure S2.** The nucleotide and amino acid sequences for ChlorON-1 in the pET28b(+) vector with an N-terminal polyhistidine tag (highlighted in cyan). The ChlorON-1 sequence (green text) is cloned between the NdeI and BamHI restriction sites (highlighted in gray) with the K143W and R195L mutations (blue text). The stop codon is shown as an asterisk (*). The C139 site that was selected for mutagenesis is highlighted in yellow. The codons at position 139 encoding for the C139H variant and ChlorON-1-PRO are listed in Table S2.

**Table S1.** The primers and polymerase chain reaction (PCR) conditions used to generate the ChlorON-1 site-saturation mutagenesis (SSM) library at position 139 (red text).

<table>
<tr><th colspan="4">Primer Design</th></tr>
<tr><th colspan="2">Description</th><th colspan="2">Primer Sequence (5' to 3')</th></tr>
<tr><td colspan="2" rowspan="3">139 SSM Forward Primers</td><td colspan="2">CTGACCGCGGCGGACTGGNDTCGTAGCAAGCGTACCT</td></tr>
<tr><td colspan="2">CTGACCGCGGCGGACTGGVHGCGTAGCAAGCGTACCT</td></tr>
<tr><td colspan="2">CTGACCGCGGCGGACTGGTGGCGTAGCAAGCGTACCT</td></tr>
<tr><td colspan="2">139 SSM Reverse Primer</td><td colspan="2">CCAGTCCGCCGCGGTCAGGCTGTTGGTCATAACC</td></tr>
<tr><th colspan="4">PCR Volumes</th></tr>
<tr><th colspan="2">Solution</th><th>Concentration</th><th>Volume (μL)</th></tr>
<tr><td colspan="2">ChlorON-1 Template</td><td>10 ng/μL</td><td>1</td></tr>
<tr><td colspan="2">139 SSM Forward Primer</td><td>10 μM</td><td>1.5</td></tr>
<tr><td colspan="2">139 SSM Reverse Primer</td><td>10 μM</td><td>1.5</td></tr>
<tr><td colspan="2">Phusion Hot Start Flex</td><td>2X</td><td>12.5</td></tr>
<tr><td colspan="2">Autoclaved Water</td><td>–</td><td>8.5</td></tr>
<tr><td colspan="2">Total</td><td>–</td><td>25</td></tr>
<tr><th colspan="4">PCR Thermocycler Settings</th></tr>
<tr><th>Step</th><th>Temperature (°C)</th><th>Time (s)</th><th>Cycle Number</th></tr>
<tr><td rowspan="2">Template Denaturation</td><td>95</td><td>30</td><td>1</td></tr>
<tr><td>95</td><td>10</td><td rowspan="3">30</td></tr>
<tr><td>Annealing</td><td>56.3</td><td>30</td></tr>
<tr><td>Short Extension</td><td>72</td><td>275</td></tr>
<tr><td>Long Extension</td><td>72</td><td>600</td><td>1</td></tr>
</table>

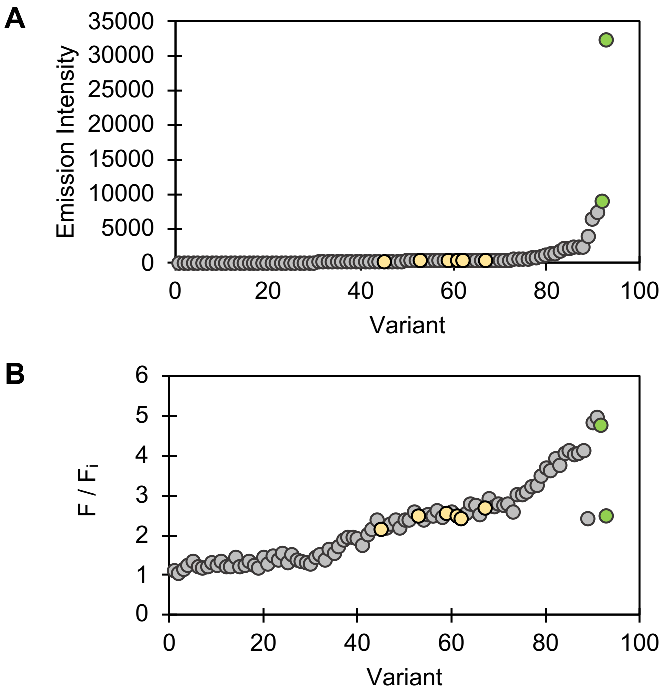


**Figure S3.** Results of the ChlorON-1 SSM library at position 139. (A) The emission intensities at 515 nm ($\lambda_{Ex}$ = 485 nm) with 25 mM NaCl and (B) turn-on responses ($F/F_i$) in the presence of 0 mM ($F_i$) and 25 mM NaCl (F) are shown for the ChlorON-1 parent (yellow circles), all variants (gray circles), and the top two variants with the highest emission intensity (green circles).

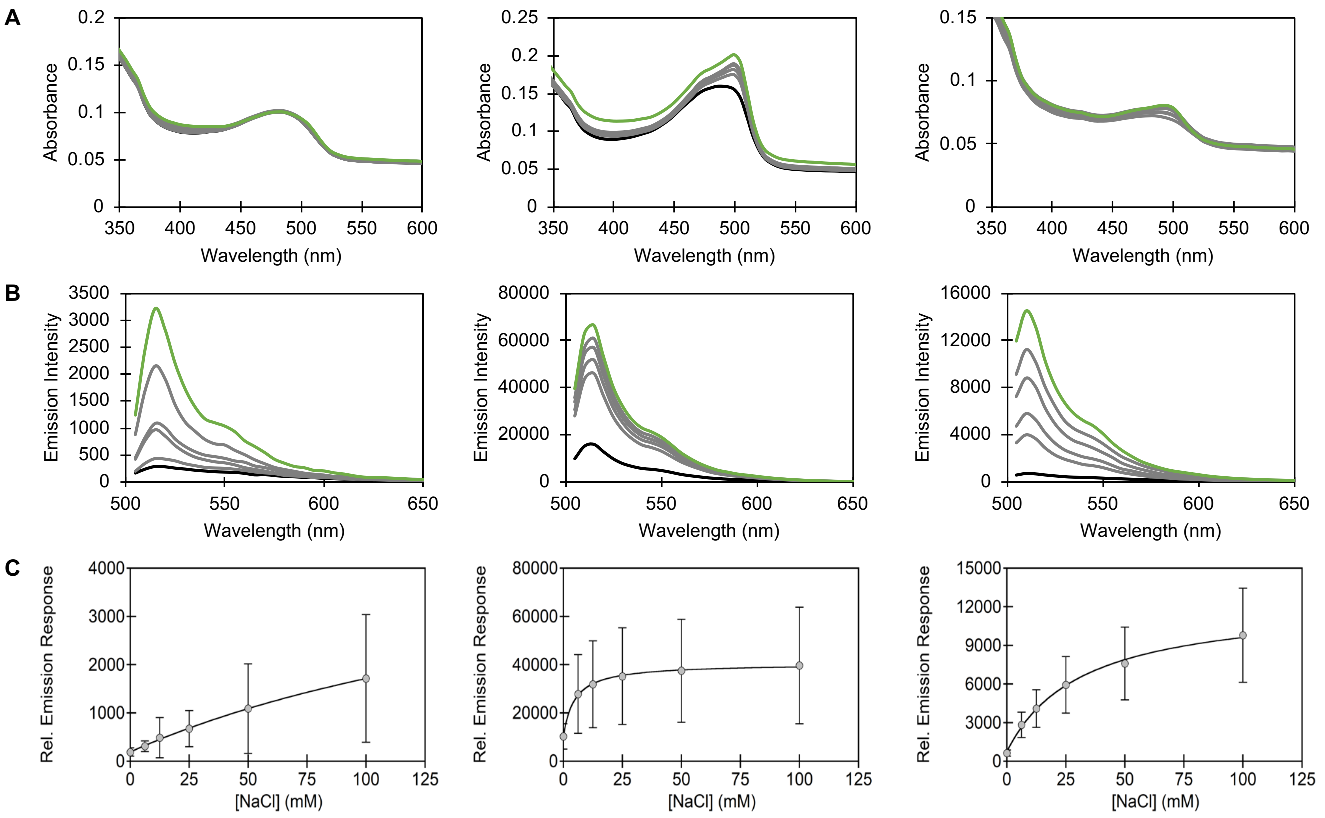


**Figure S4.** Representative (A) absorbance and (B) emission spectra for the ChlorON-1 parent (left), ChlorON-1 C139H (middle), and ChlorON-1 C139N (ChlorON-1-PRO, right) from the rescreen in *E. coli* lysate. All spectra were collected in 50 mM sodium phosphate buffer at pH 7 with 0 (black trace), 6.25, 12.5, 25, 50 (gray traces), and 100 (green trace) mM NaCl. Absorbance spectra were collected from 350–600 nm, and the fluorescence spectra were collected from 505–650 nm with excitation provided at 485 nm. (C) Relative emission response at 515 nm ($\lambda_{Ex}$ = 485 nm) for ChlorON-1 (left), ChlorON-1 C139H (middle), and ChlorON-1-PRO (right) used to determine the dissociation constant ($K_d$). The average of three biological replicates with standard deviation is reported (Table S2).

**Table S2.** Summary of the top variants from the C139 site-saturation mutagenesis library.

| Protein | Mutations | Nucleotide | $F_f$ / $F_i$ [a] | $F_f$ / $F_i$ ± SD [b] | $K_d$ ± SD (mM) [c] |
|---|---|---|---|---|---|
| ChlorON-1 | – | – | 2.46 | 8.24 ± 2.8 | 205 ± 34 |
| ChlorON-1-C139H | C139H | CAT | 2.47 | 3.76 ± 0.40 | 4.66 ± 0.37 |
| ChlorON-1-PRO | C139N | AAT | 4.73 | 15.8 ± 3.1 | 31.5 ± 1.2 |

[a] *Turn-on fluorescence response ($F_f$ / $F_i$) to 0 mM ($F_i$) and 25 mM NaCl ($F_f$) from the library screening in cell lysate in 20 mM sodium phosphate buffer at pH 7.4.*

[b] *Turn-on fluorescence response ($F_f$ / $F_i$) to 0 mM ($F_i$) and 100 mM NaCl ($F_f$) from the rescreening in cell lysate in 50 mM sodium phosphate buffer at pH 7. The average of three biological replicates with SD (standard deviation) is reported.*

[c] *Apparent dissociation constants ($K_d$) determined from the rescreening data in cell lysate from Figure S4. The average of three biological replicates with SD is reported.*

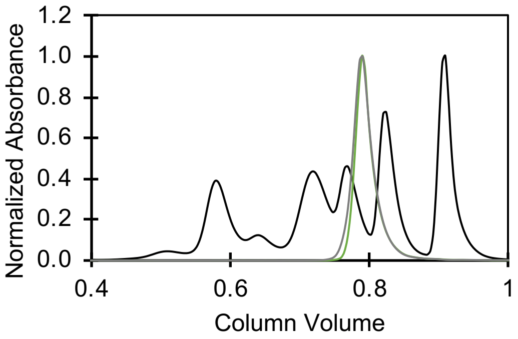


**Figure S5.** Size exclusion chromatography trace for batch 1 (green) and batch 2 (gray) of ChlorON-1-PRO with a protein standard (black). The normalized absorbance of each batch at 490 nm is shown relative to the normalized absorbance at 280 nm for the protein standard with the peaks corresponding to thyroglobulin (670 kDa), gamma globulin (158 kDa), ovalbumin (44 kDa), myoglobin (17 kDa), and vitamin B12 (1.35 kDa). The vitamin B12 absorbance was used to normalize the protein standard.

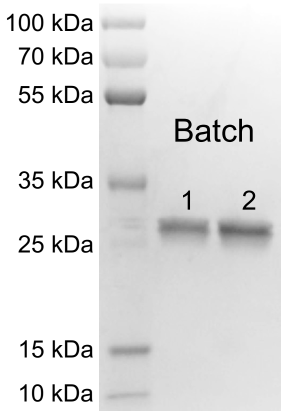


**Figure S6.** SDS-PAGE gel stained with Coomassie dye showing the protein ladder (left), batch 1 of purified ChlorON-1-PRO (middle), and batch 2 of purified ChlorON-1-PRO (right). The theoretical molecular weight for ChlorON-1-PRO is ~26.6 kDa.

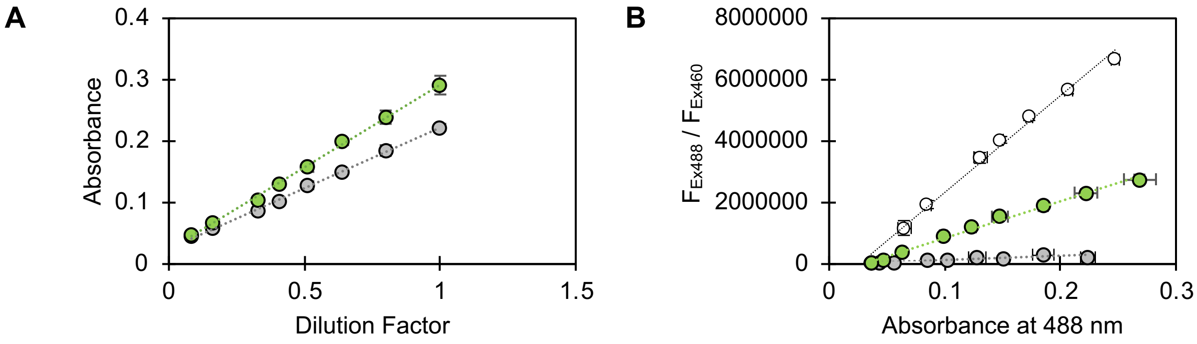


**Figure S7.** Standard curves used to determine the (A) extinction coefficient and (B) quantum yield of ChlorON-1-PRO with 1 mM (gray circles) and 201 mM (green circles) NaCl. The absorbance of ChlorON-1-PRO at 490 nm and 494 nm are shown in panel A for 1 mM and 201 mM NaCl, respectively, versus the dilution factor of each sample. The quantum yield was determined using fluorescein as a reference (white circles) in panel B. The ratios ($F_{Ex488}$/ $F_{Ex460}$) of the integrated emissions from 506–750 nm with excitation provided at 488 nm ($F_{Ex488}$) and from 490–750 nm with excitation provided at 460 nm ($F_{Ex460}$) are plotted versus the absorbance values at 488 nm in panel B. Linear trendlines are shown as dotted lines. The average of four technical replicates across two protein batches with standard deviation is reported.

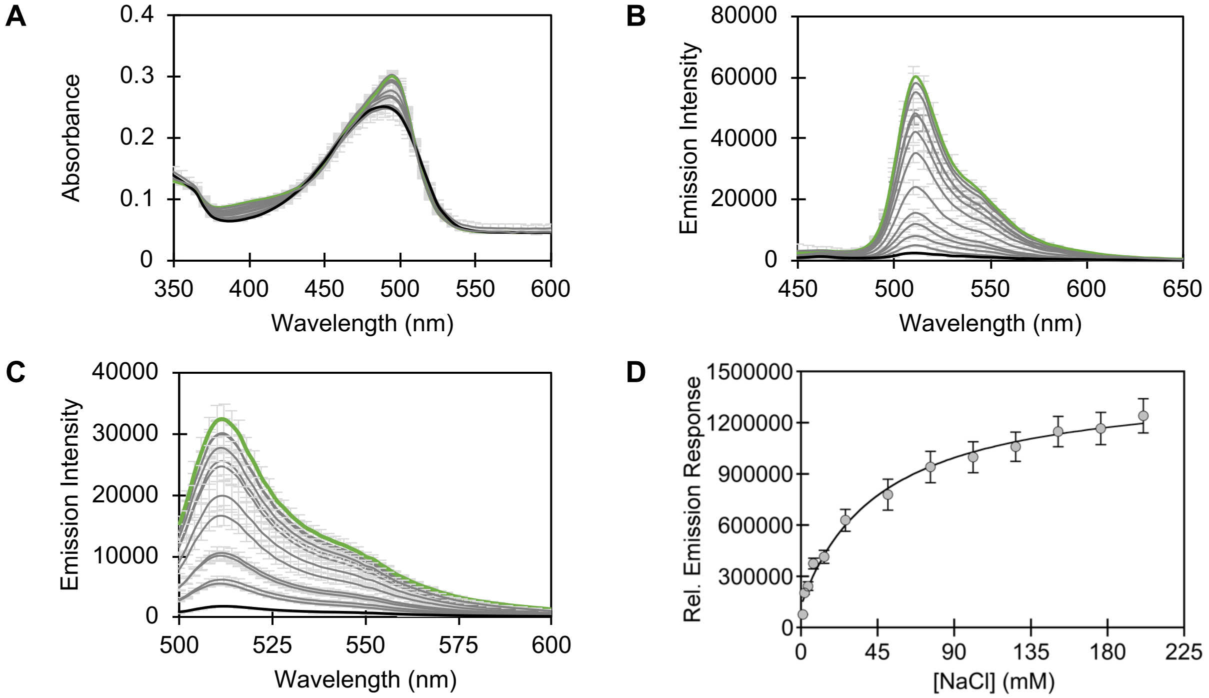


**Figure S8.** Spectroscopic characterization of ~10 µM ChlorON-1-PRO in 50 mM sodium phosphate buffer at pH 7 with 1 (black trace), 2, 4.1, 7.3, 13.5, 26, 51, 76, 101, 126, 151, 176, (gray traces) and 201 (green trace) mM NaCl. (A) Absorbance spectra from 350–600 nm. (B) Emission spectra from 450–650 nm with excitation provided at 400 nm. (C) Emission spectra from 500–650 nm with excitation provided at 485 nm. (D) The integrated emission response from panel C was used to calculate the apparent dissociation constant. The average of four technical replicates across two protein batches with standard deviation is reported.

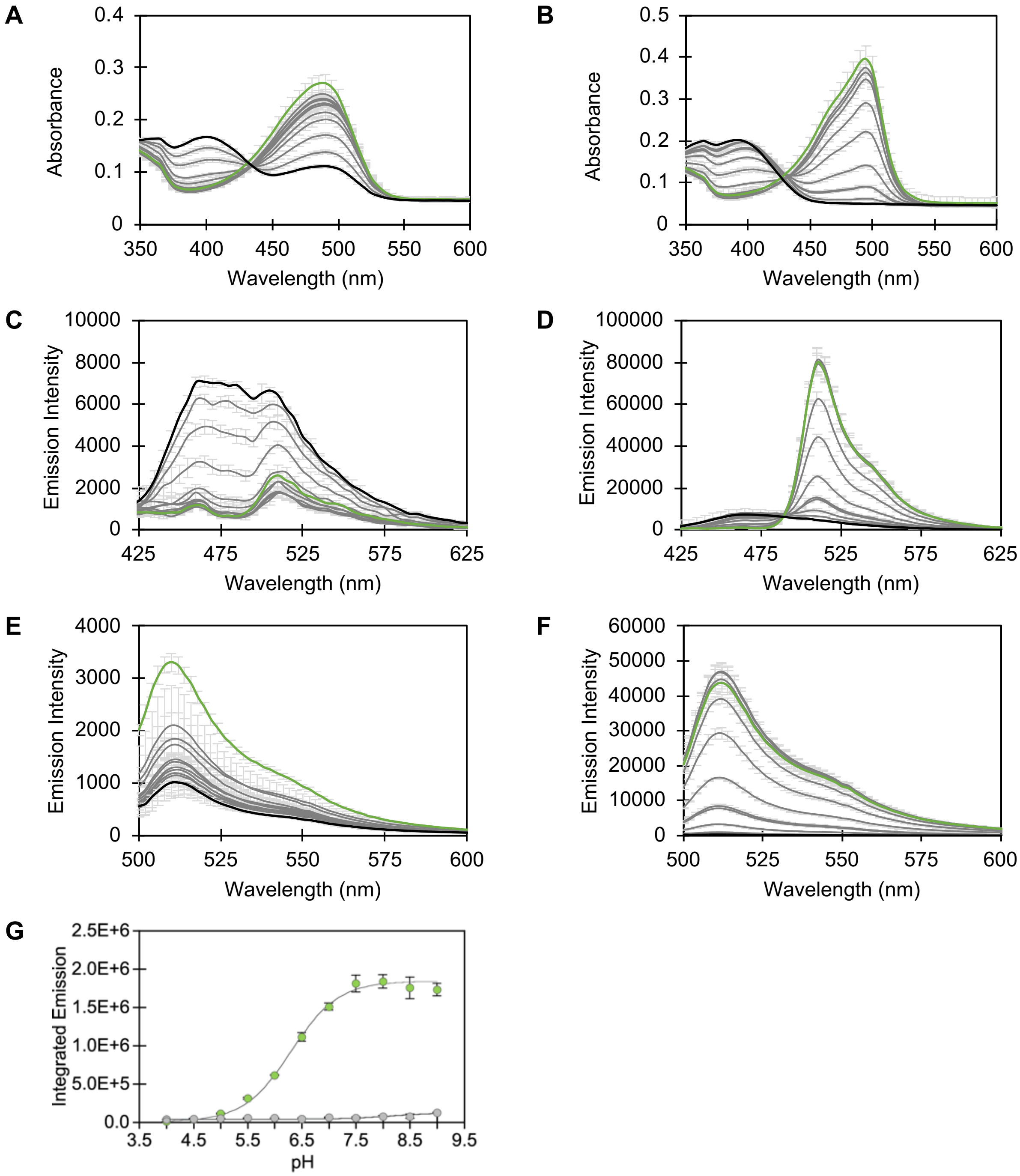


**Figure S9.** Spectroscopic characterization of ~10 μM ChlorON-1-PRO with chloride at varying pH. Absorbance spectra in the presence of (A) 1 mM and (B) 201 mM NaCl. Fluorescence spectra from 450–650 nm in the presence of (C) 1 mM and (D) 201 mM NaCl with excitation provided at 400 nm. Fluorescence spectra from 500–650 nm in the presence of (E) 1 mM and (F) 201 mM NaCl with excitation provided at 485 nm. All spectra were collected in 50 mM sodium acetate buffer at pH 4 (black trace), 4.5, 5, and 5.5 (gray traces) and 50 mM sodium phosphate buffer at pH 6, 6.5, 7, 7.5, 8, 8.5, (gray traces) and 9 (green trace). (G) The integrated emission responses from panels E and F were used to calculate the chromophore p$K_a$. The average of four technical replicates across two protein batches with standard deviation is reported.

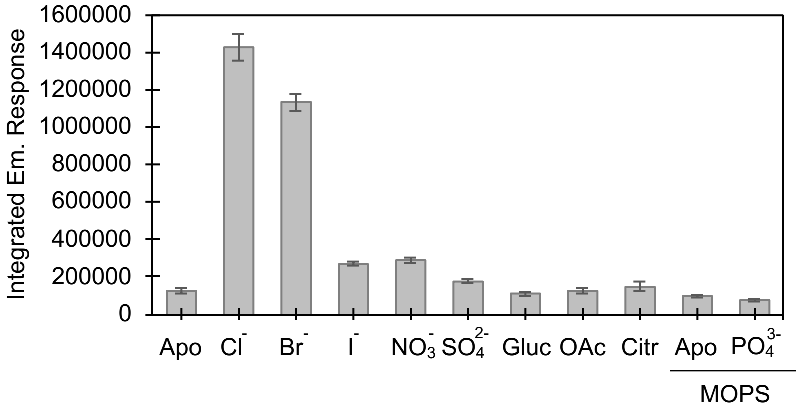


**Figure S10.** Anion selectivity of ~10 μM ChlorON-1-PRO in 50 mM sodium phosphate buffer or 50 mM MOPS buffer at pH 7 with 1 mM (Apo) and 201 mM NaCl ($Cl^-$), 200 mM sodium bromide ($Br^-$), iodide ($I^-$), nitrate ($NO_3^-$), sulfate ($SO_4^{2-}$), gluconate (Gluc), acetate (OAc), citrate (Citr), and phosphate ($PO_4^{3-}$) which is predominantly dihydrogen and hydrogen phosphate at pH 7. Excitation provided at 485 nm, and the emission was collected and integrated from 500–650 nm. The average integrated emission response of four technical replicates across two protein batches with standard deviation is reported (Table S3). Abbreviation: Em, emission.

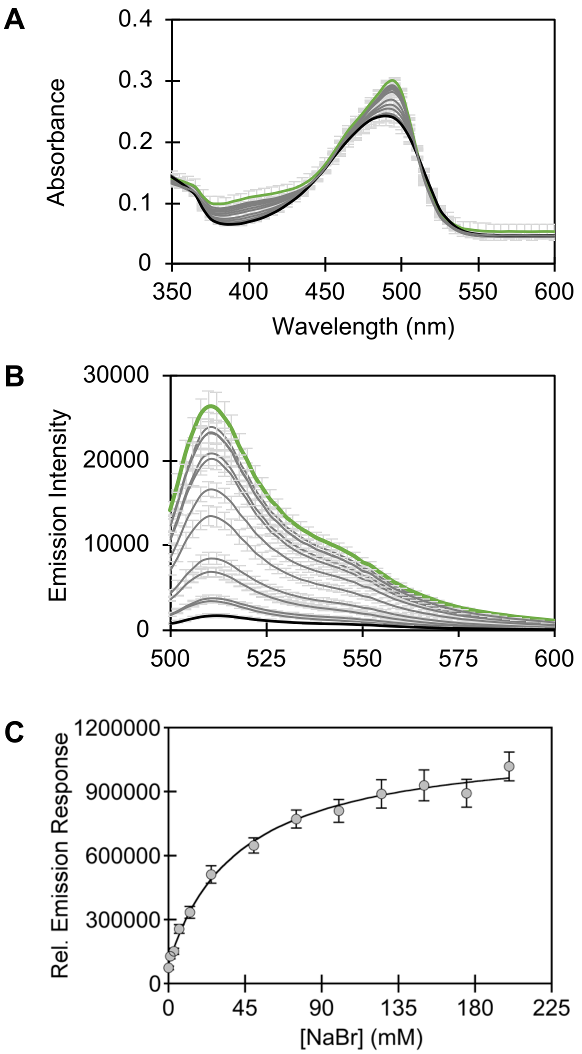


**Figure S11.** Spectroscopic characterization of ~10 μM ChlorON-1-PRO in 50 mM sodium phosphate buffer at pH 7 with 0 (black trace), 1, 3.1, 6.3, 12.5, 25, 50, 75, 100, 125, 150, 175, (gray traces) and 200 (green trace) mM NaBr. (A) Absorbance spectra from 350–600 nm. (B) Emission spectra from 500–650 nm with excitation provided at 485 nm. (C) The integrated emission response from Panel B was used to calculate the apparent dissociation constant. The average of four technical replicates across two protein batches with standard deviation is reported (Table S3).

**Table S3.** Summary for the fluorescence response ($F_f/F_i$) of ChlorON-1-PRO in the presence of 1 mM NaCl ($F_i$) and 201 mM chloride or 200 mM anion ($F_f$) tested with the apparent dissociation constant ($K_d$), when it could be determined. The average of four technical replicates across two protein batches with standard deviation (SD) is reported. Abbreviation: N.D., not determined.

| **Anion** [a] | **$F_f/F_i$ ± SD** | **$K_d$ ± SD (mM)** |
|---|---|---|
| Chloride | 13.9 ± 2.6 | 47.4 ± 7.4 |
| Bromide | 11.6 ± 2.8 | 41.3 ± 2.2 |
| Iodide | 2.21 ± 0.29 | N.D. |
| Nitrate | 2.38 ± 0.21 | N.D. |
| Sulfate | 1.45 ± 0.10 | N.D. |
| Gluconate | no response | N.D. |
| Acetate | no response | N.D. |
| Citrate | no response | N.D. |
| Phosphate [b] | no response | N.D. |

[a] *Determined in 50 mM sodium phosphate buffer at pH 7.*
[b] *Determined in 50 mM MOPS buffer at pH 7.*

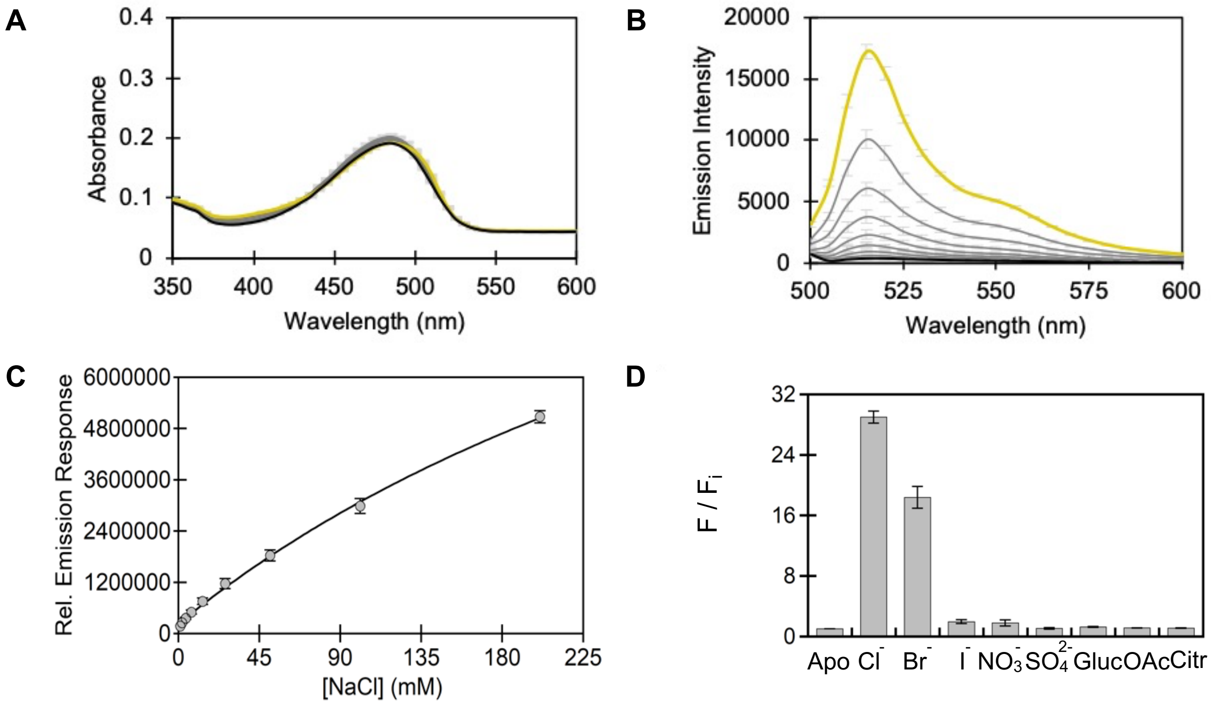


**Figure S12.** Spectroscopic characterization of ~10 μM ChlorON-1 in 50 mM sodium phosphate buffer at pH 7 with 1 (black trace), 3.9, 6.9, 13.3, 25.5, 50, 99, (gray traces) and 197 (yellow trace) mM NaCl. (A) Absorbance spectra from 350–600 nm. (B) Emission spectra from 450–650 nm with excitation provided at 400 nm. (C) Emission spectra from 500–650 nm with excitation provided at 485 nm. (D) The integrated emission response from panel C was used to calculate the apparent dissociation constant. The average of four technical replicates across two protein batches with standard deviation is reported. These data were adapted from Reference 3.

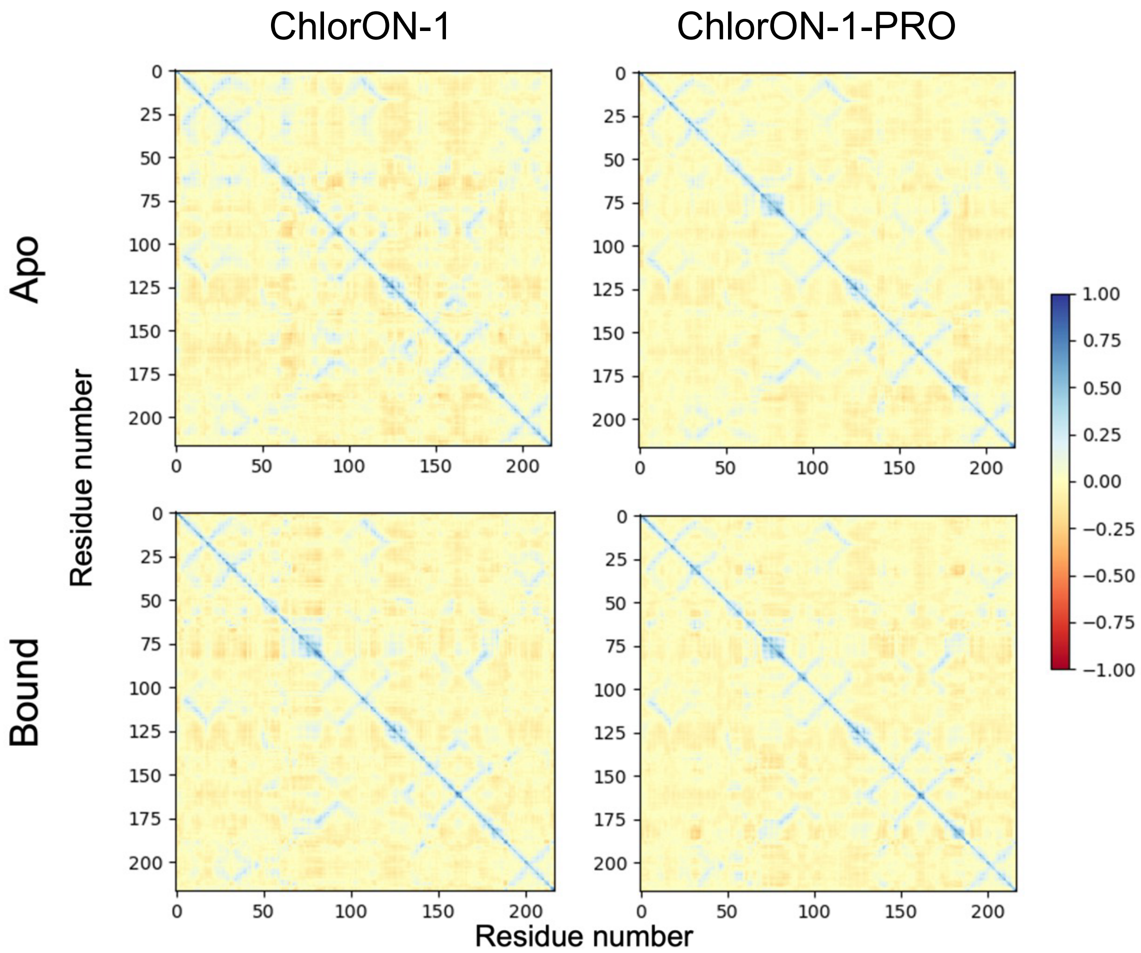


**Figure S13.** Dynamic cross correlation analysis of the apo and chloride bound structures of ChlorON-1 (left) and ChlorON-1-PRO (right) to evaluate the extent of coordinated residue motions. Positively correlated motions are shown in blue and anti-correlated motions are shown in red. Uncorrelated or independent movements are colored in yellow.

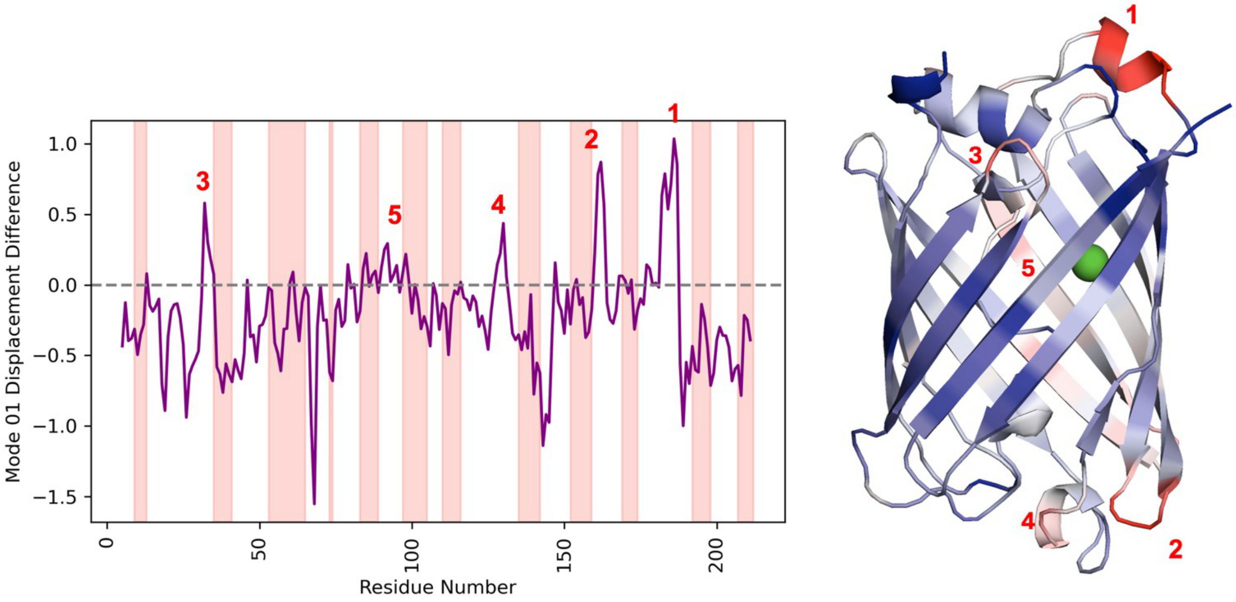


**Figure S14.** Normal mode analysis of ChlorON-1 and ChlorON-1-PRO to examine the global breathing motions of the β-barrel structure in the chloride bound states. Left: The plot illustrates the difference in displacement along mode 01 between the two systems, with more positive values indicating greater motion in ChlorON-1-PRO and negative values indicating greater motion in ChlorON-1. Pink vertical bands highlight residues within 13 Å of the chromophore to include the proximal β-barrel residues. Right: These differences are mapped onto the β-barrel structure where red corresponds to positive values and blue to negative values. Regions with the highest difference are numbered 1–5 and labeled on both the normal mode plot and the β-barrel structure. The bound chloride ion is shown as a green sphere.

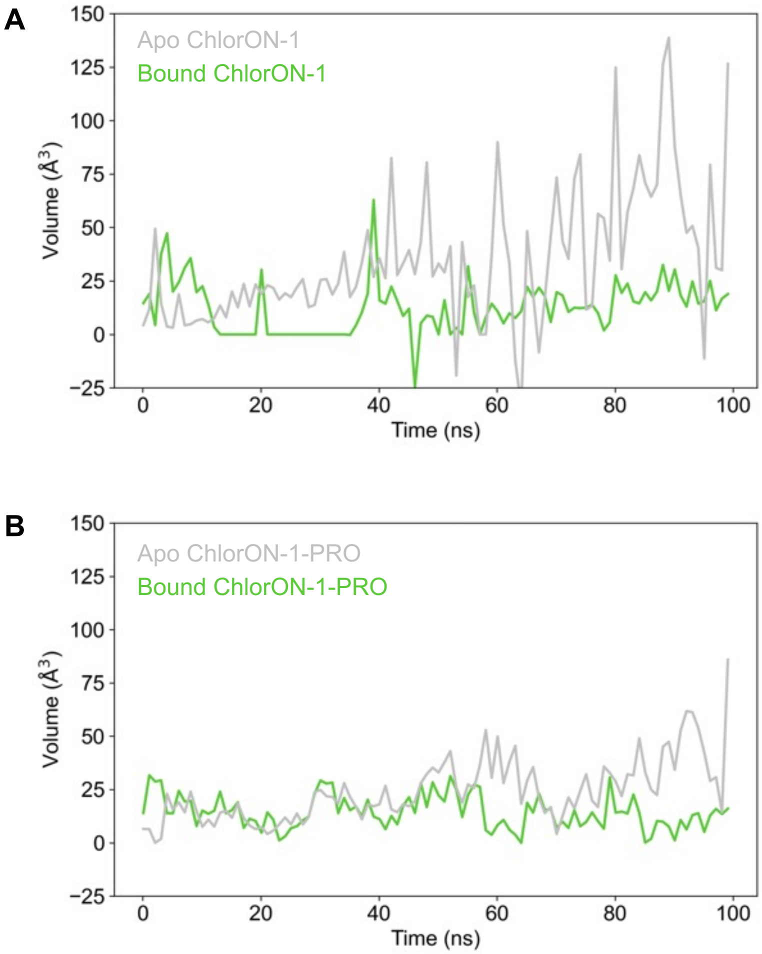


**Figure S15.** Overlay of the change in water occupied volume within 7 Å of the chromophore over time. Water volume was calculated as the difference in volume between individual structure snapshots extracted as Protein Data Bank files with and without water for every 10$^{th}$ frame throughout the simulation. These data are shown for the apo (gray) and bound (green) forms of (A) ChlorON-1 and (B) ChlorON-1-PRO.

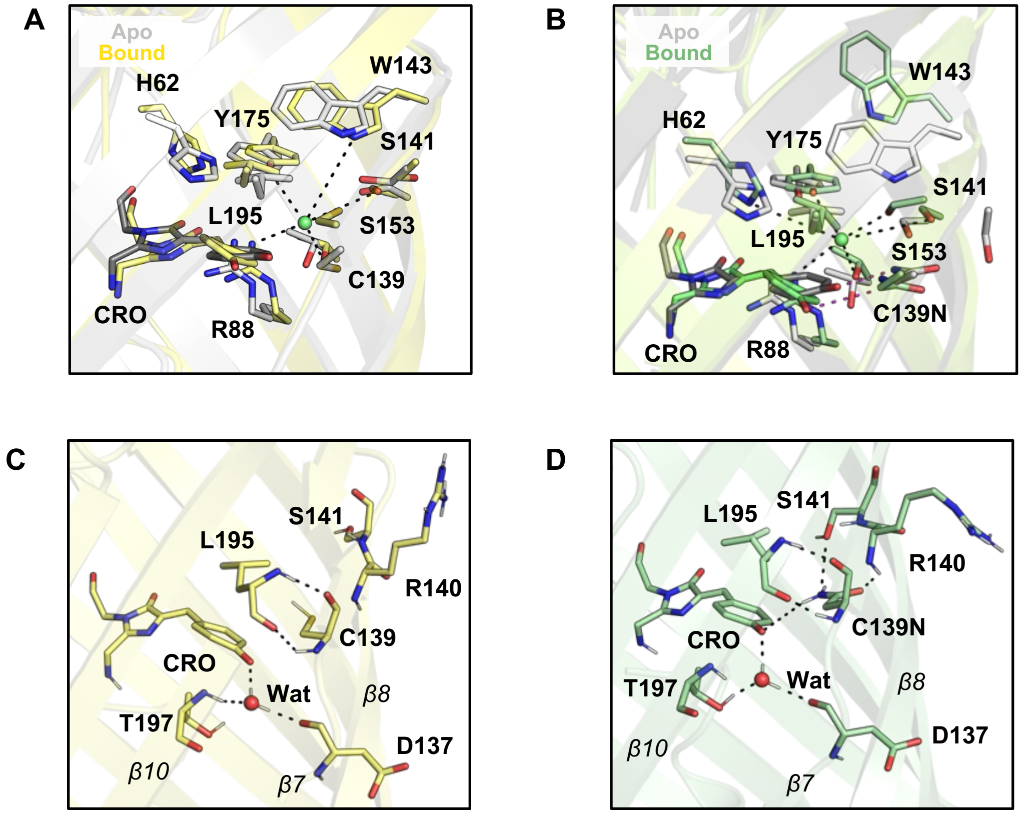


**Figure S16.** Overlay of the molecular dynamics simulations for apo (gray) and chloride bound (yellow or green) (A) ChlorON-1 and (B) ChlorON-1-PRO. Interactions between adjacent residues and chloride and the chromophore are shown as dashed lines. Comparison of the chloride-bound (C) ChlorON-1 and (D) ChlorON-1-PRO structures to illustrate the new hydrogen-bonding interactions between C139N and S141 and R140 in ChlorON-1-PRO. Key β-strands are labeled. Abbreviations: CRO, chromophore; Wat, water.

**Table S4.** Chemical structure of the chromophore (CR2) shown in the schematic with labeled atoms (top). Hydrogen bond analysis for the apo and chloride bound states of ChlorON-1 (middle) and ChlorON-1-PRO (bottom). Occupancies are provided for each interaction between the chromophore and adjacent residues with values greater than 20% observed during the MD simulations.

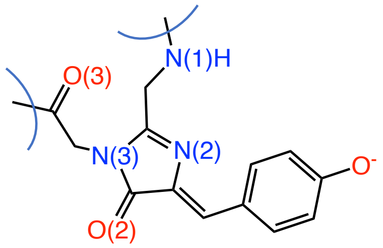


| Apo ChlorON-1 | | | Bound ChlorON-1 | | |
|---|---|---|---|---|---|
| Acceptor | Donor | Occupancy | Acceptor | Donor | Occupancy |
| CR2_60@O3 | TYR_104@OH | 92.0 | CR2_60@O3 | TYR_104@OH | 48.8 |
| | | | CR2_60@O3 | TYR_84@OH | 20.6 |
| CR2_60@O$^-$ | Solvent | 100.0 | CR2_60@O$^-$ | Solvent | 96.0 |
| CR2_60@O2 | ARG_88@NH1 | 58.4 | CR2_60@O2 | ARG_88@NH1 | 66.5 |
| CR2_60@O2 | HIE_62@NE2 | 43.6 | CR2_60@O2 | HIE_62@NE2 | 69.3 |
| CR2_60@O3 | HIE_62@N | 24.9 | | | |
| CR2_60@O2 | ARG_88@NH2 | 84.3 | CR2_60@O2 | ARG_88@NH2 | 71.0 |
| CR2_60@N2 | GLH_210@OE2 | 20.4 | | | |

| Apo ChlorON-1-PRO | | | Bound ChlorON-1-PRO | | |
|---|---|---|---|---|---|
| Acceptor | Donor | Occupancy | Acceptor | Donor | Occupancy |
| CR2_60@O3 | TYR_104@OH | 94.1 | CR2_60@O3 | TYR_104@OH | 92.8 |
| CR2_60@O$^-$ | Solvent | 89.1 | CR2_60@O$^-$ | Solvent | 93.2 |
| CR2_60@O2 | ARG_88@NH1 | 51.0 | CR2_60@O2 | ARG_88@NH1 | 80.7 |
| | | | CR2_60@O2 | HIE_62@NE2 | 70.7 |
| CR2_60@O2 | ARG_88@NH2 | 86.5 | CR2_60@O2 | ARG_88@NH2 | 61.4 |
| CR2_60@N2 | GLH_210@OE2 | 57.1 | CR2_60@N2 | GLH_210@OE2 | 41.5 |
| CR2_60@O$^-$ | ASN_139@ND2 | 79.8 | CR2_60@O$^-$ | ASN_139@ND2 | 32.6 |
| | | | PRO_57@O | CR2_60@N1 | 21.6 |
| CR2_60@O3 | HIE_62@N | 65.5 | | | |

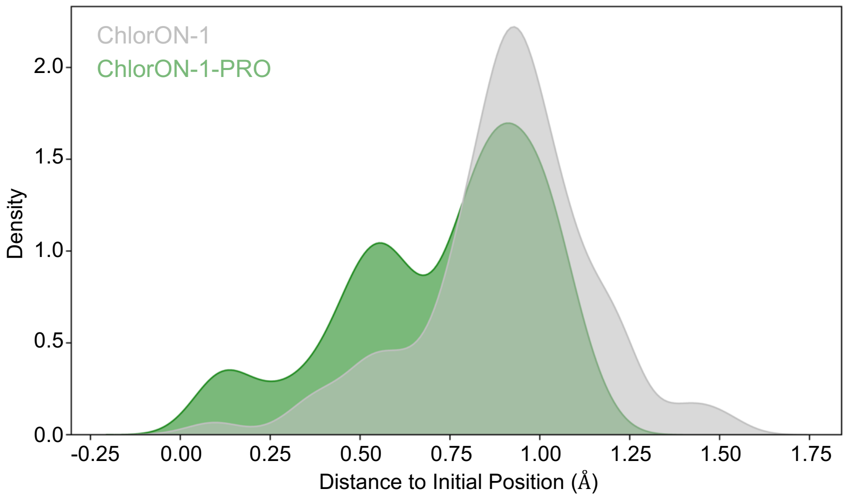


**Figure S17.** Spatial distribution of the bound chloride ion relative to its initial position over the course of the simulation within the binding pocket of ChlorON-1 (gray) and ChlorON-1-PRO (green). Density plots illustrate greater positional variability in ChlorON-1 than ChlorON-1-PRO.

atg cat cat cac cat cac cat ggt acc gag ctc gga tcc ATG GTG TCC AAG GGC GAG GAG GAC AAT ATG

Met His His His His His His Gly Thr Glu Leu Gly Ser Met Val Ser Lys Gly Glu Glu Asp Asn Met

GCC TCT CTG CCA GCC ACC CAC GAG CTG CAC ATC TTC GGC TCT ATC AAC GGC GTG GAC TTT GAT ATG GTG

1 Ala Ser Leu Pro Ala Thr His Glu Leu His Ile Phe Gly Ser Ile Asn Gly Val Asp Phe Asp Met Val

GGA CAG GGA ACC GGA AAC CCA AAT GAC GGC TAC GAG GAG CTG AAT CTG AAG TCT ACA AAG GGC GAT CTG

24 Gly Gln Gly Thr Gly Asn Pro Asn Asp Gly Tyr Glu Glu Leu Asn Leu Lys Ser Thr Lys Gly Asp Leu

CAG TTC AGC CCT TGG ATT CTG GTG CCA CAC ATC GGC TAT GGC TTT CAC CAG TAT CTG CCC TAC CCT GAC

47 Gln Phe Ser Pro Trp Ile Leu Val Pro His Ile Gly Tyr Gly Phe His Gln Tyr Leu Pro Tyr Pro Asp

GGC ATG TCT CCT TTC CAG GCC GCC ATG GTG GAT GGC AGC GGC TAC CAG GTG CAC AGG ACA ATG CAG TTT

70 Gly Met Ser Pro Phe Gln Ala Ala Met Val Asp Gly Ser Gly Tyr Gln Val His Arg Thr Met Gln Phe

GAG GAC GGC GCC TCC CTG ACC GTG AAC TAC CGC TAT ACA TAC GAG GGC TCT CAC ATC AAG GGA GAG GCA

93 Glu Asp Gly Ala Ser Leu Thr Val Asn Tyr Arg Tyr Thr Tyr Glu Gly Ser His Ile Lys Gly Glu Ala

CAG GTG AAG GGA ACC GGA TTC CCA GCA GAT GGA CCC GTG ATG ACC AAC AGC CTG ACA GCA GCA GAC TGG

116 Gln Val Lys Gly Thr Gly Phe Pro Ala Asp Gly Pro Val Met Thr Asn Ser Leu Thr Ala Ala Asp Trp

TGC CGG TCC AAG TGG ACA TAT CCC AAT GAT AAG ACC ATC ATC AGC ACC TTC AAG TGG TCC TAT ACC ACA

139 Asn Arg Ser Lys Trp Thr Tyr Pro Asn Asp Lys Thr Ile Ile Ser Thr Phe Lys Trp Ser Tyr Thr Thr

GGC AAC GGC AAG CGG TAC AGA AGC ACC GCC CGG ACC ACA TAT ACA TTT GCC AAG CCC ATG GCC GCC AAC

162 Gly Asn Gly Lys Arg Tyr Arg Ser Thr Ala Arg Thr Thr Tyr Thr Phe Ala Lys Pro Met Ala Ala Asn

TAT CTG AAG AAT CAG CCT ATG TAC GTG TTC CTG AAG ACC GAG CTG AAG CAC TCC AAG ACA GAG CTG AAT

185 Tyr Leu Lys Asn Gln Pro Met Tyr Val Phe Leu Lys Thr Glu Leu Lys His Ser Lys Thr Glu Leu Asn

TTC AAG GAG TGG CAG AAG GCC TTT ACC GAC GTG ATG GGC ATG GAT GAG CTG TAC AAG tga gaa ttc

208 Phe Lys Glu Trp Gln Lys Ala Phe Thr Asp Val Met Gly Met Asp Glu Leu Tyr Lys * Glu Phe

**Figure S18.** The nucleotide and amino acid sequences for ChlorON-1-PRO in the pcDNA3.1(+)-N-6His vector (black text) with an N-terminal polyhistidine tag (highlighted in cyan). The ChlorON-1-PRO sequence (green text) is cloned between the BamHI and EcoRI restriction sites (highlighted in gray) with the K143W and R195L mutations (blue text) and the C139N mutation (highlighted in yellow). The stop codon is shown as an asterisk (*).

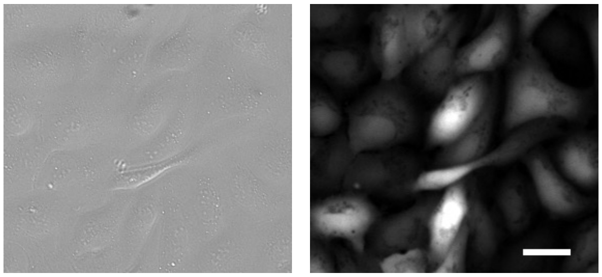

**Figure S19.** Representative differential interference contrast (DIC, left) and fluorescence images (right) of U-2 OS cells stably expressing ChlorON-1-PRO in 137 mM NaCl buffer. Scale bar = 30 µm.

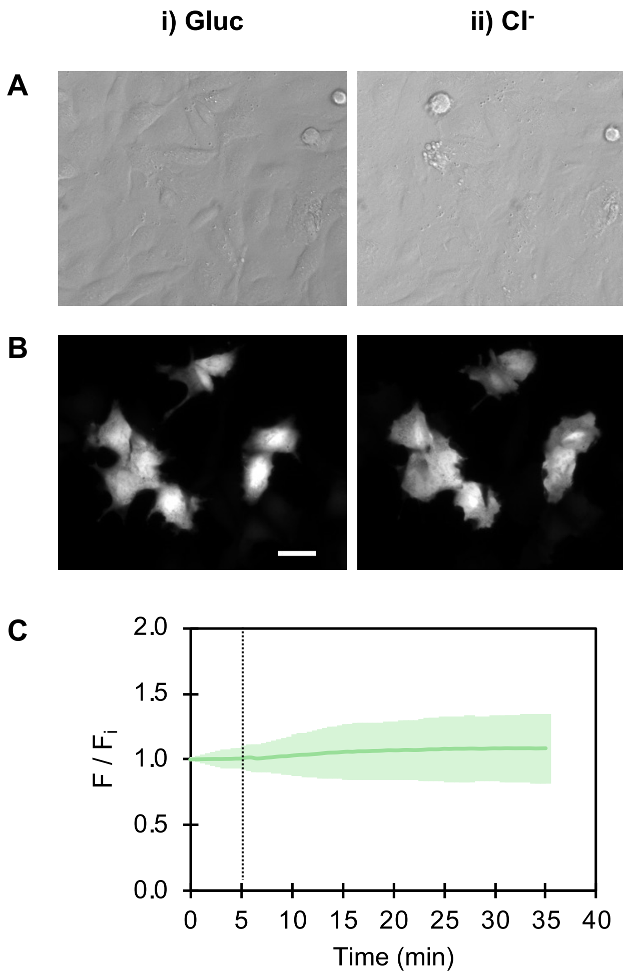


**Figure S20.** Representative fluorescence images of U-2 OS cells transiently expressing mNeonGreen. (A) DIC and (B) fluorescence images of cells in (i) 137 mM gluconate buffer (Gluc, $F_i$) at $t$ = 0 min and (ii) 68.5 mM Gluc/68.5 mM NaCl buffer ($Cl^-$, F) at $t$ = 35 min. Scale bar = 30 μm. (C) Time course of the mNeonGreen emission response ($F/F_i$) over time ($n$ = 78 regions of interest, ROI). Dashed line represents the start of the $Cl^-$ buffer perfusion. The average emission response ($F/F_i$) of three different fields from one biological replicate with standard deviation is reported.

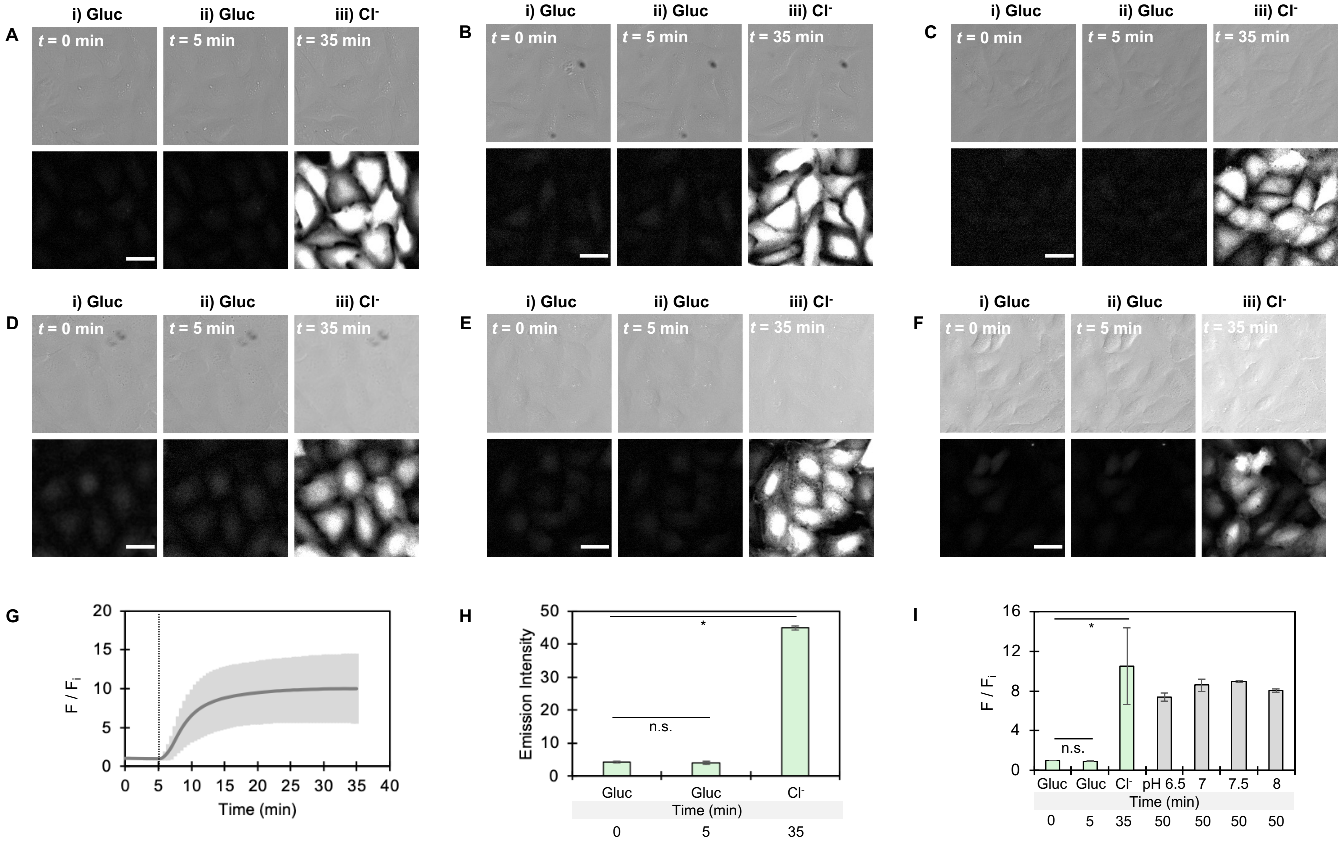


**Figure S21.** Representative DIC and fluorescence images of U-2 OS cells stably expressing ChlorON-1-PRO treated with DMSO. Images from six biological replicates are shown in panels A–F of cells in the presence of 137 mM NaGluc buffer (Gluc) supplemented with DMSO at (i) $t$ = 0 min ($F_i$) and (ii) $t$ = 5 min (F) followed by perfusion with (iii) 68.5 mM Gluc/68.5 mM NaCl buffer ($Cl^-$) supplemented with 0.2% DMSO at $t$ = 35 min (F). Scale bar = 30 µm. (G) Time course of the ChlorON-1-PRO turn-on fluorescence response ($F/F_i$) with DMSO ($n$ = 4003 ROIs). Dashed line represents the start of the perfusion with $Cl^-$ buffer. Bar graphs of the average (H) emissions and (I) turn-on emission responses ($F/F_i$) for each condition listed (green bars). The ChlorON-1-PRO pH clamping data with chloride from Figure S27–S30 (gray bars) are also shown. Statistical significances were determined using the average emissions and emission responses ($F/F_i$) at $t$ = 5 min and $t$ = 35 min for six biological replicates ($N$ = 6) relative to $F_i$ (* = $p$-value < 0.05; n.s. = not significant). Values are reported as the average ± standard deviation of three fields from each biological replicate.

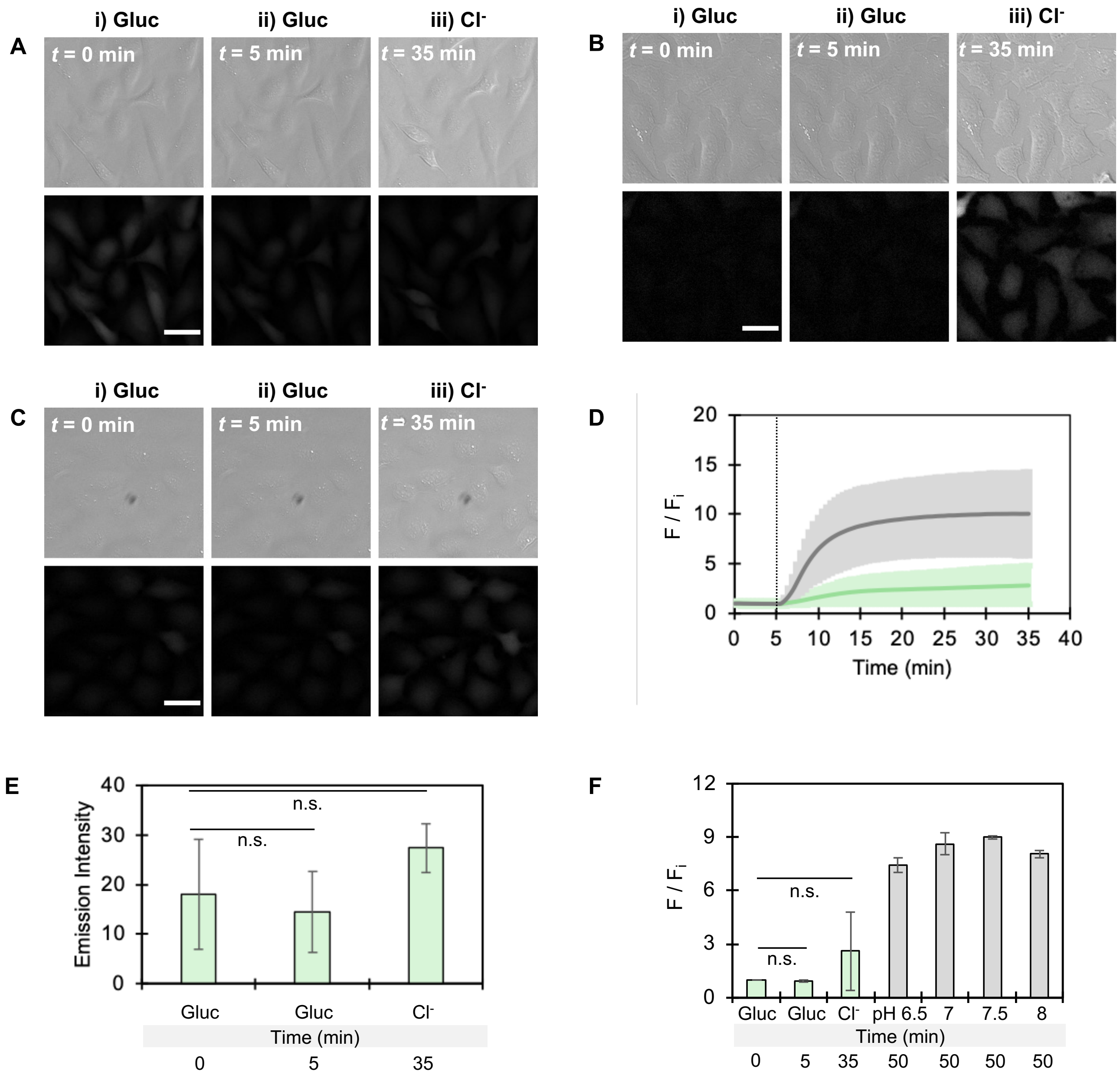


**Figure S22.** Representative DIC and fluorescence images of U-2 OS cells stably expressing ChlorON-1-PRO treated with IAA-94. Images from three biological replicates are shown in panels A–C of cells in the presence of 137 mM NaGluc buffer (Gluc) supplemented with 100 μM IAA-94 at (i) $t$ = 0 min ($F_i$) and (ii) $t$ = 5 min (F) followed by perfusion with (iii) 68.5 mM Gluc/68.5 mM NaCl buffer ($Cl^-$) supplemented with 100 μM IAA-94 at $t$ = 35 min (F). Scale bar = 30 μm. (D) Time course of the ChlorON-1-PRO turn-on fluorescence response ($F/F_i$) with the DMSO vehicle control (gray trace) from Figure S21 and 100 μM IAA-94 (green trace) ($n$ = 3094 ROIs). Dashed line represents the start of the $Cl^-$ buffer perfusion. Bar graphs of the average (E) emissions and (F) turn-on emission responses ($F/F_i$) for each condition listed (green bars). The ChlorON-1-PRO pH clamping data with chloride from Figure S27–S30 (gray bars) are also shown. Statistical significances were determined using the average emissions and emission responses ($F/F_i$) at $t$ = 5 min and $t$ = 35 min for three biological replicates ($N$ = 3) relative to $F_i$ (n.s. = not significant, $p$-value > 0.05). Values are reported as the average ± standard deviation of three fields from each biological replicate.

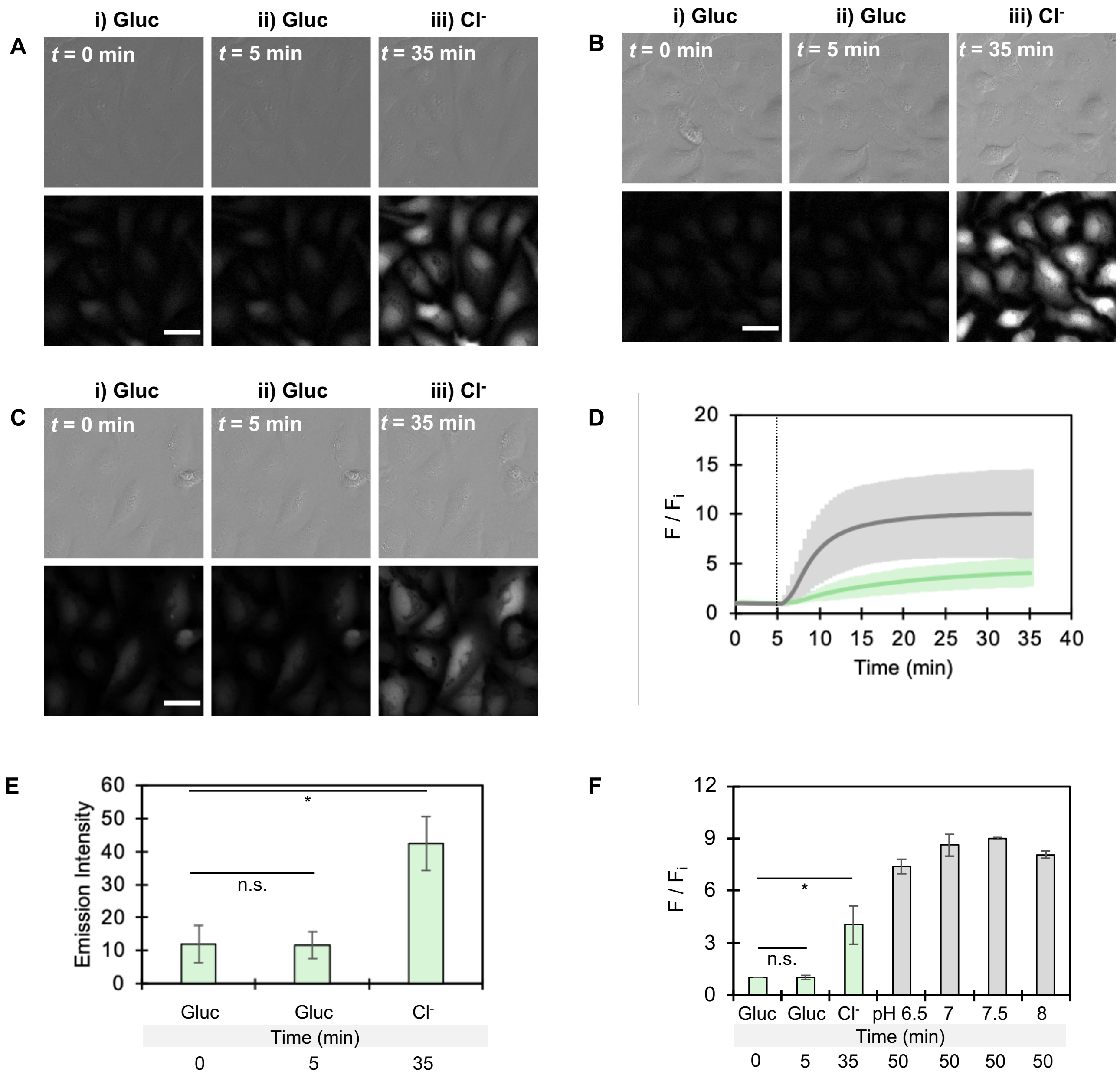


**Figure S23.** Representative DIC and fluorescence images of U-2 OS cells stably expressing ChlorON-1-PRO treated with CaCCinh-A01. Images from three biological replicates are shown in panels A–C of cells in the presence of 137 mM NaGluc buffer (Gluc) supplemented with 20 μM CaCCinh-A01 at (i) $t$ = 0 min ($F_i$) and (ii) $t$ = 5 min (F) followed by perfusion with (iii) 68.5 mM Gluc/68.5 mM NaCl buffer ($Cl^-$) supplemented with 20 μM CaCCinh-A01 at $t$ = 35 min (F). Scale bar = 30 μm. (D) Time course of the ChlorON-1-PRO turn-on fluorescence response ($F/F_i$) with the DMSO vehicle control (gray trace) from Figure S21 and 20 μM CaCCinh-A01 treatment (green trace) ($n$ = 2733 ROIs). Dashed line represents the start of the $Cl^-$ buffer perfusion. Bar graphs of the average (E) emissions and (F) turn-on emission responses ($F/F_i$) for each condition listed (green bars). The ChlorON-1-PRO pH clamping data with chloride from Figure S27–S30 (gray bars) are also shown. Statistical significances were determined using the average emissions and emission responses ($F/F_i$) at $t$ = 5 min and $t$ = 35 min for three biological replicates ($N$ = 3) relative to $F_i$ (* = $p$-value < 0.05; n.s. = not significant). Values are reported as the average ± standard deviation of three fields from each biological replicate.

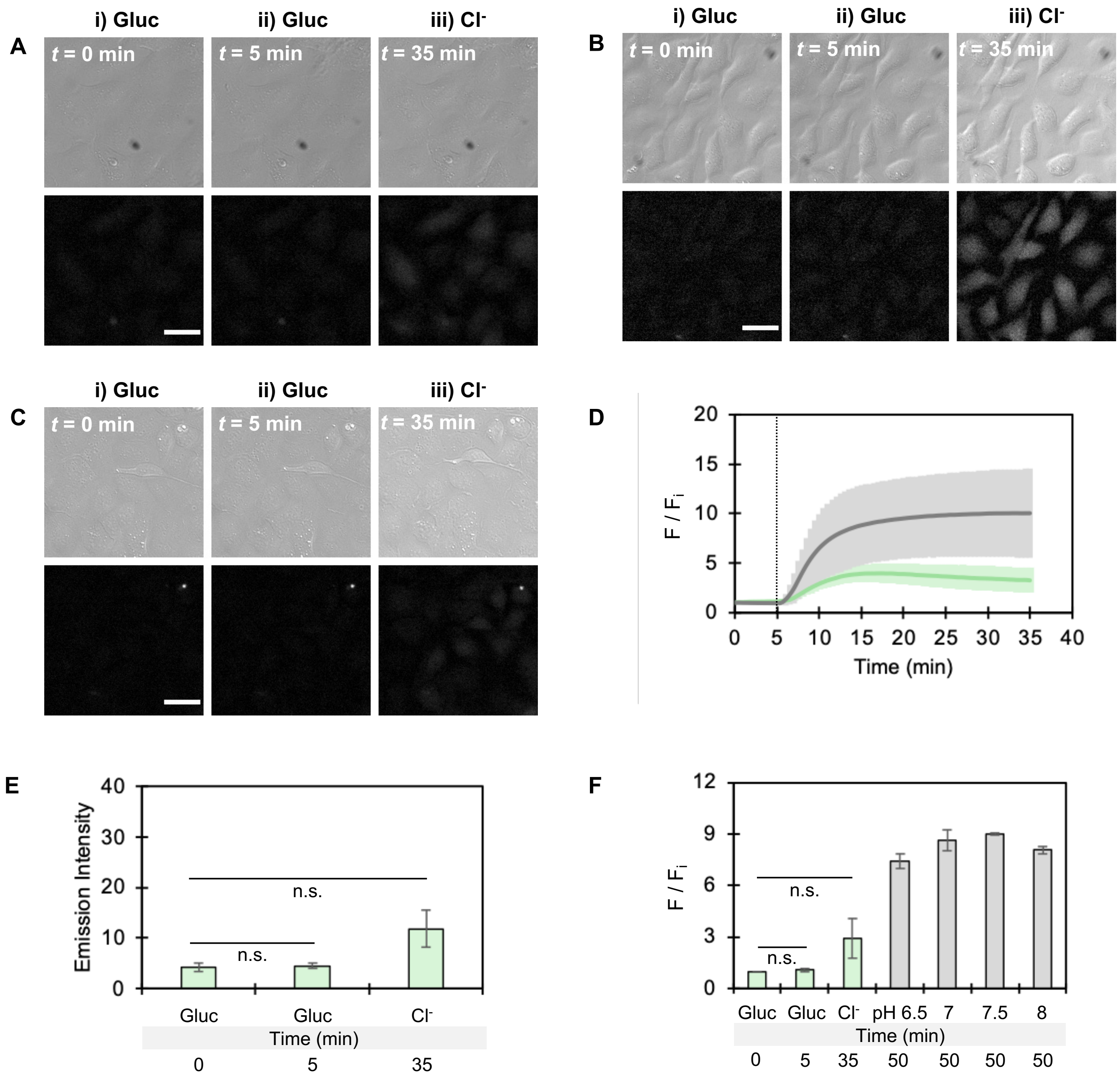


**Figure S24.** Representative DIC and fluorescence images of U-2 OS cells stably expressing ChlorON-1-PRO treated with BAPTA-AM ester. Images from three biological replicates are shown in panels A–C of cells in the presence of 137 mM NaGluc buffer (Gluc) supplemented with 20 μM BAPTA-AM ester at (i) $t$ = 0 min ($F_i$) and (ii) $t$ = 5 min (F) followed by perfusion with (iii) 68.5 mM Gluc/68.5 mM NaCl buffer ($Cl^-$) supplemented with 20 μM BAPTA-AM ester at $t$ = 35 min (F). Scale bar = 30 μm. (D) Time course of the ChlorON-1-PRO turn-on fluorescence response ($F/F_i$) with the DMSO vehicle control (gray trace) from Figure S21 and 20 μM BAPTA-AM treatment (green trace) ($n$ = 2741 ROIs). Dashed line represents the start of the $Cl^-$ buffer perfusion. Bar graphs of the average (E) emissions and (F) turn-on emission responses ($F/F_i$) for each condition listed (green bars). The ChlorON-1-PRO pH clamping data with chloride from Figure S27–S30 (gray bars) are also shown. Statistical significances were determined using the average emissions and emission responses ($F/F_i$) at $t$ = 5 min and $t$ = 35 min for three biological replicates ($N$ = 3) relative to $F_i$ (n.s. = not significant, $p$-value > 0.05). Values are reported as the average ± standard deviation of three fields from each biological replicate. Abbreviation: AM, acetoxymethyl.

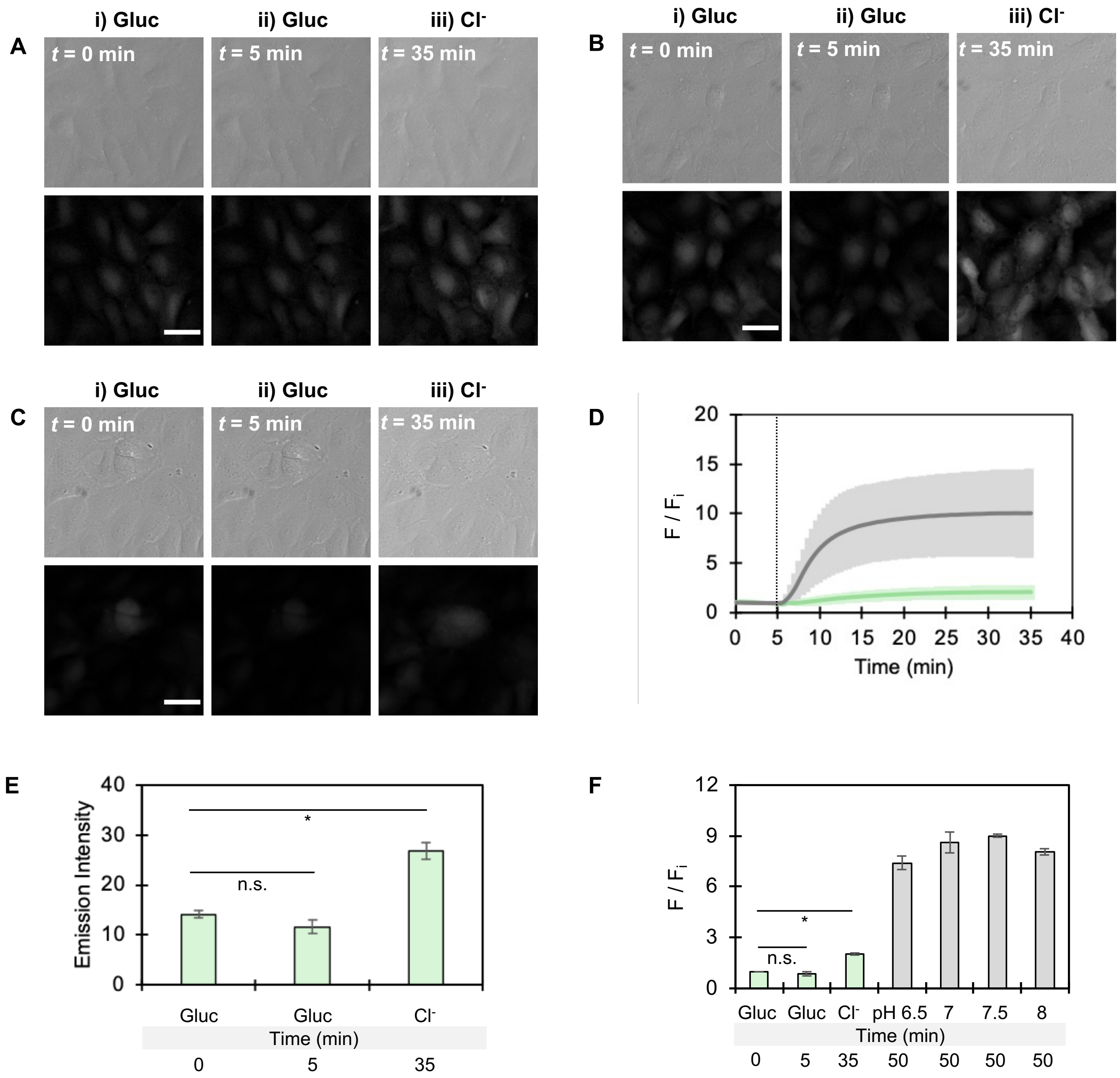


**Figure S25.** Representative DIC and fluorescence images of U-2 OS cells stably expressing ChlorON-1-PRO treated with niclosamide. Images from three biological replicates are shown in panels A–C of cells in the presence of 137 mM NaGluc buffer (Gluc) supplemented with 5 μM niclosamide at (i) $t$ = 0 min ($F_i$) and (ii) $t$ = 5 min (F) followed by perfusion with (iii) 68.5 mM Gluc/68.5 mM NaCl buffer ($Cl^-$) supplemented with 5 μM niclosamide at $t$ = 35 min (F). Scale bar = 30 μm. (D) Time course of the ChlorON-1-PRO turn-on fluorescence response ($F/F_i$) with the DMSO vehicle control (gray trace) from Figure S21 and 5 μM niclosamide treatment (green trace) ($n$ = 2898 ROIs). Dashed line represents the start of the $Cl^-$ buffer perfusion. Bar graphs of the average (E) emissions and (F) turn-on emission responses ($F/F_i$) for each condition listed (green bars). The ChlorON-1-PRO pH clamping data with chloride from Figure S27–S30 (gray bars) are also shown. Statistical significances were determined using the average emissions and emission responses ($F/F_i$) at $t$ = 5 min and $t$ = 35 min for three biological replicates ($N$ = 3) relative to $F_i$ (* = $p$-value < 0.05; n.s. = not significant). Values are reported as the average ± standard deviation of three fields from each biological replicate.

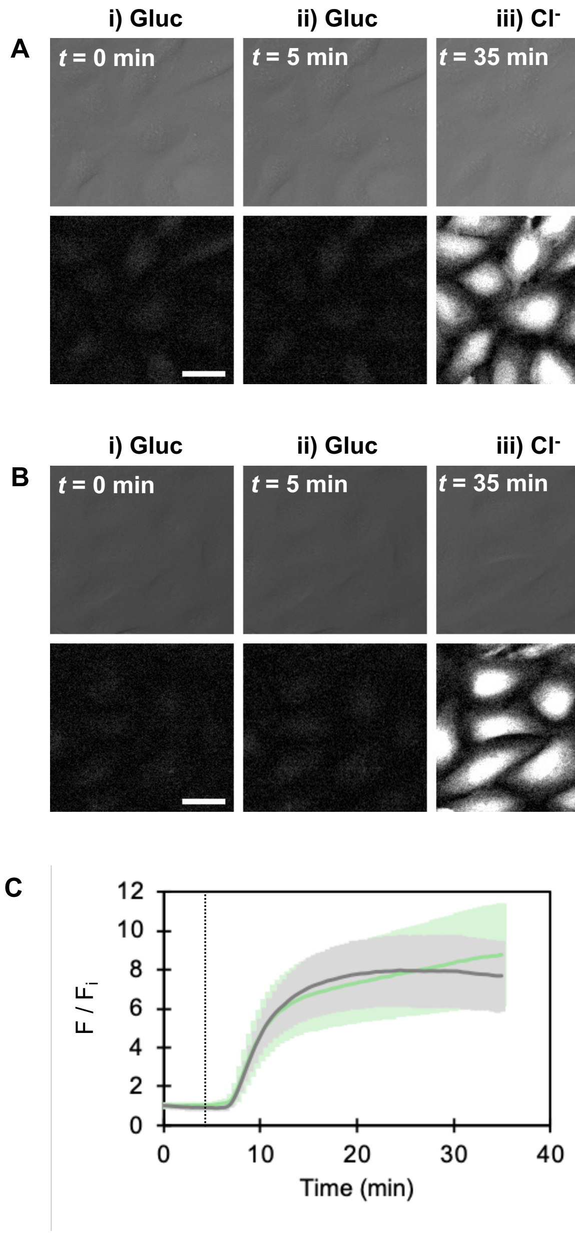


**Figure S26.** Representative DIC (top row) and fluorescence images (bottom row) of U-2 OS cells stably expressing ChlorON-1-PRO treated with (A) 0.1% DMSO (vehicle control) or (B) 10 µM $E_{act}$. Images are shown in the presence of 137 mM NaGluc buffer (Gluc) supplemented with DMSO or 10 µM $E_{act}$ at (i) $t$ = 0 min ($F_i$) and (ii) $t$ = 5 min (F) followed by perfusion with (iii) 68.5 mM Gluc/68.5 mM NaCl buffer ($Cl^-$) supplemented with DMSO or $E_{act}$ at $t$ = 35 min (F). Scale bar = 30 µm. (C) Time course of the ChlorON-1-PRO turn-on fluorescence response ($F/F_i$) with the DMSO vehicle control (gray trace) and 10 µM $E_{act}$ treatment (green trace). Dashed line represents the start of the $Cl^-$ buffer perfusion. Values are reported as the average ± standard deviation of three fields from each biological replicate.

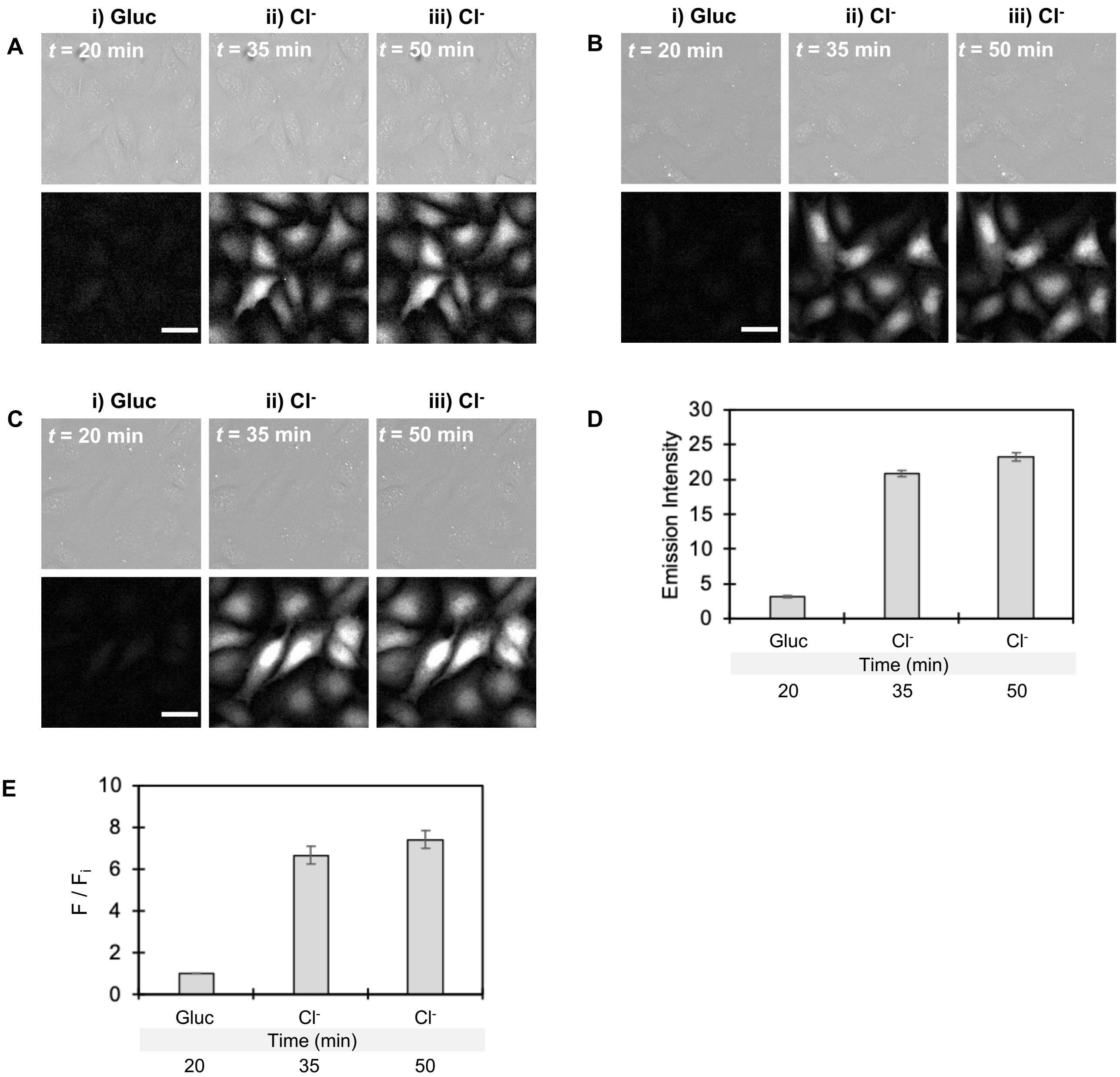


**Figure S27.** Representative DIC (top row) and fluorescence images (bottom row) of U-2 OS cells stably expressing ChlorON-1-PRO clamped at pH 6.5. Images from three biological replicates are shown in panels A–C of cells in (i) 140 mM Gluc clamping buffer at pH 6.5 containing 5 μM nigericin and 5 μM valinomycin at *t* = 20 min ($F_i$) followed by 70 mM Gluc/70 mM $Cl^-$ ($Cl^-$) clamping buffer at pH 6.5 containing 5 μM nigericin and 5 μM valinomycin at (ii) *t* = 35 min (F) and (iii) *t* = 50 min (F). Scale bar = 30 μm. Bar graphs of the ChlorON-1-PRO (D) emissions and (E) turn-on fluorescence responses ($F/F_i$) for each treatment. Values are reported as the average ± standard deviation of three fields from three biological replicates.

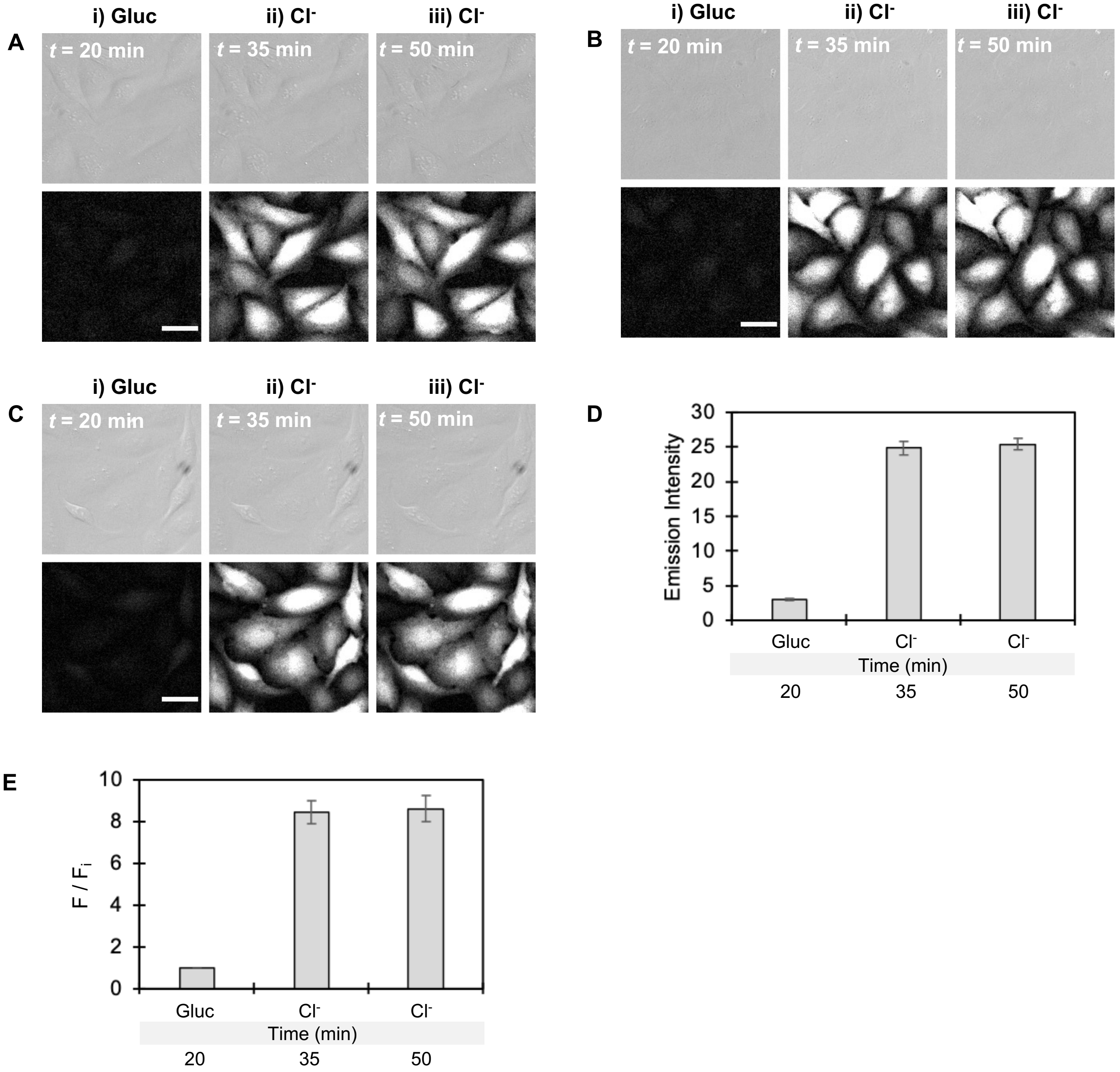


**Figure S28.** Representative DIC (top row) and fluorescence images (bottom row) of U-2 OS cells stably expressing ChlorON-1-PRO clamped at pH 7. Images from three biological replicates are shown in panels A–C of cells in (i) 140 mM Gluc clamping buffer at pH 7 containing 5 μM nigericin and 5 μM valinomycin at *t* = 20 min ($F_i$) followed by 70 mM Gluc/70 mM $Cl^-$ ($Cl^-$) clamping buffer at pH 7 containing 5 μM nigericin and 5 μM valinomycin at (ii) *t* = 35 min (F) and (iii) *t* = 50 min (F). Scale bar = 30 μm. Bar graphs of the ChlorON-1-PRO (D) emissions and (E) turn-on fluorescence responses ($F/F_i$) for each treatment. Values are reported as the average ± standard deviation of three fields from three biological replicates.

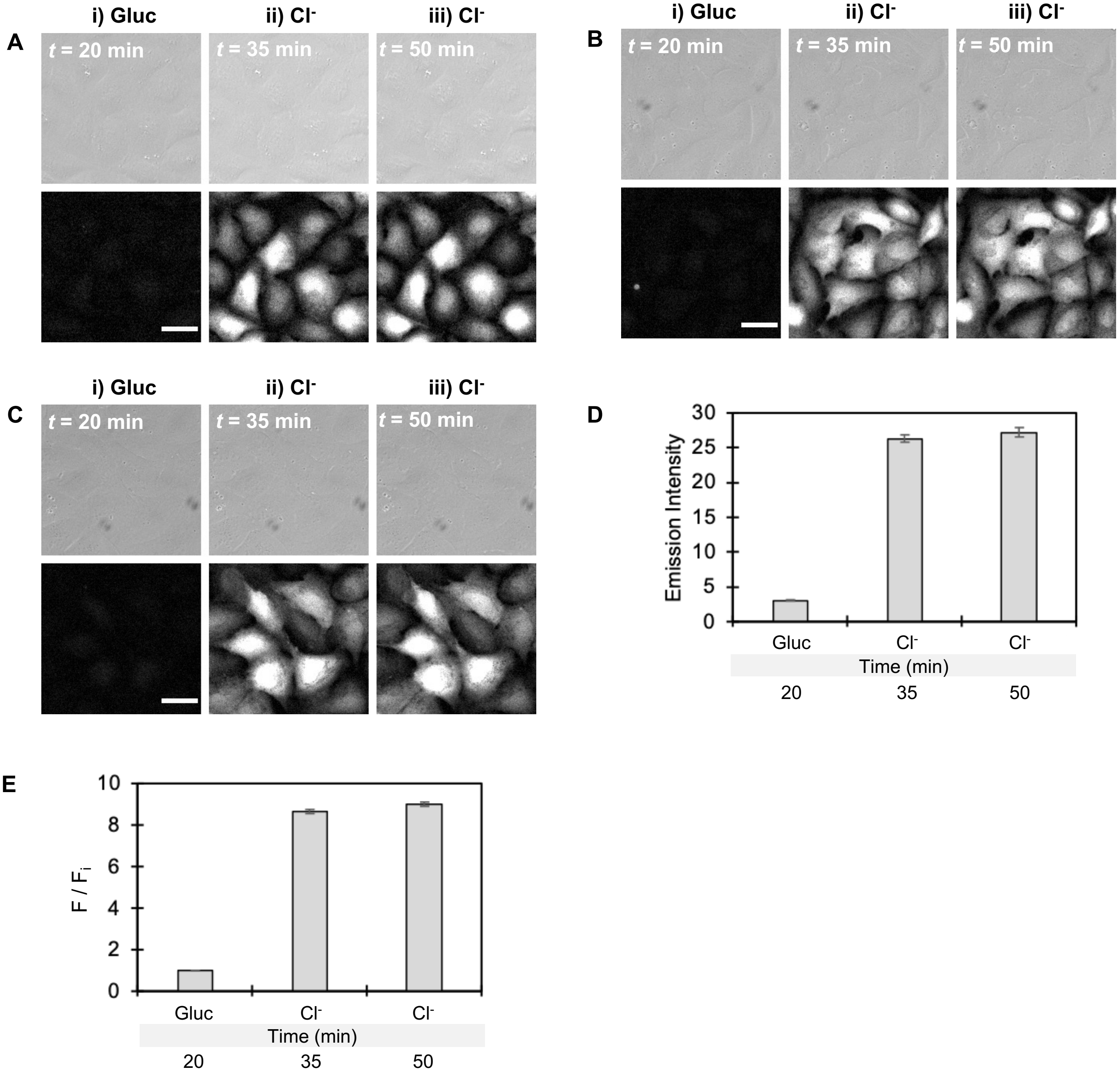


**Figure S29.** Representative DIC (top row) and fluorescence images (bottom row) of U-2 OS cells stably expressing ChlorON-1-PRO clamped at pH 7.5. Images from three biological replicates are shown in panels A–C of cells in (i) 140 mM Gluc clamping buffer at pH 7.5 containing 5 μM nigericin and 5 μM valinomycin at *t* = 20 min ($F_i$) followed by 70 mM Gluc/70 mM $Cl^-$ ($Cl^-$) clamping buffer at pH 7.5 containing 5 μM nigericin and 5 μM valinomycin at (ii) *t* = 35 min (F) and (iii) *t* = 50 min (F). Scale bar = 30 μm. Bar graphs of the ChlorON-1-PRO (D) emissions and (E) turn-on fluorescence responses ($F/F_i$) for each treatment. Values are reported as the average ± standard deviation of three fields from three biological replicates.

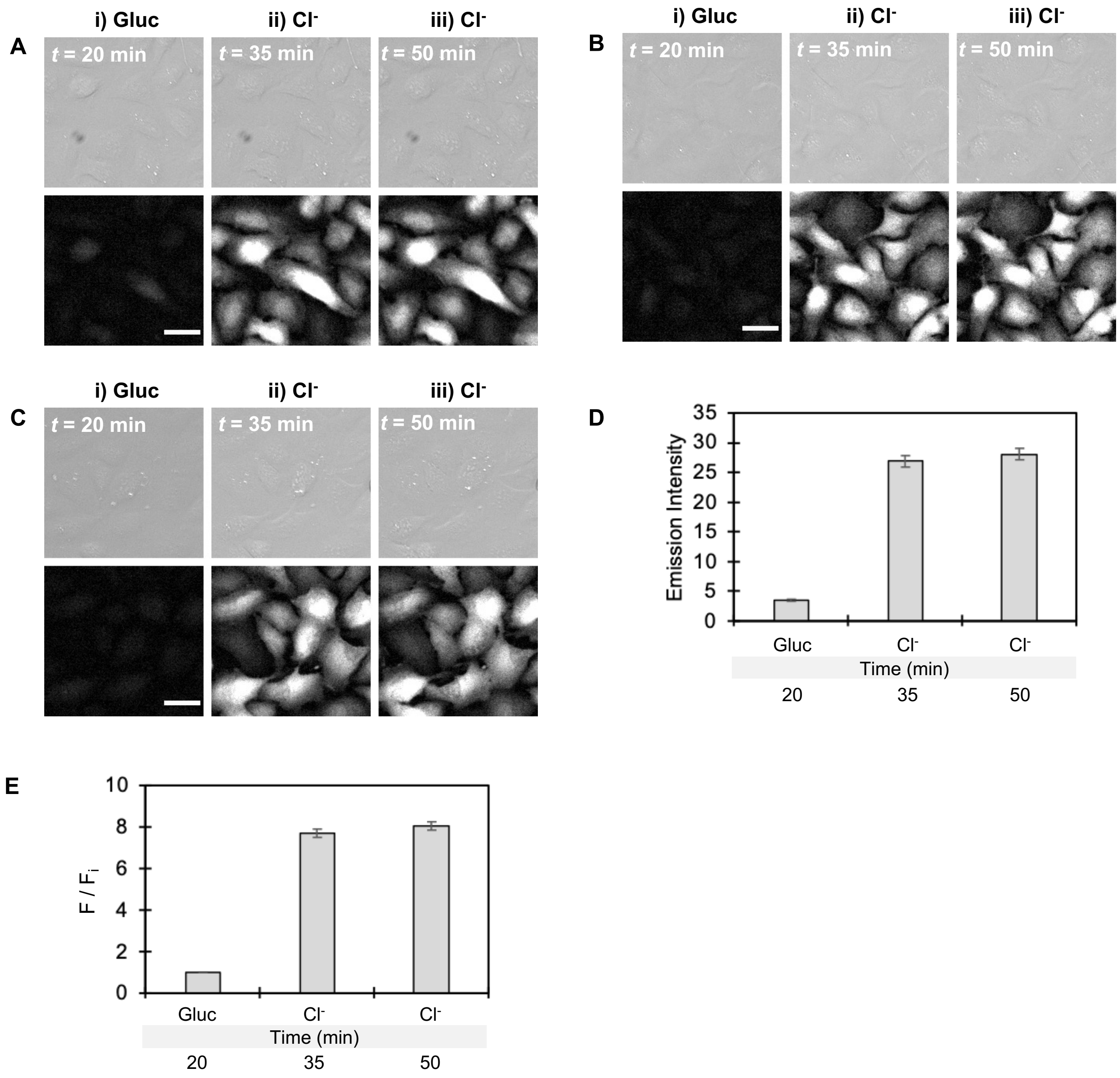


**Figure S30.** Representative DIC (top row) and fluorescence images (bottom row) of U-2 OS cells stably expressing ChlorON-1-PRO clamped at pH 8. Images from three biological replicates are shown in panels A–C of cells in (i) 140 mM Gluc clamping buffer at pH 8 containing 5 μM nigericin and 5 μM valinomycin at *t* = 20 min ($F_i$) followed by 70 mM Gluc/70 mM $Cl^-$ ($Cl^-$) clamping buffer at pH 8 containing 5 μM nigericin and 5 μM valinomycin at (ii) *t* = 35 min (F) and (iii) *t* = 50 min (F). Scale bar = 30 μm. Bar graphs of the ChlorON-1-PRO (D) emissions and (E) turn-on fluorescence responses ($F/F_i$) for each treatment. Values are reported as the average ± standard deviation of three fields from three biological replicates.

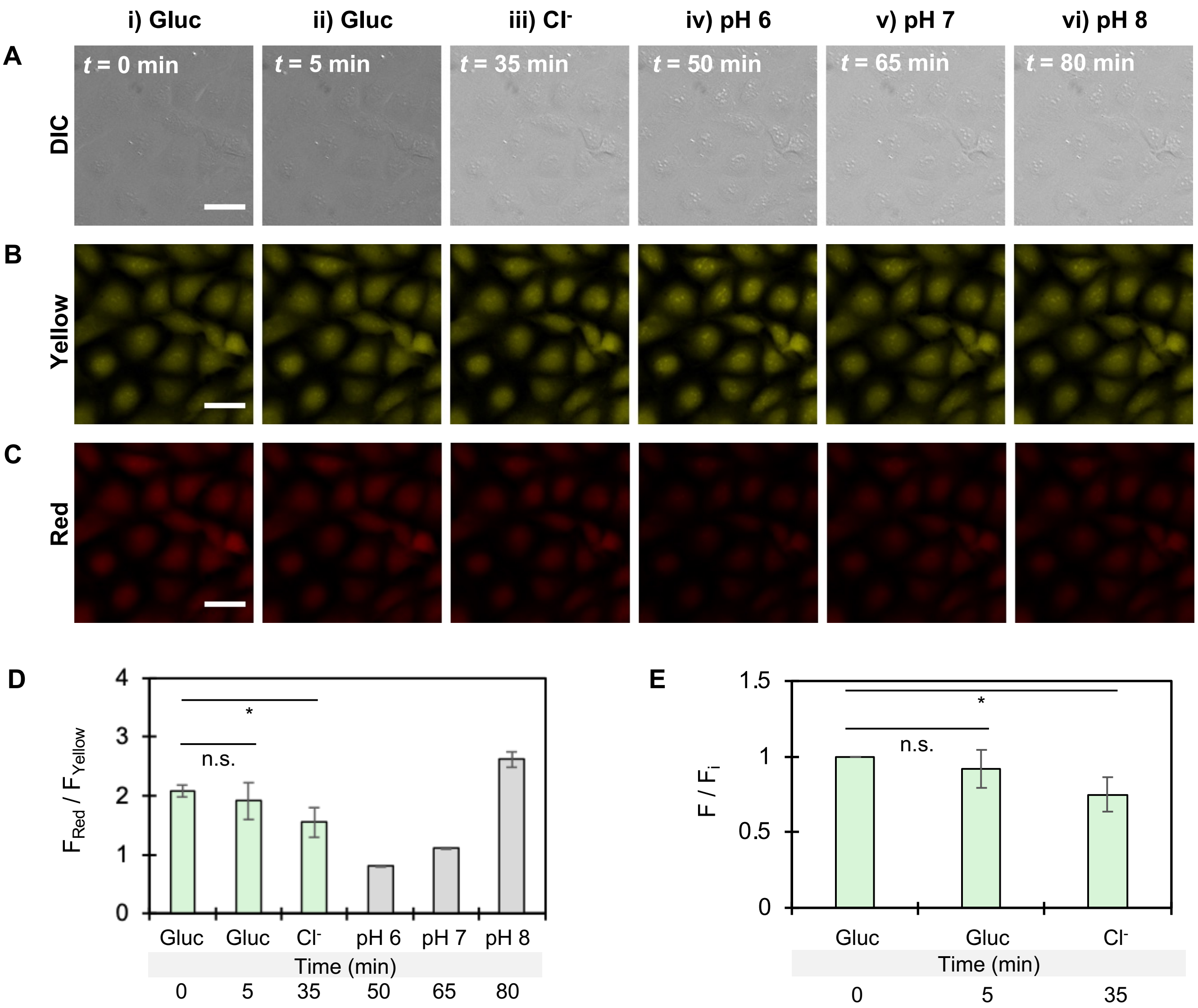


**Figure S31.** Representative (A) DIC and fluorescence images of U-2 OS cells stained with SNARF-1-AM ester in buffer. Fluorescence images are shown for the SNARF-1 emission at (B) 585 nm ($F_{Yellow}$) and (C) 640 nm ($F_{Red}$) in the presence of 137 mM NaGluc (Gluc) buffer at (i) *t* = 0 min ($F_i$) and (ii) *t* = 5 min, (iii) 68.5 mM Gluc/68.5 mM NaCl ($Cl^-$) buffer at *t* = 35 min, and 70 mM Gluc/70 mM $Cl^-$ clamping buffer containing 5 μM nigericin and 5 μM valinomycin at (iv) pH 6 at *t* = 50 min, (v) pH 7 at *t* = 65 min, and (vi) pH 8 at *t* = 80 min. Excitation was provided at 500 nm for both emission wavelengths. Scale bar = 30 μm. Bar graphs of the SNARF-1 (D) ratiometric emissions ($F_{Red}$ /$F_{Yellow}$) and (E) emission responses ($F/F_i$) of $F_{Red}$ /$F_{Yellow}$ for each condition listed. Statistical significances were determined using the average ratiometric emissions ($F_{Red}$ /$F_{Yellow}$) and emission responses ($F/F_i$) at *t* = 5 min and *t* = 35 min for three biological replicates (*N* = 3) relative to $F_i$ (* = *p*-value < 0.05; n.s. = not significant, *p*-value > 0.05). Values are reported as the average ± standard deviation of two fields from each biological replicate.

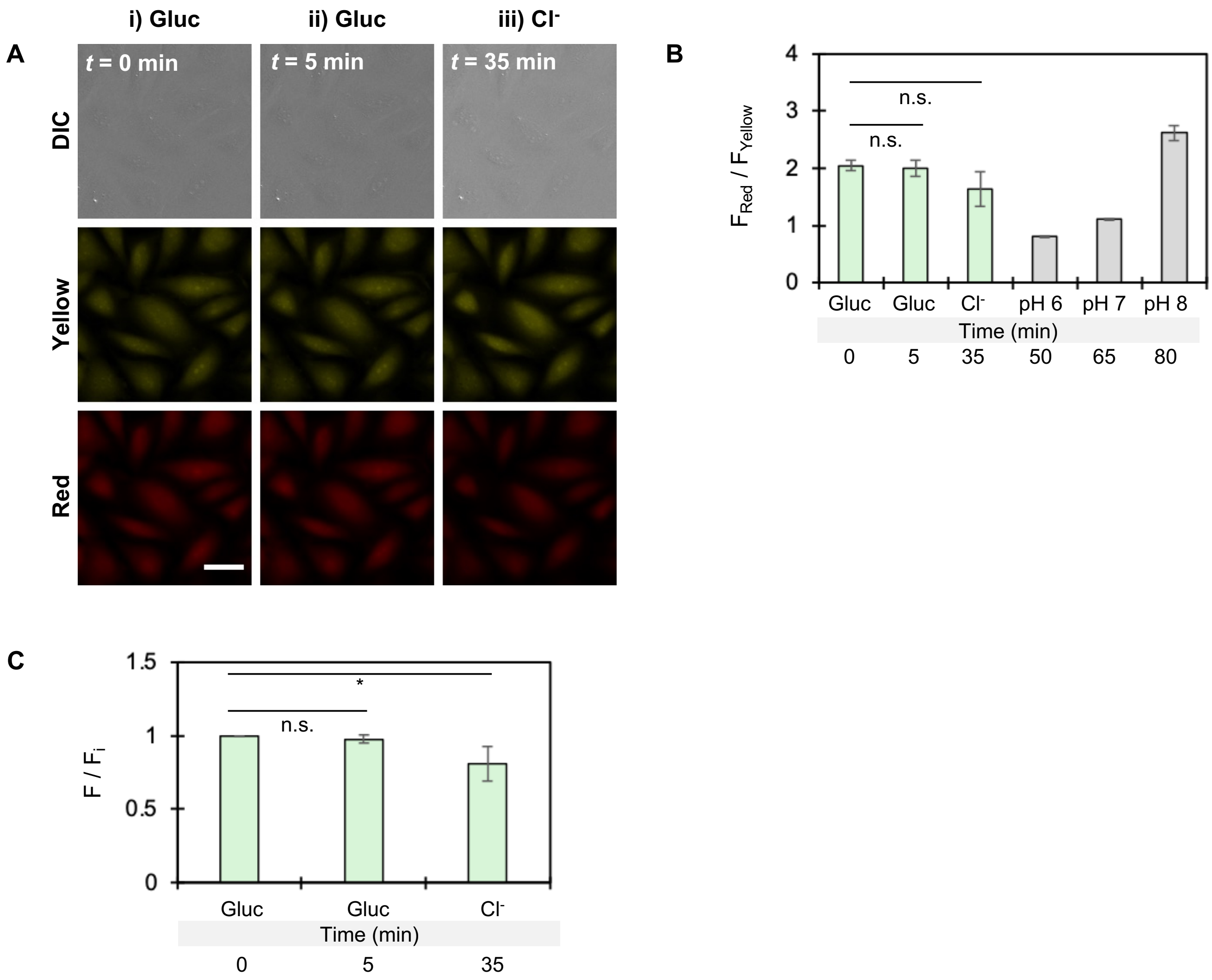


**Figure S32.** (A) Representative DIC (top row) and fluorescence images of U-2 OS cells stained with SNARF-1-AM ester treated with DMSO. Fluorescence images are shown for the SNARF-1 emission at 585 nm ($F_{Yellow}$, middle row) and 640 nm ($F_{Red}$, bottom row) in the presence of 137 mM NaGluc (Gluc) buffer at (i) *t* = 0 min ($F_i$) and (ii) *t* = 5 min (F) followed by perfusion with (iii) 68.5 mM Gluc/68.5 mM NaCl ($Cl^-$) buffer at *t* = 35 min (F). All buffers were supplemented with DMSO. Excitation was provided at 500 nm for both wavelengths. Scale bar = 30 μm. Bar graphs of the SNARF-1 (B) ratiometric emissions ($F_{Red}$ /$F_{Yellow}$) and (C) emission responses ($F/F_i$) of $F_{Red}$ /$F_{Yellow}$ for each condition tested (green bars). The SNARF-1 pH clamping data from Figure S31 (gray bars) are also shown. Statistical significances were determined using the average ratiometric emissions ($F_{Red}$ /$F_{Yellow}$) and emission responses ($F/F_i$) at *t* = 5 min and *t* = 35 min for three biological replicates (*N* = 3) relative to $F_i$ (* = *p*-value < 0.05; n.s. = not significant, *p*-value > 0.05). Values are reported as the average ± standard deviation of two fields from each biological replicate.

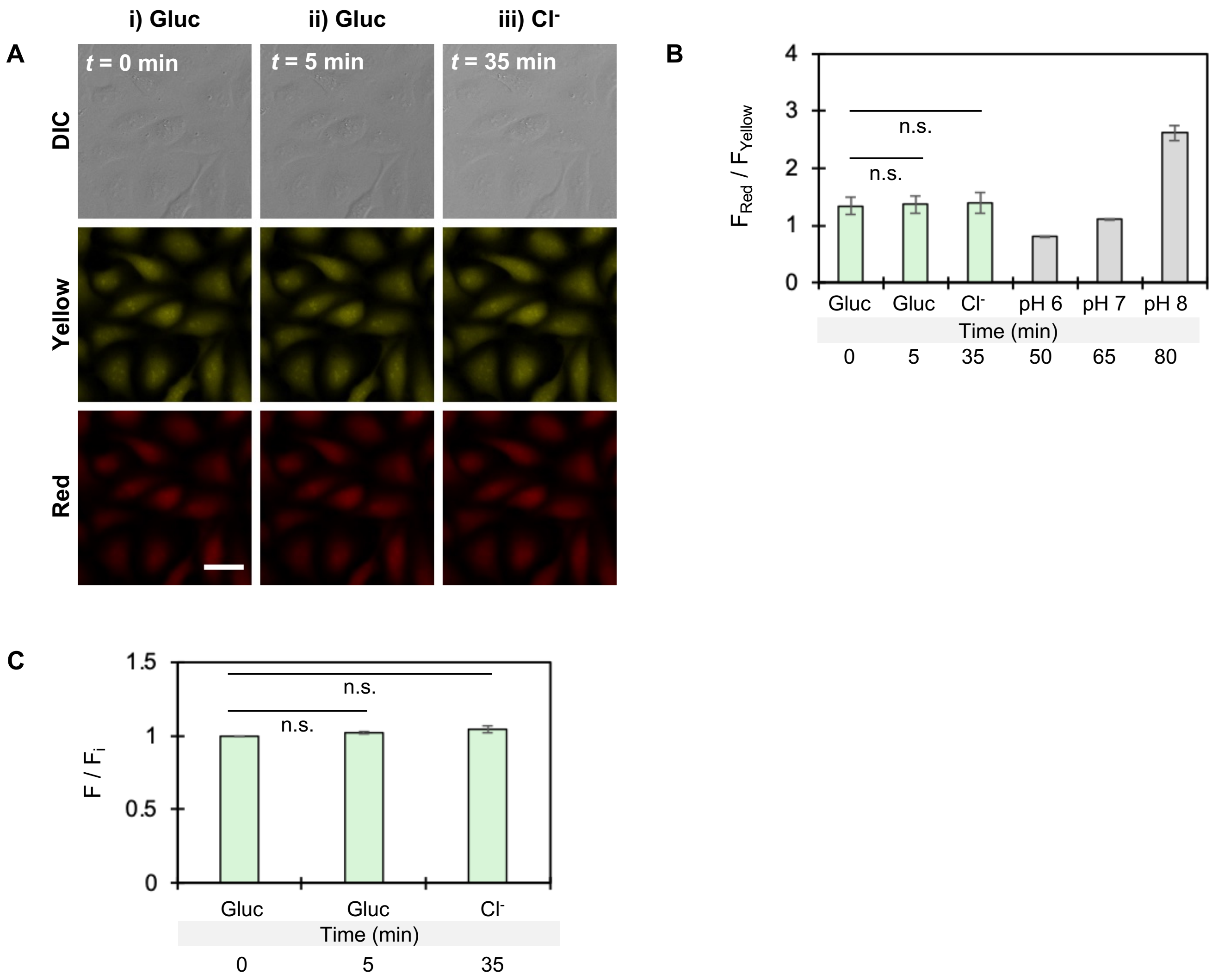


**Figure S33.** (A) Representative DIC (top row) and fluorescence images of U-2 OS cells stained with SNARF-1-AM ester treated with IAA-94. Fluorescence images are shown for the SNARF-1 emission at 585 nm ($F_{Yellow}$, middle row) and 640 nm ($F_{Red}$, bottom row) in the presence of 137 mM NaGluc (Gluc) buffer at (i) $t$ = 0 min ($F_i$) and (ii) $t$ = 5 min (F) followed by perfusion with (iii) 68.5 mM Gluc/68.5 mM NaCl ($Cl^-$) buffer at $t$ = 35 min (F). All buffers were supplemented with 100 μM IAA-94. Excitation was provided at 500 nm for both emission wavelengths. Scale bar = 30 μm. Bar graphs of the SNARF-1 (B) ratiometric emissions ($F_{Red}$ /$F_{Yellow}$) and (C) emission responses ($F/F_i$) of $F_{Red}$ /$F_{Yellow}$ for each condition tested (green bars). The SNARF-1 pH clamping data from Figure S31 (gray bars) are also shown. Statistical significances were determined using the average ratiometric emissions ($F_{Red}$ /$F_{Yellow}$) and emission responses ($F/F_i$) at $t$ = 5 min and $t$ = 35 min for three biological replicates ($N$ = 3) relative to $F_i$ (n.s. = not significant, $p$-value > 0.05). Values are reported as the average ± standard deviation of two fields from each biological replicate.

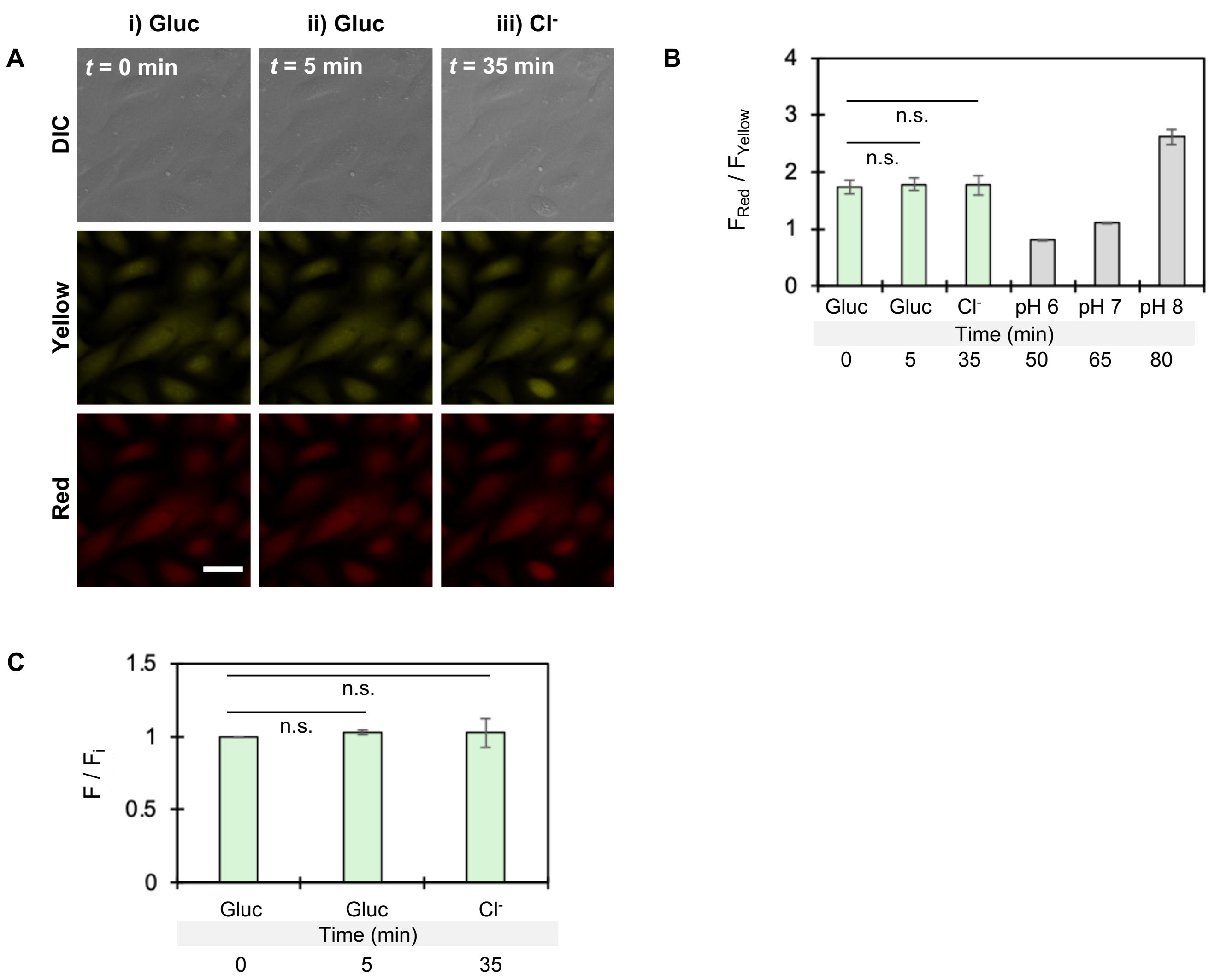


**Figure S34.** (A) Representative DIC (top row) and fluorescence images of U-2 OS cells stained with SNARF-1-AM ester treated with CaCCinh-A01. Fluorescence images are shown for the SNARF-1 emission at 585 nm ($F_{Yellow}$, middle row) and 640 nm ($F_{Red}$, bottom row) in the presence of 137 mM NaGluc (Gluc) buffer at (i) $t$ = 0 min ($F_i$) and (ii) $t$ = 5 min (F) followed by perfusion with (iii) 68.5 mM Gluc/68.5 mM NaCl ($Cl^-$) buffer at $t$ = 35 min (F). All buffers were supplemented with 20 μM CaCCinh-A01. Excitation was provided at 500 nm for both emission wavelengths. Scale bar = 30 μm. Bar graphs of the SNARF-1 (B) ratiometric emissions ($F_{Red}$ /$F_{Yellow}$) and (C) emission responses ($F/F_i$) of $F_{Red}$ /$F_{Yellow}$ for each condition tested (green bars). The SNARF-1 pH clamping data from Figure S31 (gray bars) are also shown. Statistical significances were determined using the average ratiometric emissions ($F_{Red}$ /$F_{Yellow}$) and emission responses ($F/F_i$) at $t$ = 5 min and $t$ = 35 min for three biological replicates ($N$ = 3) relative to $F_i$ (n.s. = not significant, $p$-value > 0.05). Values are reported as the average ± standard deviation of two fields from each biological replicate.

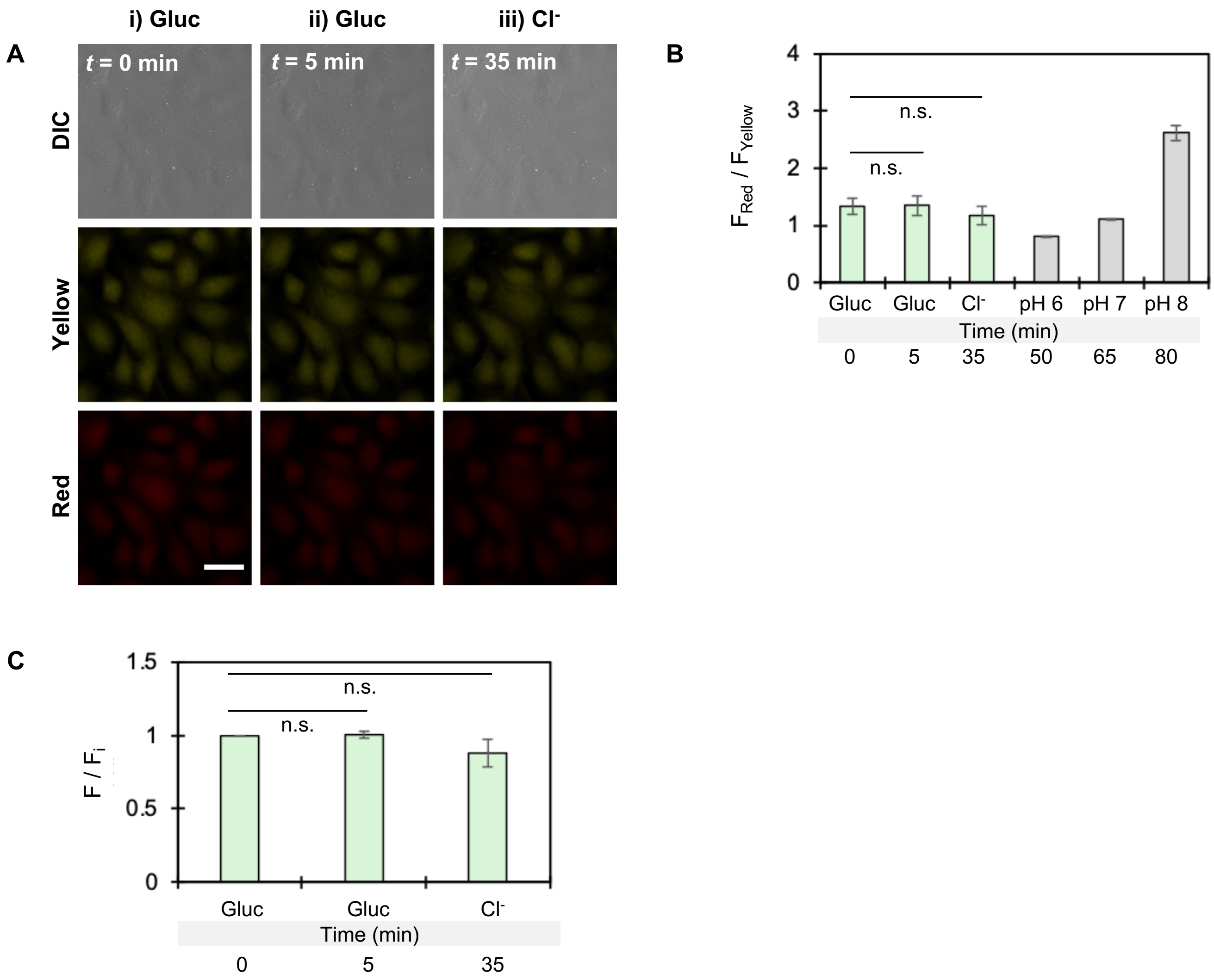


**Figure S35.** (A) Representative DIC (top row) and fluorescence images of U-2 OS cells stained with SNARF-1-AM ester treated with BAPTA-AM ester. Fluorescence images are shown for the SNARF-1 emission at 585 nm ($F_{Yellow}$, middle row) and 640 nm ($F_{Red}$, bottom row) in the presence of 137 mM NaGluc (Gluc) buffer at (i) $t$ = 0 min ($F_i$) and (ii) $t$ = 5 min (F) followed by perfusion with (iii) 68.5 mM Gluc/68.5 mM NaCl ($Cl^-$) buffer at $t$ = 35 min (F). All buffers were supplemented with 20 µM BAPTA-AM ester. Excitation was provided at 500 nm for both emission wavelengths. Scale bar = 30 µm. Bar graphs of the SNARF-1 (B) ratiometric emissions ($F_{Red}$ /$F_{Yellow}$) and (C) emission responses (F/$F_i$) of $F_{Red}$ /$F_{Yellow}$ for each condition tested (green bars). The SNARF-1 pH clamping data from Figure S31 (gray bars) are also shown. Statistical significances were determined using the average ratiometric emissions ($F_{Red}$ /$F_{Yellow}$) and emission responses (F/$F_i$) at $t$ = 5 min and $t$ = 35 for three biological replicates ($N$ = 3) relative to $F_i$ (n.s. = not significant, $p$-value > 0.05). Values are reported as the average ± standard deviation of two fields from each biological replicate.

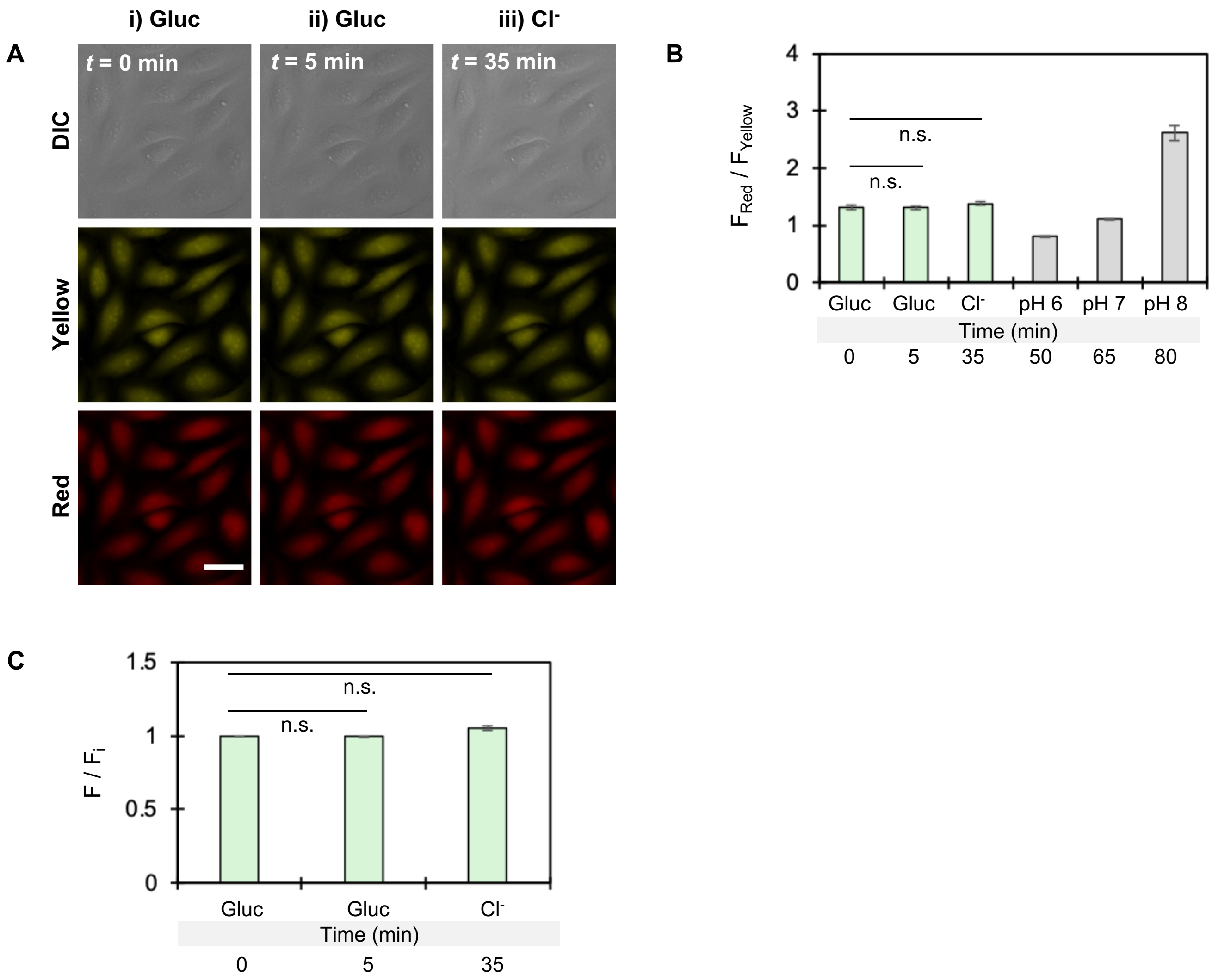


**Figure S36.** (A) Representative DIC (top row) and fluorescence images of U-2 OS cells stained with SNARF-1-AM ester treated with niclosamide. Fluorescence images are shown for the SNARF-1 emission at 585 nm ($F_{Yellow}$, middle row) and 640 nm ($F_{Red}$, bottom row) in the presence of 137 mM NaGluc (Gluc) buffer at (i) $t$ = 0 min ($F_i$) and (ii) $t$ = 5 min (F) followed by perfusion with (iii) 68.5 mM Gluc/68.5 mM NaCl ($Cl^-$) buffer at $t$ = 35 min (F). All buffers were supplemented with 5 µM niclosamide. Excitation was provided at 500 nm for both emission wavelengths. Scale bar = 30 µm. Bar graphs of the SNARF-1 (B) ratiometric emissions ($F_{Red}$ /$F_{Yellow}$) and (C) emission responses ($F/F_i$) of $F_{Red}$ /$F_{Yellow}$ for each condition tested (green bars). The SNARF-1 pH clamping data from Figure S31 (gray bars) are also shown. Statistical significances were determined using the average ratiometric emissions ($F_{Red}$ /$F_{Yellow}$) and emission responses ($F/F_i$) at $t$ = 5 min and $t$ = 35 min for three biological replicates ($N$ = 3) relative to $F_i$ (n.s. = not significant, $p$-value > 0.05). Values are reported as the average ± standard deviation of two fields from each biological replicate.

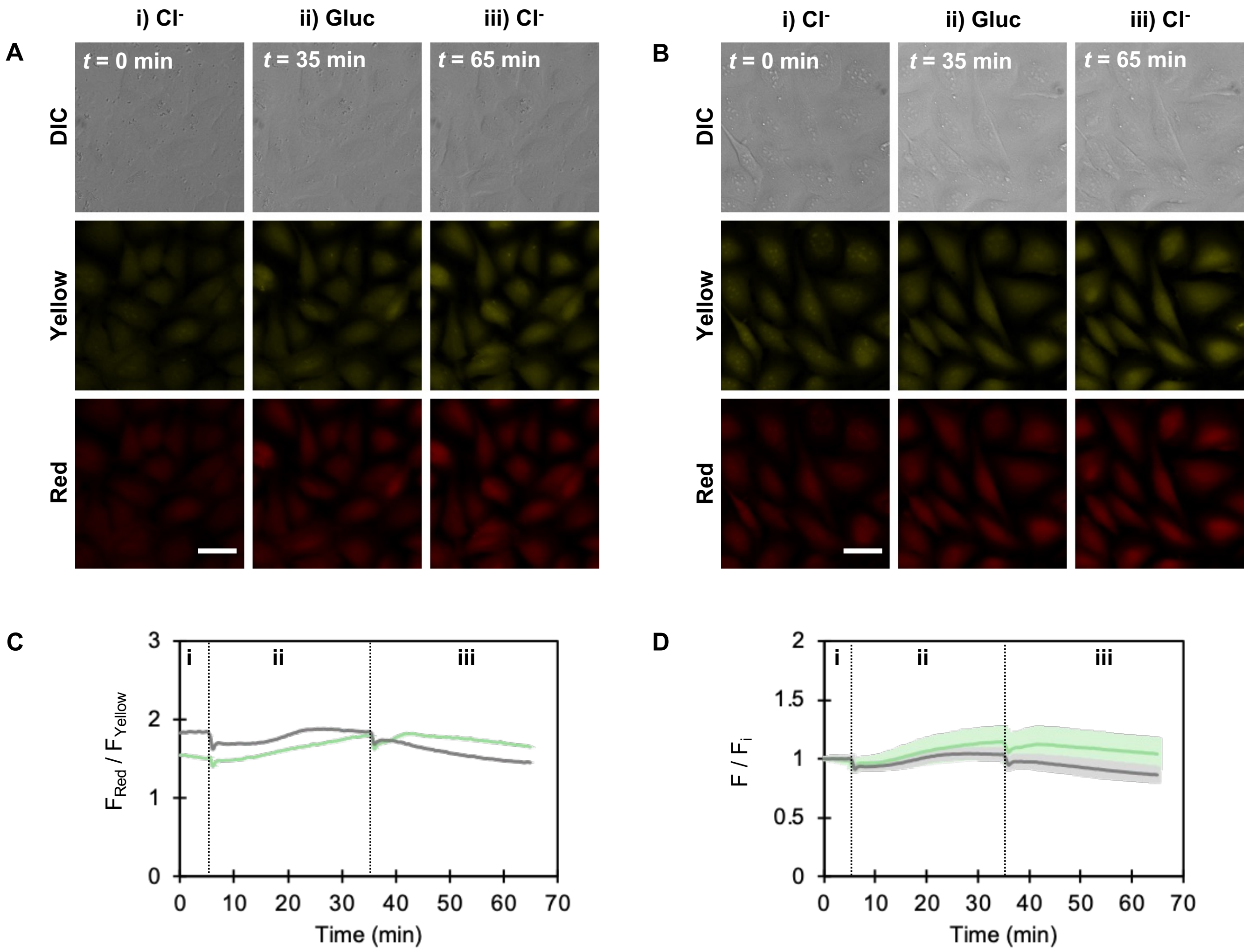


**Figure S37.** Representative DIC (top row) and fluorescence images of U-2 OS cells stained with SNARF-1-AM ester treated with (A) 0.1% DMSO and (B) 10 µM $E_{act}$. Fluorescence images are shown for the SNARF-1 emission at 585 nm ($F_{Yellow}$, middle row) and 640 nm ($F_{Red}$, bottom row) in the presence of (i) 137 mM NaCl ($Cl^-$) buffer at $t$ = 0 min ($F_i$) followed by perfusion with (ii) 137 mM NaGluc (Gluc) buffer at $t$ = 35 min and (iii) 137 mM NaCl ($Cl^-$) at $t$ = 65 min. All buffers were supplemented with 0.1% DMSO or 10 µM $E_{act}$. Excitation was provided at 500 nm for both emission wavelengths. Scale bar = 30 µm. Time course of the SNARF-1 (C) ratiometric emission ($F_{Red}/F_{Yellow}$) and (D) fluorescence response at each time point (F) relative to $t$ = 0 min ($F_i$) with 0.1% DMSO (gray trace) or 10 µM $E_{act}$ (green trace). The first dashed line represents the start of the Gluc buffer perfusion, and the second dashed line represents the start of the $Cl^-$ buffer perfusion. Values are reported as the average ± standard deviation of two fields from three biological replicates.

```
run("Concatenate...", "all_open open");
run("Split Channels");
selectWindow("green");
run("StackReg", "transformation=Translation");
setOption("ScaleConversions", false);
run("Subtract Background...", "rolling=75 stack");
run("Despeckle", "stack");
run("Remove Outliers...", "radius=3 threshold=500 which=Bright");
run("Z Project...", "projection=[Max Intensity]");
run("Duplicate...", "title=G-threshold");
run("Threshold...");
setThreshold(15, 6000);
setThreshold(100, 6000); //for the chloride efflux assay
run("Create Selection");
run("Create Mask");
run("Watershed");
run("Analyze Particles...", "size=10-Infinity exclude clear add");
waitForUser("Check ROIs");
selectWindow("green");
roiManager("multi measure");
```

**Figure S38.** ImageJ macro used to analyze all ChlorON-1-PRO fluorescence microscopy images.

```
run("Concatenate...", "all_open open");
run("Split Channels");
selectWindow("red");
run("StackReg ", "transformation=Translation");
run("Subtract Background...", "rolling=75 stack");
selectWindow("yellow");
run("StackReg ", "transformation=Translation");
run("Subtract Background...", "rolling=75 stack");
run("Z Project...", "projection=[Max Intensity]");
run("Duplicate...", "title=Y-threshold");
run("Threshold...");
setThreshold(200, 6000);
run("Create Selection");
run("Create Mask");
run("Watershed");
run("Analyze Particles...", "size=10-Infinity exclude clear add");
waitForUser("Check ROIs");
selectWindow("yellow");
roiManager("multi measure");
selectWindow("red");
roiManager("multi measure");
```

**Figure S39.** ImageJ macro used to analyze all SNARF-1 fluorescence microscopy images.

### References


(1) The PyMOL Molecular Graphics System, Version 3.1 Schrödinger, LLC.

(2) Webb, B.; Sali, A. Comparative protein structure modeling using MODELLER. *Curr. Protoc. Bioinformatics* **2016**, *54*, 5.6.1–5.6.37. DOI: 10.1002/cpbi.3.

(3) Tutol, J. N.; Ong, W. S. Y.; Phelps, S. M.; Peng, W.; Goenawan, H.; Dodani, S. C. Engineering the ChlorON series: Turn-on fluorescent protein sensors for imaging labile chloride in living cells. *ACS Cent. Sci.* **2024**, *10*, 77–86. DOI: 10.1021/acscentsci.3c01088.

(4) Kille, S.; Acevedo-Rocha, C. G.; Parra, L. P.; Zhang, Z.-G.; Opperman, D. J.; Reetz, M. T.; Acevedo, J. P. Reducing codon redundancy and screening effort of combinatorial protein libraries created by saturation mutagenesis. *ACS Synth. Biol.* **2013**, *2*, 83–92. DOI: 10.1021/sb300037w.

(5) Shen, J.; Snook, R. D. Thermal lens measurement of absolute quantum yields using quenched fluorescent samples as references. *Chem. Phys. Lett.* **1989**, *155*, 583–586. DOI: 10.1016/0009-2614(89)87477-9.

(6) Chen, C.; Pathiranage, V.; Ong, W. S. Y.; Dodani, S. C.; Walker, A. R.; Fang, C. A twisted chromophore powers a turn-on fluorescent protein chloride sensor. *Proc. Natl. Acad. Sci. U. S. A.* **2025**, *122*, e2508094122. DOI: 10.1073/pnas.2508094122.

(7) Meng, E. C.; Goddard, T. D.; Pettersen, E. F.; Couch, G. S.; Pearson, Z. J.; Morris, J. H.; Ferrin, T. E. UCSF ChimeraX: tools for structure building and analysis. *Protein Sci.* **2023**, *32*, e4792. DOI: 10.1002/pro.4792.

(8) Pettersen, E. F.; Goddard, T. D.; Huang, C. C.; Meng, E. C.; Couch, G. S.; Croll, T. I.; Morris, J. H.; Ferrin, T. E. UCSF ChimeraX: structure visualization for researchers, educators, and developers. *Protein Sci.* **2021**, *30*, 70–82. DOI: 10.1002/pro.3943.

(9) Chen, V. B.; Arendall, W. B.; Headd, J. J.; Keedy, D. A.; Immormino, R. M.; Kapral, G. J.; Murray, L. W.; Richardson, J. S.; Richardson, D. C. MolProbity: all-atom structure validation for macromolecular crystallography. *Acta Crystallogr. D Biol. Crystallogr.* **2010**, *66*, 12–21. DOI: 10.1107/s0907444909042073.

(10) Jurrus, E.; Engel, D.; Star, K.; Monson, K.; Brandi, J.; Felberg, L. E.; Brookes, D. H.; Wilson, L.; Chen, J.; Liles, K.; et al. Improvements to the APBS biomolecular solvation software suite. *Protein Sci.* **2018**, *27*, 112–128. DOI: 10.1002/pro.3280.

(11) Jorgensen, W. L.; Chandrasekhar, J.; Madura, J. D.; Impey, R. W.; Klein, M. L. Comparison of simple potential functions for simulating liquid water. *J. Chem. Phys.* **1983**, *79*, 926–935. DOI: 10.1063/1.445869.

(12) Case, D. A.; Belfon, K.; Ben-Shalom, I. Y.; Brozell, S. R.; Cerutti, D. S.; Cheatham, T. E. I.; Cruzeiro, V. W. D.; Darden, T. A.; Duke, R. E.; Giambasu, G; et al. Amber 2020. University of California: San Francisco, 2020.

(13) Maier, J. A.; Martinez, C.; Kasavajhala, K.; Wickstrom, L.; Hauser, K. E.; Simmerling, C. Ff14SB: improving the accuracy of protein side chain and backbone parameters from ff99sb. *J. Chem. Theory Comput.* **2015**, *11*, 3696–3713. DOI: 10.1021/acs.jctc.5b00255.

(14) Hix, M. A.; Walker, A. R. AutoParams: an automated web-based tool to generate force field parameters for molecular dynamics simulations. *J. Chem. Inf. Model.* **2023**, *63*, 6293–6301. DOI: 10.1021/acs.jcim.3c01049.

(15) Lee, T. S.; Cerutti, D. S.; Mermelstein, D.; Lin, C.; Legrand, S.; Giese, T. J.; Roitberg, A.; Case, D. A.; Walker, R. C.; York, D. M. GPU-accelerated molecular dynamics and free energy methods in

Amber18: performance enhancements and new features. *J. Chem. Inf. Model.* **2018**, *58*, 2043–2050. DOI: 10.1021/acs.jcim.8b00462.
(16) Loncharich, R. J.; Brooks, B. R.; Pastor, R. W. Langevin dynamics of peptides: the frictional dependence of isomerization rates of N-acetylalanyl-N'-methylamide. *Biopolymers* **1992**, *32*, 523–535. DOI: 10.1002/bip.360320508.
(17) Miyamoto, S.; Kollman, P. A. Settle: an analytical version of the shake and rattle algorithm for rigid water models. *J. Comput. Chem.* **1992**, *13*, 952–962. DOI: 10.1002/jcc.540130805.
(18) Darden, T.; York, D.; Pedersen, L. Particle mesh Ewald: an N·log(N) method for Ewald sums in large systems. *J. Chem. Phys.* **1993**, *98*, 10089–10092. DOI: 10.1063/1.464397.
(19) Roe, D. R.; Cheatham, T. E. PTRAJ and CPPTRAJ: software for processing and analysis of molecular dynamics trajectory data. *J. Chem. Theory Comput.* **2013**, *9*, 3084–3095. DOI: 10.1021/ct400341p.
(20) Durrant, J. D.; Votapka, L.; Sørensen, J.; Amaro, R. E. POVME 2.0: an enhanced tool for determining pocket shape and volume characteristics. *J. Chem. Theory Comput.* **2014**, *10*, 5047–5056. DOI: 10.1021/ct500381c.
(21) Virtanen, P.; Gommers, R.; Oliphant, T. E.; Haberland, M.; Reddy, T.; Cournapeau, D.; Burovski, E.; Peterson, P.; Weckesser, W.; Bright, J.; van der Walt, S. J.; et al. SciPy 1.0: fundamental algorithms for scientific computing in python. *Nat. Methods* **2020**, *17*, 261–272. DOI: 10.1038/s41592-019-0686-2.
(22) Cock, P. J. A.; Antao, T.; Chang, J. T.; Chapman, B. A.; Cox, C. J.; Dalke, A.; Friedberg, I.; Hamelryck, T.; Kauff, F.; Wilczynski, B.; de Hoon, M. J. L. Biopython: freely available python tools for computational molecular biology and bioinformatics. *Bioinformatics* **2009**, *25*, 1422–1423. DOI: 10.1093/bioinformatics/btp163.
(23) Peng, W.; Tutol, J. N.; Phelps, S. M.; Kam, H.; Lynd, J. K.; Dodani, S. C. Directed evolution of a genetically encoded indicator for chloride. *ACS Synth. Biol.* **2025**, *14*, 1009–1013. DOI: 10.1021/acssynbio.4c00818.
(24) Seksek, O.; Henry-Toulmé, N.; Sureau, F.; Bolard, J. SNARF-1 as an intracellular pH indicator in laser microspectrofluorometry: a critical assessment. *Anal. Biochem.* **1991**, *193*, 49–54. DOI: 10.1016/0003-2697(91)90042-r.
(25) Schindelin, J.; Arganda-Carreras, I.; Frise, E.; Kaynig, V.; Longair, M.; Pietzsch, T.; Preibisch, S.; Rueden, C.; Saalfeld, S.; Schmid, B.; et al. Fiji: an open-source platform for biological-image analysis. *Nat Methods* **2012**, *9*, 676–682. DOI: 10.1038/nmeth.2019.
(26) Statistical significances were calculated using the GraphPad QuickCalcs *t*-test Calculator. https://www.graphpad.com/quickcalcs/ttest1/ (accessed 2025-07-09).
